# Multimodal-Multisensory Experiments: Design and Implementation

**DOI:** 10.1101/2020.12.01.405795

**Authors:** Moein Razavi, Takashi Yamauchi, Vahid Janfaza, Anton Leontyev, Shanle Longmire-Monford, Joseph Orr

## Abstract

The human mind is multimodal. Yet most behavioral studies rely on century-old measures of behavior - task accuracy and latency (response time). Multimodal and multisensory analysis of human behavior creates a better understanding of how the mind works. The problem is that designing and implementing these experiments is technically complex and costly. This paper introduces versatile and economical means of developing multimodal-multisensory human experiments. We provide an experimental design framework that automatically integrates and synchronizes measures including electroencephalogram (EEG), galvanic skin response (GSR), eye-tracking, virtual reality (**VR**), body movement, mouse/cursor motion and response time. Unlike proprietary systems (e.g., iMotions), our system is free and open-source; it integrates **PsychoPy, Unity** and Lab Streaming Layer (**LSL**). The system embeds LSL inside **PsychoPy/Unity** for the synchronization of multiple sensory signals - gaze motion, electroencephalogram (EEG), galvanic skin response (GSR), mouse/cursor movement, and body motion - with low-cost consumer-grade devices in a simple behavioral task designed by **PsychoPy** and a virtual reality environment designed by **Unity**. This tutorial shows a step-by-step process by which a complex multimodal-multisensory experiment can be designed and implemented in a few hours. When conducting the experiment, all of the data synchronization and recoding of the data to disk will be done automatically.

## 1. Introduction

### 1.1. The mind is multimodal

Psychology is the scientific study of the brain, mind and behavior. The brain supervises different autonomic functions such as cardiac activity, respiration, perspiration, etc. Current methods to study human behavior include self-report, observation, task performance, gaze, gait and body motion, and physiological measures such as electroencephalogram (EEG), electrocardiogram (ECG), functional magnetic resonance imaging (fMRI). These measures are indicators of human behavior; however, the gap here is that they are mostly studied separately from each other, while human behavior is inherently multimodal and multisensory, where different measures are connected and dependent on each other. Signals from multiple sources have overlap in locations (spatial) or timing (temporal) in the brain; hence, a single measurement cannot be as informative as multiple measurements in distinguishing between different functions [1].

A more accurate behavioral measurement needs different measures to be recorded and analyzed concurrently [2]. Hochenberger (2015) suggests that multimodal experiments facilitate the perception of senses that operate in parallel [3]. Similar to using multiple classifiers to improve accuracy in a classification task, combining different measures in studies of brain functionality and its association with behavior helps improve the predictions of human behavior [4].

Designing and developing a multimodal and multisensory study is complicated and costly. All events and time series must be recorded and timestamped together; data from multiple measurements and multiple subjects should be stored in an easily accessible and analyzable format. All the devices must start recording at the same time without the effort of running the devices one by one. For psychological and cognitive science experiments, an instant and easy to setup system is vital, due to the need to collect data from a large group of participants (around 50-60 participants).

There are several proprietary systems that ease multimodal-multisensory experimentation (e.g., iMotions). Yet, it is costly and challenging to integrate different devices with these systems. Due to limitations of proprietary software the stimulus presentation and data acquisition should be in different software which makes it difficult for 1) synchronization (since different software may have different processing times which results in different delays) and 2) instant and easy experiment setup. Here we present a system that allows stimulus presentation, data acquisition and recording in the same [opensource] software and how to make this process automatic and adaptable for various multimodal-multisensory experiments.

#### Contributions

To summarize, this paper makes the following contributions.

- For psychological and cognitive science experiments, an instant and easy-to-setup system that can be used for multimodal-multisensory experiments is vital. We have designed a comprehensive, customizable and fully opensource system in **PsychoPy** and **Unity** platforms which can be used for that purpose. To the best of our knowledge, our system is the first of this kind.
- We have provided a tutorial on how to make stimulus presentation and data acquisition in the same [opensource] software and how to make the process of synchronizing multiple sensors, devices and markers (from environment and the user response) and saving the data on disk automatically.
- We have created several applications and also customized the SDKs of different devices to create multimodal-multisensory experiments using LSL and made the source files open access. By studying the source files, users will find how to customize their devices’ SDKs for this purpose.
- Due to limitation of the proprietary systems for supporting different devices, we have provided a platform which makes it possible for all opensource devices to be synchronized together automatically. Also, with the aid of our system, all non-opensource devices which can send data to one of the opensource platforms such as C, C++, C#, Python, Java and Octave can be synchronized.

In what follows, section 1.2 reviews previous works and their findings that elucidate the importance of using multisensory experiments. Section 2 discusses major challenges of implementing multisensory experiments and the available methods of addressing those challenges. Section 3 presents an overview of the tools and methods utilized in the proposed system. Sections 4 and 5, provide a step-by-step tutorial on developing multimodal/multisensory experiments in **PsychoPy** and **Unity** platforms, respectively. Finally, section 6 presents the results and discusses the cases of use and potential future works.

### 1.2. Related Work

Although many past studies provided useful information about human behavior using only a single source, multiple sources are involved with actual behavior in the natural environment [5]. A few studies tried to integrate multiple measures into the experimental psychology/neuroscience portal. Reeves et al. (2007) state the importance of using multiple measures in the goals of Augmented Cognition (AUGCOG) [6]; they discuss the combination of multiple measures together as a factor for the technologies that improve Cognitive State Assessment (CSA). Jimenez-Molina et al. (2018) showed that analyzing all measures of electrodermal activity (EDA), photoplethysmogram (PPG), EEG, temperature and pupil dilation at the same time, significantly improves the classification accuracy in a web-browsing workload classification task, compared to using a single measure or a combination of some of them [7].

Several studies to date have put these recommendations to work by integrating several measures concurrently. Born et al. (2019) used EEG, GSR and eye-tracking to predict task performance in a task load experiment; they found that low-beta frequency bands, pupil dilations and phasic components of GSR were correlated with task difficulty [8]. They also showed that the statistical results of analyzing EEG and GSR together were more reliable than analyzing them individually. Leontyev et al. combined user response time and mouse movement features with machine learning technics and found an improvement in the accuracy of predicting attention-deficit/hyperactivity disorder (ADHD) [9–11]. Yamauchi et al. combined behavioral measures and multiple mouse motion features to better predict people’s emotions and cognitive conflict in computer tasks [12,13]. Yamauchi et al. further demonstrated that people’s emotional experiences change as their tactile sense (touching a plant) was augmented with visual sense (“seeing” their touch) in a multisensory interface system [14]. Chen et al. (2012) tried to identify possible correlations between increasing levels of cognitive demand and modalities from speech, digital pen, and freehand gesture to eye activity, galvanic skin response, and EEG [15]. Lazzeri et al. (2014) used physiological signals, eye gaze, video and audio acquisition to perform an integrated affective and behavioral analysis in Human-Robot Interaction (HRI) [16]; by acquiring synchronized data from multiple sources, they investigated how autism patients can interact with affective robots. Charles and Nixon (2019) reviewed 58 articles on mental workload tasks. They found that physiological measurements such as ECG, respiration, GSR, blood pressure, EOG and EEG need to be triangulated because though they are sensitive to mental workload, no single measure satisfies to predict mental workload [17]. Lohani et al. (2019) suggest that analyzing multiple measures such as head movement together with physiological measures (e.g., EEG, heart rate, etc.) can be used for the drivers to detect cognitive states (e.g., distraction) [18]. Gibson et al. (2014) integrated questionnaires, qualitative methods, and physiological measures including ECG, respiration, electrodermal activity (EDA) and skin temperature to study activity settings in disabled youth [19]; they stated that using multiple measures reflects a better real-world setting of the youth experiences. Sciarini and Nicholson (2009) used EEG, eye blink, respiration, cardiovascular activity and speech measures in a workload task performance [20]; as they outlined, using only one measure is not sufficient in the multidimensional tasks in a dynamic environment. Thus, multiple measures should be considered. The authors also highlighted the lack of clear guidance on how to integrate different systems as an important issue.

Integrating multiple measurements is quite complicated. Although the aforementioned studies integrated multiple measures, they did not synchronize these measures together. They have different sampling rates and also lack a coherent and easy to implement method for combining the measures. Asynchronized multiple measurs are prone to error and make it difficult to assess their interactions with each other.

The following section provides a summary of the challenges for device integration and different methods of addressing them.

## 2. Multisensory Experiments

### 2. 1. Challenges of Implementation

There are several challenges to be addressed during the integration process:

1. All events and time series must be recorded and timestamped together, with the same timing reference, to be analyzable.
2. Because multiple subjects (N>50) are needed in psychological experiments for statistical analysis, data from multiple measures and multiple subjects should be stored in a format that can be easily analyzed together.
3. A method should be used to record the data from all the devices simultaneously without the need to run the devices one by one.

For the first challenge, UDP (User Datagram Protocol) and TCP (Transmission Control Protocol) offer potential solutions. However, UDP is unreliable since it is a connectionless protocol that does not guarantee data delivery, order, or duplicate protection. On the other hand, TCP is a connection-oriented protocol that guarantees errorless, reliable and ordered data streaming and it works with internet protocol (IP) for data streaming. Thus, for reliable data transport, TCP is preferred.

For the second and third challenges, there are at least two options. One is to use proprietary software (e.g., iMotions, Biopac, etc.) which has its own advantages. For example, it has extensive support and is relatively easy to implement. However, proprietary software is costly and restrictive; it often forces the researchers to purchase other proprietary devices that are functional only for that particular software design. Because of that, the users are limited in the number of experimental manipulations they can introduce. Finally, proprietary software developers often charge steep licensing fees, which limit people’s access (especially in developing countries). Thus, it is imperative to devise a system that will address the problem of simultaneous observation, but that is also accessible to everyone.

Another method, which is more versatile and preferred, is to employ opensource tools that are available for multiple platforms (e.g., LSL). **LSL** (https://github.com/sccn/labstreaminglayer/wiki) is available on almost all open-source platforms including, Python, C, C++, C#, Java, Octave, etc. Currently, the majority of the consumer and research-grade devices support LSL. Above all, as long as these devices are capable of sending their data to the platforms like Python, C, MATLAB, etc., **LSL** can be used for streaming their data. **LSL** uses TCP for stream transport and UDP for stream discovery and time synchronization.

**LSL** is developed by the Swartz Center for Computational Neuroscience (SCCN) at the University of California San Diego (UCSD).

**LSL** can be embedded in free and open-source stimulus presentation platforms like **PsychoPy** (https://www.psychopy.org/) and **Unity** (https://unity.com/, Figure 1).

**Figure 1.**
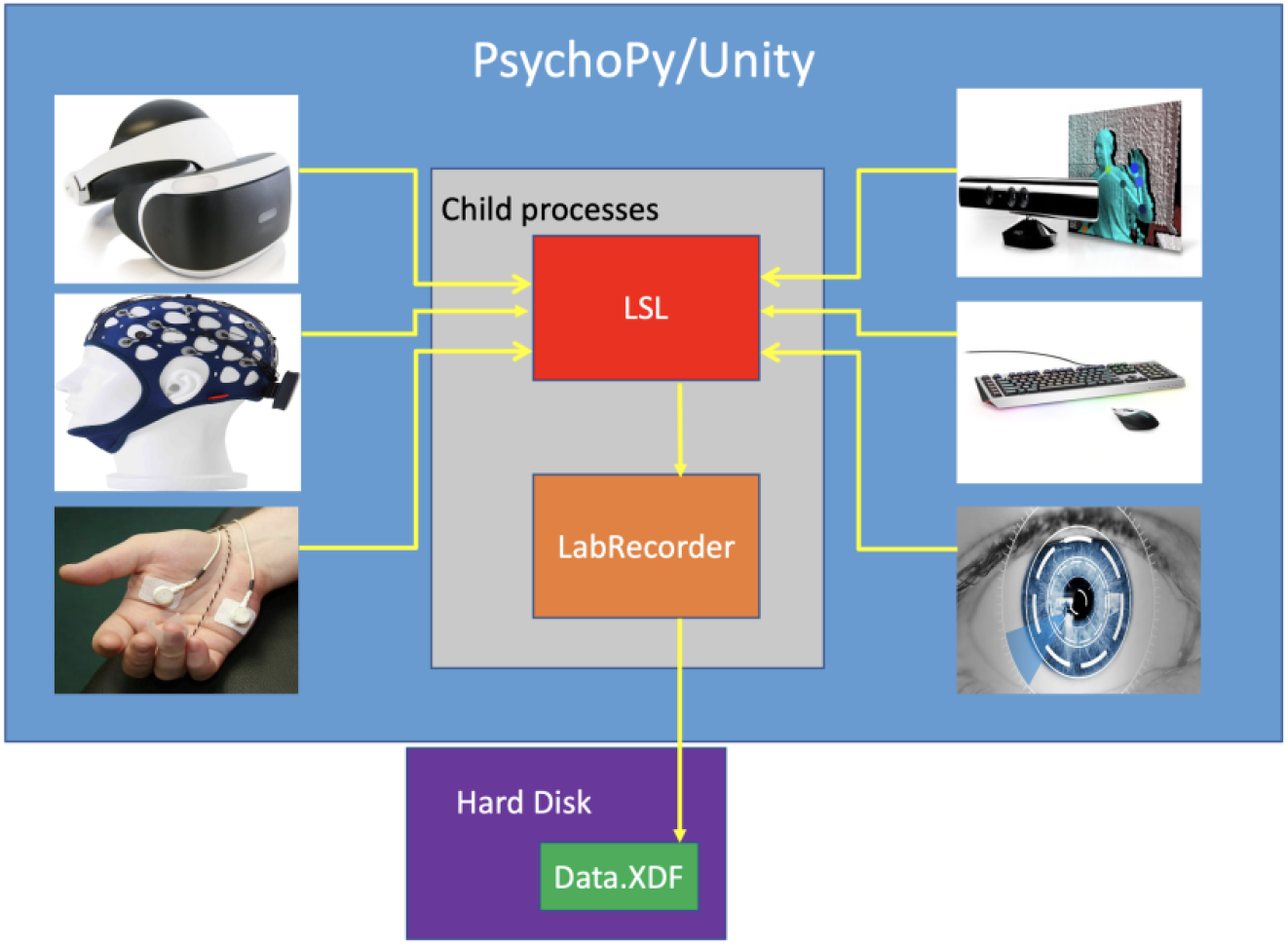
Schematic architecture for integrating devices: Objects that are defined in **PsychoPy/Unity** (usually task markers) can send data directly to **LSL**. Also, different devices can send data to **LSL** by calling child processes inside **PsychoPy/Unity**

Wang et al. (2014) used **PsychoPy**, EEG, and **LSL** for Brain-Computer Interface (BCI) stimulus presentations. They used these to synchronize the stimulus markers and EEG measurements [21].

Figure 2, shows the overall structure of the method proposed in this paper.

**Figure 2.**
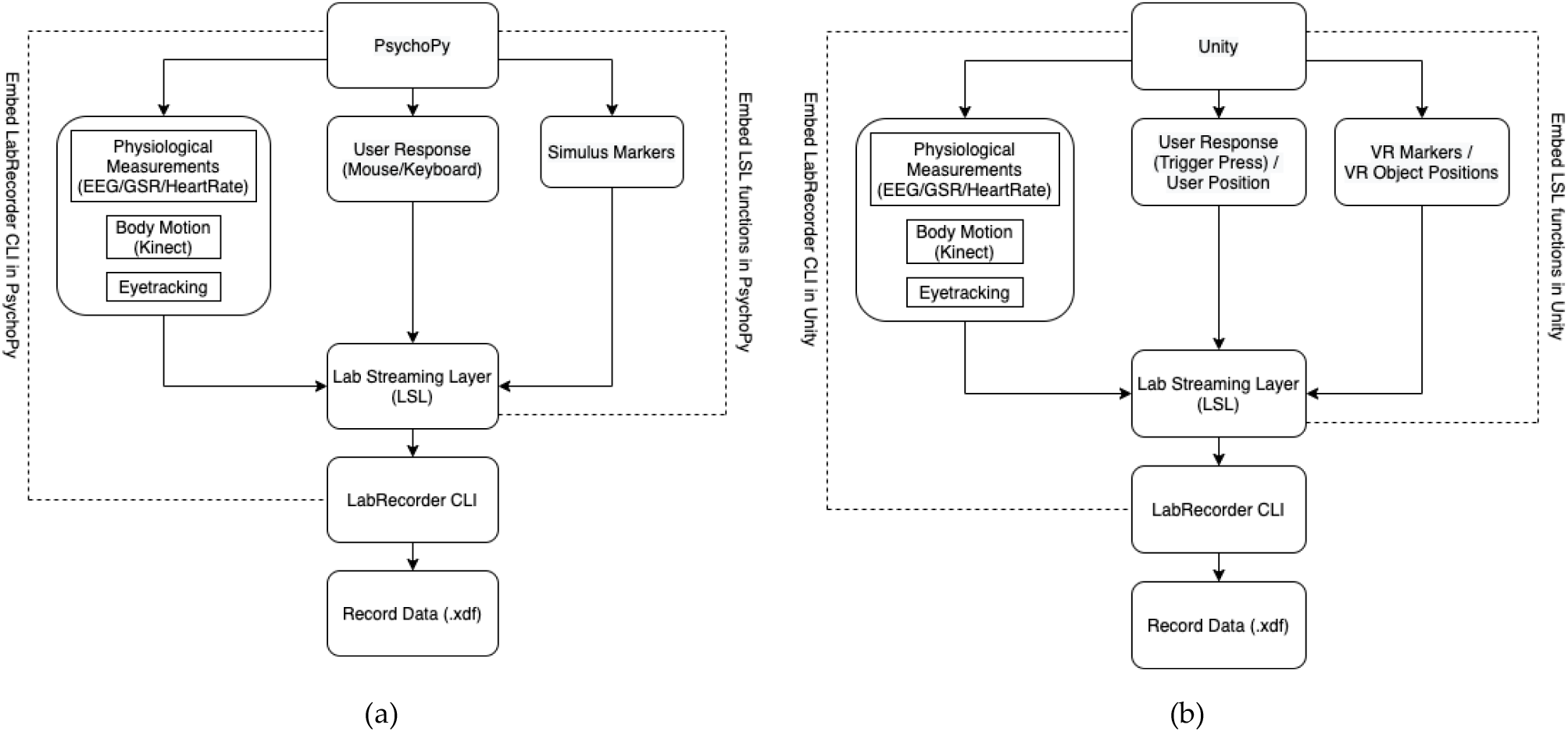
Flowchart of creating an automatic multisensory experiment using (a) **PsychoPy**; (b) **Unity**

As shown in Figure 2, creating a multimodal/multisensory experiment can be done in **PsychoPy/Unity** environment. Different devices send data to **LSL** using **LSL** functions that are embedded in **PsychoPy/Unity**. Then by embedding LabRecorder in **PsychoPy/Unity**, data can be recorded on the disk as a .*xdf* file.

The devices used for this paper can be called and start recording by running child processes in Python and C#, which allows the processes to be directly embedded in **PsychoPy** and **Unity**, respectively. Finally, everything can be started from a main **PsychoPy** experiment or **Unity VR** project.

In the following, an introduction to **PsychoPy/Unity, LSL,** and **LabRecorder** (https://github.com/labstreaminglayer/App-LabRecorder/releases) is provided.

## 3. Overview: PsychoPy, Unity, Lab Streaming Layer (LSL), Lab Recorder

Our overall strategy for building a multimodal and multisensory experiment is to bridge **PsychoPy/Unity, LSL**, and **LabRecorder**. The principle of integration is to call different devices from **PsychoPy/Unity** to send their data streams to **LSL; LabRecorder** starts recording the streams available on **LSL** from **command line**, which would also be embedded in **PsychoPy/Unity**.

**PsychoPy** allows for the building of behavioral experiments with little to no programming experience using premade templates for stimulus presentation and response collection. **Unity** enables the creation of 2D and 3D gaming environments. **LSL** is a set of libraries for synchronous collection of multiple time series in research experiments (https://labstreaminglayer.readthedocs.io/info/intro.html). **LabRecorder** is an **LSL** application that allows saving all the streams that are available on **LSL** on disk in a single .*xdf* file.

### PsychoPy

**PsychoPy** is an open-source software package written in the Python programming language primarily for use in Neuroscience and Experimental Psychology research [22,23] (https://www.psychopy.org/). **PsychoPy** has three main building blocks for constructing behavioral experiments: stimulus/response components, routines and loops (Figure 3).

**Figure 3.**
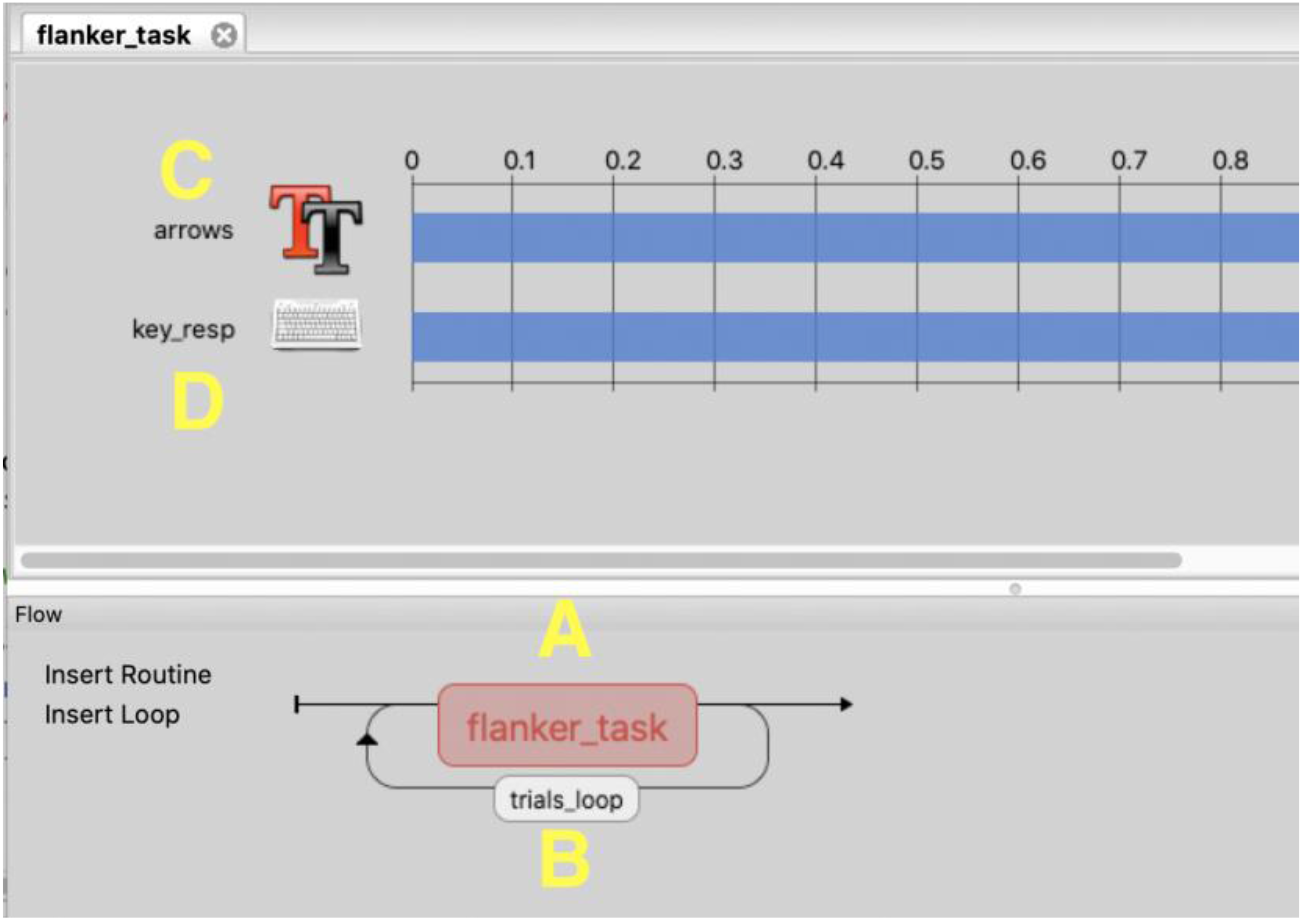
**PsychoPy** components: A) routine B) loop C) text stimulus D) keyboard response

Stimulus components are premade templates for displaying various types of stimuli: geometric shapes, pictures, videos, audio signals, etc. The user can control which stimuli they want to present, in what order and for how long (along with other stimulus-specific properties). Similarly, response components allow for recording different types of responses: key press, mouse clicks/moves, as well as vocal responses.

Stimulus and response components are organized within routines, which are a sequence of events within one experimental trial. For example, consider a Flanker task (Figure 3) in which a participant is presented with five arrows, with the middle arrow pointing to the same direction as surrounding arrows (congruent) or to the opposite direction (incongruent). The participant has to press the key on the keyboard indicating the direction of the middle arrow (“left” or “right” arrow key). In this task, the stimulus component “arrows” (Figure 3C) presents the arrangement of letters for a given trial, while component “key_resp” (Figure 3D) records the participant’s response. Altogether, they constitute the routine “flanker_task” (Figure 3A).

The trial parameters (in this case, which succession of arrows to show, e.g., →→→→→ or ←←→←←) are controlled by the loop “trials_loop” (Figure 3B). Loops contain information about the variables that are supposed to change from trial to trial. As the name suggests, loops repeat routines updating the routine components with the values prescribed by the experimenter.

**PsychoPy** provides an option to include custom python code, which can be embedded in the beginning or the end of the experiment, in the beginning or the end of each routine, and for each frame of the screen; the code written in Each Frame will run every refreshment cycle of the monitor screen (Figure 4).

**Figure 4.**
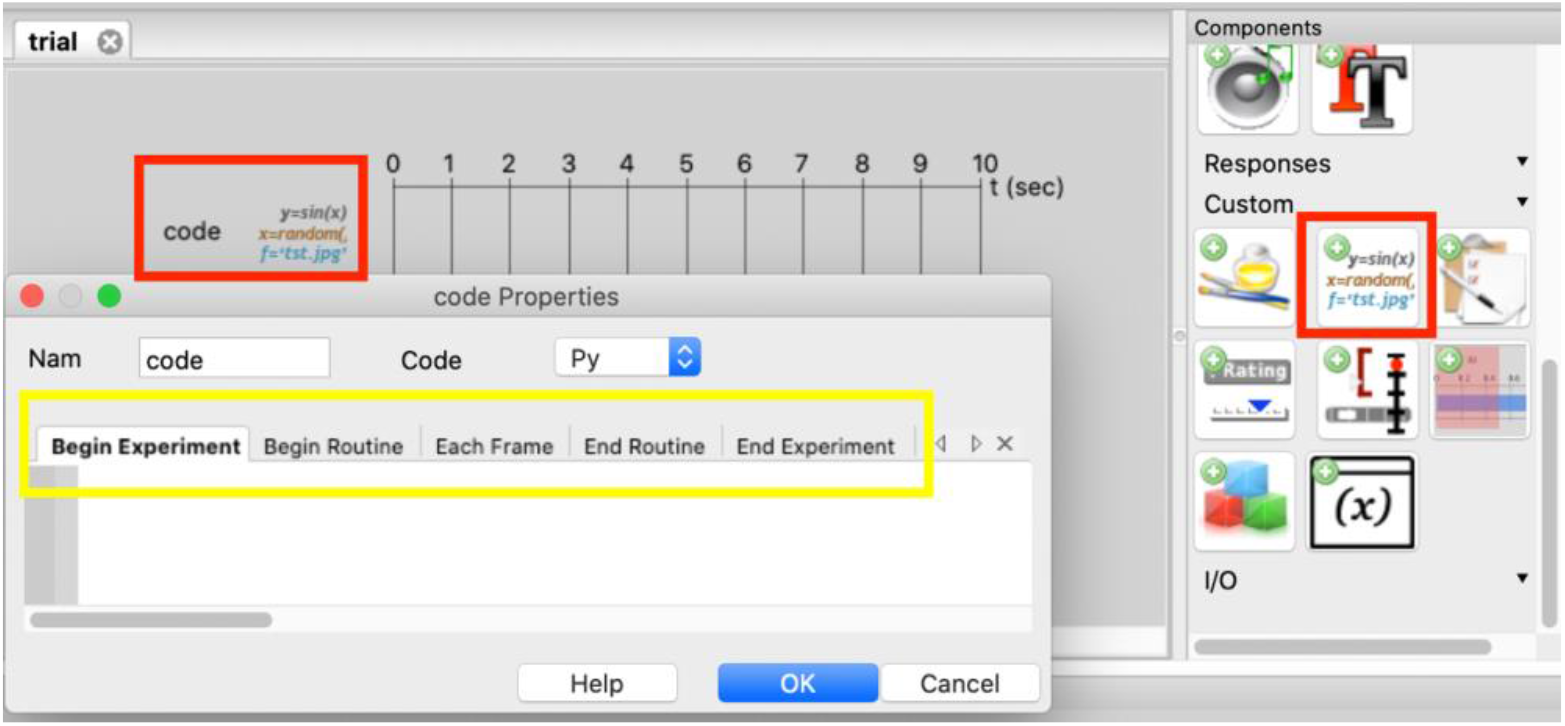
**PsychoPy** python custom code component

### Unity

**Unity** is a game engine platform that allows creating games in 2D and 3D environments. It enables the users to script in C# for handling the scenes and objects. In **Unity**, the game environment is called a **Scene**, and each component in the environment (e.g., characters) is called a **GameObject**. Every **GameObject** will be defined in a **Scene**. Complete documentation of **Unity** is available on https://docs.unity3d.com/Manual/UsingTheEditor.html (Figure 5).

**Figure 5.**
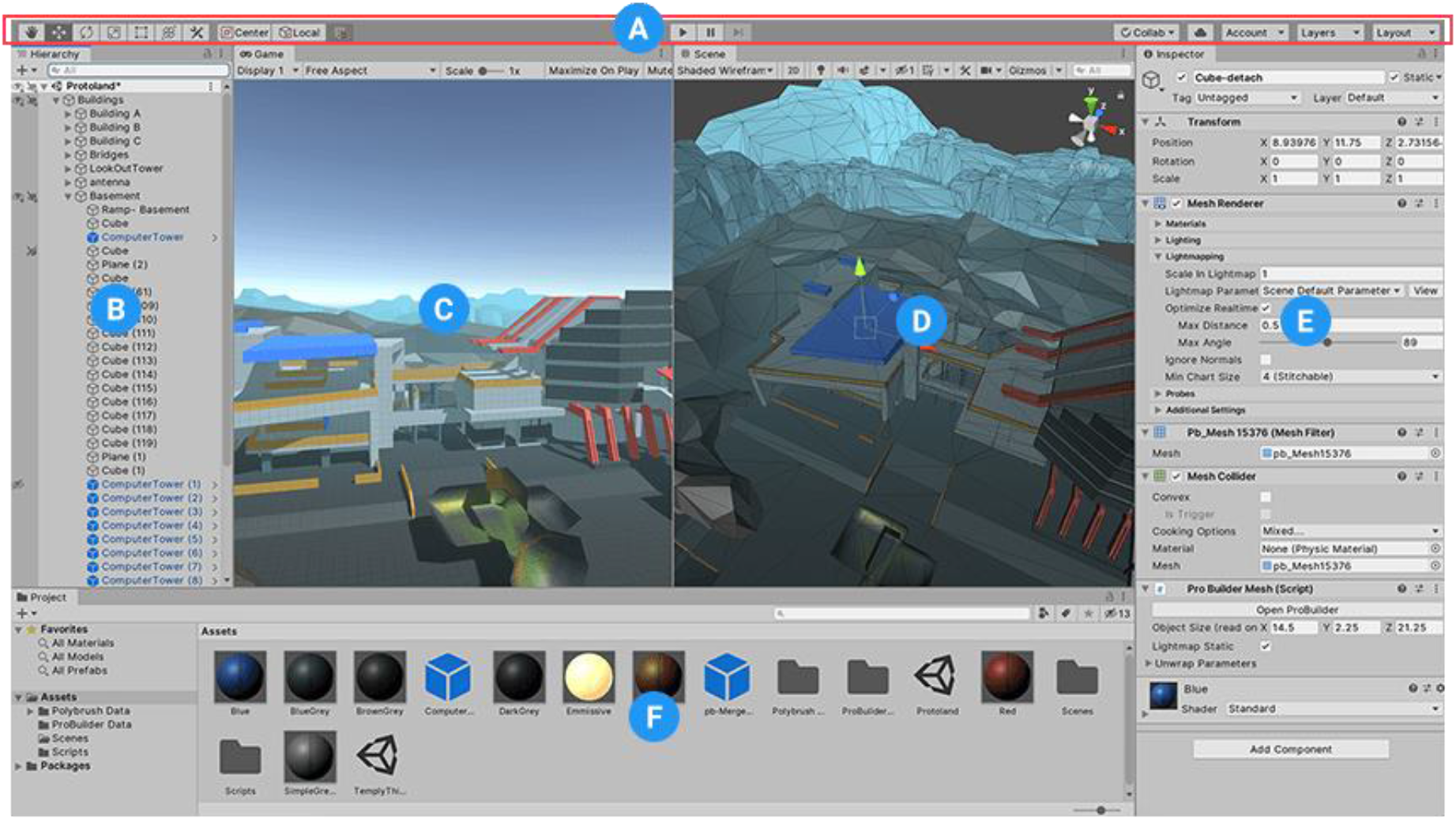
(A) Left: tools for manipulating the Scene view and the GameObjects within Scene; Centre: play, pause and step buttons. (B) Hierarchy of every GameObject in the Scene shows how GameObjects attach together. (C) The Game view shows the final rendered game. The play button can start the game simulation. (D) The Scene view allows to visually navigate and edit your Scene. (E) The Inspector Window allows to view and edit all the properties of the currently selected GameObject. (F) The Project window shows the imported Assets of the game.

We created a multisensory experiment by embedding **LSL** in the **PsychoPy** custom code component and **Unity**. This process is discussed in detail in sections 4.4 and 5 for **PsychoPy** and **Unity**, respectively.

### Lab Streaming Layer

The Lab Streaming Layer (**LSL**) connects a **Psychopy** experiment or a game in **Unity** (stimulus presentation and data acquisition) with multiple sensor devices. **LSL** consists of a core library and applications built on top of that library, which allows synchronous collection of multiple time series in research experiments and recording the collected data on disk. In **LSL**, a single measurement from a device (from all channels) is called a **Sample**. Samples can be sent individually for improved latency or in chunks of multiple samples for improved throughput. All the information about data streams (series of sampled data) is sent through an XML as the metadata which contains **name, type, channel_count, channel_format** and **source_id** for each stream. Through Stream Outlet, chunks or samples of data are made available on **LSL** network and these streams are visible to all the computers connected to the same local network, LAN (Local Area Network) or WLAN (Wireless Local Area Network). Streams can be distinguished with their assigned name, type and other queries created on the XML metadata. The streams that are available on the **LSL** network can be received by the computers that call the Stream Inlet.

To define a stream in **LSL**, stream outlets are created by calling StreamOutlet functions that take StreamInfo as the input. StreamInfo includes name, type, number of channels, channel format (string, int, float, etc.) and source ID as the input. Name, type and source ID are arbitrary and can be defined by the user; the number of channels and channels format depends on the characteristics of the streams. Then whenever needed, data can be transported by calling the functions that push the data in Samples or Chunks to the **LSL**. Once all the stream outlets are available on the **LSL** network, they can be saved in a single XDF using **LabRecorder** application, or they can be received in another platform by means of StreamInlet functions.

This complicated data acquisition and synchronization process can be semi-automated by **LabRecorder**, which is explained in the next section.

### LabRecorder

**LabRecorder** is responsible for recording the streams available on **LSL** on the hard disk (it can be downloaded from https://github.com/labstreaminglayer/App-LabRecorder/releases). Figure 6 shows **LabRecorder** GUI; names of the streams that are available on **LSL** can be seen on the left panel. However, in the latest version of **LabRecorder**, there is no need for **LabRecoder** GUI to record the data. **LabRecorder** can start recording the available **LSL** streams by writing one line of code that opens the **LabRecorder** command line interface (**LabRecorderCLI**) in the background. This process is explained in Sections 4 and 5.

**Figure 6.**
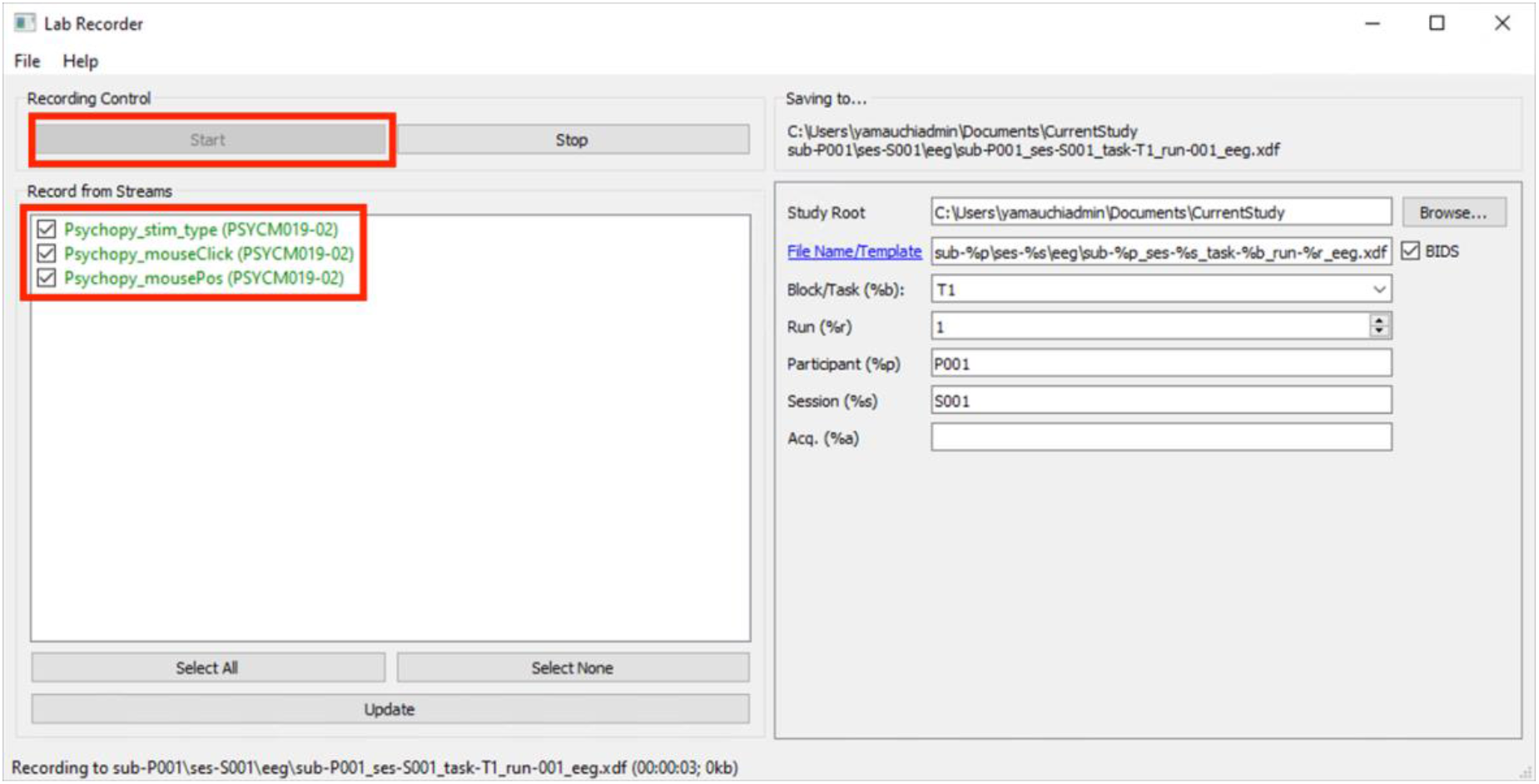
**LabRecorder** GUI

In sections 4 and 5, a case study explains how to integrate multiple devices such as EEG, GSR, Eyetracking, Bodymotion, mouse trajectories, button click, and task-related markers within a stimulus presentation software, **PsychoPy** (section 4) and **Unity** (section 5). The program automatically saves all the recording data into a single .*xdf* file in a user-specified folder. All these would be done with the aid of the **LSL** embedded in **PsychoPy/Unity**.

We used opensource software for the integration of different sensors and devices. Also, different consumer-grade and affordable devices are used, including 1) for EEG, g.tec Unicorn, Muse, Neurosky Mindwave [24], BrainProducts LiveAmp, OpenBCI Cyton (8-Channel) and OpenBCI Cyton + Daisy (16-Channel), 2) for GSR, Gazepoint biometrics device and also e-Health Sensor Platform v2.0 for **Arduino**, 3) for eyetracking Gazepoint, 4) for body motion, Microsoft Kinect Sensor V2 for Windows. A basic sample experiment in **PsychoPy** and a basic **VR** environment in **Unity** that integrate different devices are available on our GitHub:PsychoPy Example GitHub and VR Example GitHub. Table 1 shows the list of the devices used for the case study.

**Table 1.** The devices used for integration in the sample experiment.

| Type | Device | Method for sending data to LSL | Link |
| --- | --- | --- | --- |
| Mouse clicks/coordinates | Mouse | define LSL stream in PsychoPy | ----- |
| EEG | g.tec Unicorn | C++ application called from PsychoPy/Unity | https://github.com/moeinrazavi/Unicorn_LSL |
|  | BrainProducts LiveAmp | C++ application called from PsychoPy/Unity | https://github.com/moeinrazavi/LiveAmp-LSL |
|  | OpenBCI Cyton (+Daisy) | Python script called from PsychoPy/Unity | https://github.com/moeinrazavi/OpenBCI-LSL |
|  | NeuroSky Mindwave | NeuroSky Python library and functions in Psychopy/Call a Python script in Unity | ----- |
|  | Muse | Muse Python library and functions in Psychopy/Call a Python Script in Unity | Link 1: BlueMuse_Application<br>Link 2: BlueMuse_Installation_Guide |
| GSR | eHealth Sensor v2.0 Arduino Shield | Python script called from PsychoPy/Unity | https://github.com/moeinrazavi/eHealth-GSR-LSL |
|  | Gazepoint Biometrics Device | Python script called from PsychoPy/Unity | https://github.com/moeinrazavi/Gazepoint-Eyetracking-GSR-HeartRate--LSL |
| Eyetracking | Gazepoint | Python script called from PsychoPy/Unity | https://github.com/moeinrazavi/Gazepoint-Eyetracking-GSR-HeartRate--LSL |
| Body Motion | Kinect | C++ application called from PsychoPy/Unity | https://github.com/moeinrazavi/Kinect-BodyBasics-LSL |

Next two sections provide a step-by-step case study detailing the integration of multiple sensory devices using **PsychoPy/Unity, LSL** and **LabRecorder**. Also, it is explained how to make the process automatic for an easy-to-use multimodal-multisensory experiment.

## Case Study: Building a Multisensory Experiment (PsychoPy)

This section provides a tutorial for building a multisensory experiment by embedding **LSL** in **PsychoPy**. The task used in this case study is a simple version of the Flanker task with only 4 trials. On each screen, participants are presented with a sequence of arrows →→→→→, ←←←←←, →→←→→or ←←→←← (Figure 7) and are asked to navigate their mouse cursor to a box on the top left or top right of the screen based on the direction of the center arrow in each sequence. Starting with this simple task, we show step by step how to add mouse/cursor motion, EEG (g.tec Unicorn, Muse, Neurosky Mindwave, BrainProducts LiveAmp, OpenBCI Cyton and OpenBCI Cyton + Daisy), GSR (e-Health Sensor Platform v2.0 for Arduino and also Gazepoint biometrics device), Eyetracking (Gazepoint), and Body Motion (Kinect) one by one to the experiment.

**Figure 7.**
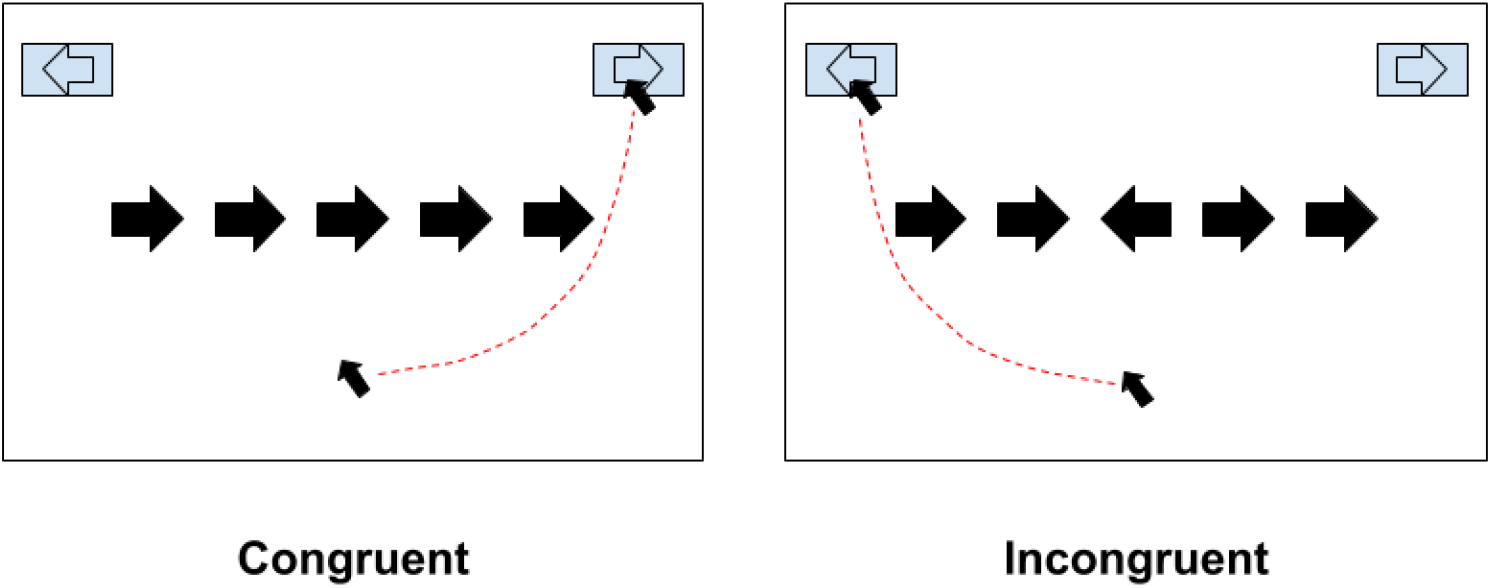
Arrow Flanker task

### 4.1. Software and Plugin Installation

**PsychoPy** provides a graphical user interface for designing a wide range of psychological experiments without any programming (see Appendix for the installation process). For example, text/picture can be added in different routines as stimuli by adding text/picture items, and also keyboard/mouse can be added as items for recording response. **PsychoPy** uses Python programming language in the background; custom Python code items can be used to add the features which are not available in the **PsychoPy** GUI (e.g., **LSL**). Routines and loops can be added for repeating one or several routines, including the stimuli, user’s response, and sending their markers to **LSL** in the experiment. Then all of these (routines, loops and custom code in routines) can be compiled and run together using **PsychoPy**. **pylsl** is a Python library that allows using **LSL** in Python (see Appendix for installing **pylsl** on **PsychoPy**). Module **Popen** from python **subprocess** library is used for sending data from different devices to **LSL**. Then, using module **popen** from python **os** library enables us to use **command line** to open **LabRecorder** in the background of the experiment. This will save all the streams automatically without needing to open the **LabRecorder** user interface (explained in section 4.3).

### 4. 2. Design

As shown in Figure 8, the example experiment contains 6 routines and one loop for the last 3 routines, which repeats the stimulus presentation. There are 3 main routines named **Initialize, Record_Start** and **Stimulus_Presentation**. The **Initialize** routine defines all the streams from the task that are intended to be sent to **LSL**. In the **Record_Start** routine, we write a command to start recording all the streams that are available on **LSL** in a .*xdf* file. Finally, in the **Stimulus_Presentation** routine, we write the script to send the task stimulus markers and mouse coordinates to **LSL**.

**Figure 8.**
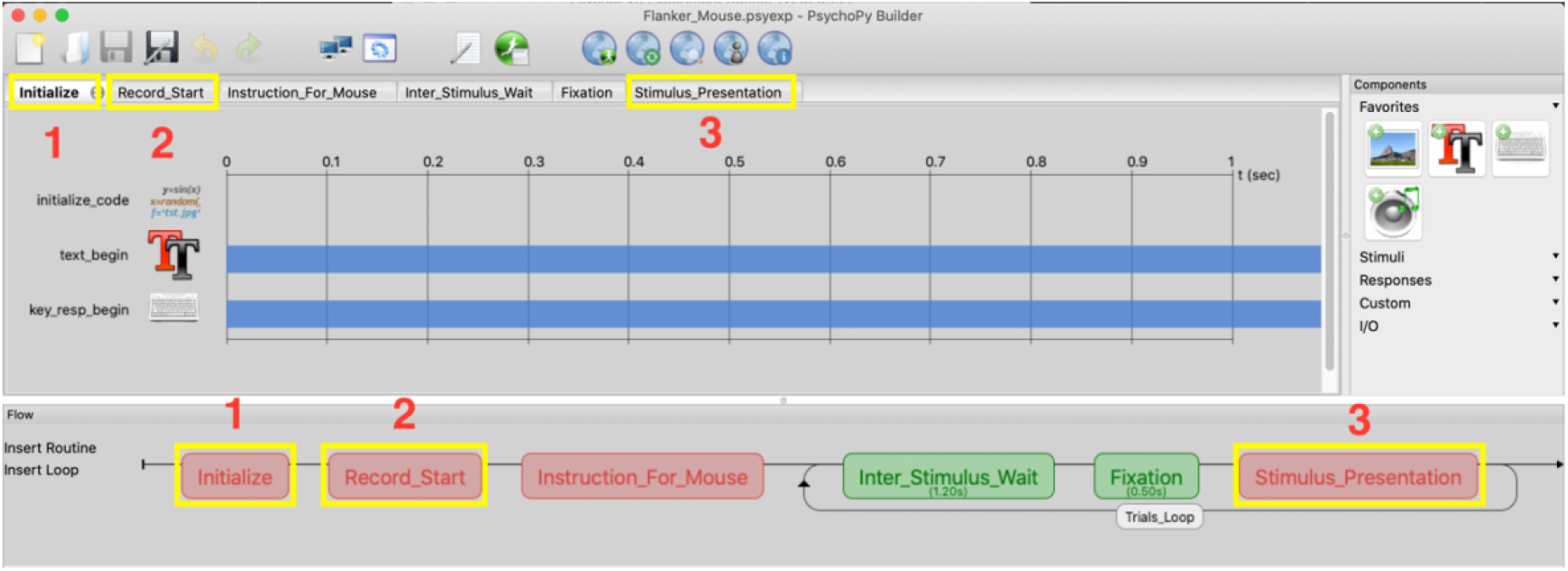
Main components of the example multisensory experiment: 1) initializing **LSL** streams 2) start recording **LSL** streams 3) stimulus presentation and sending stimulus and response markers to **LSL**

In the **Experiment Settings**, as shown in Figure 9, one of the fields is defined as **UIN**, which would be used in the recorded .*xdf* file name.

**Figure 9.**
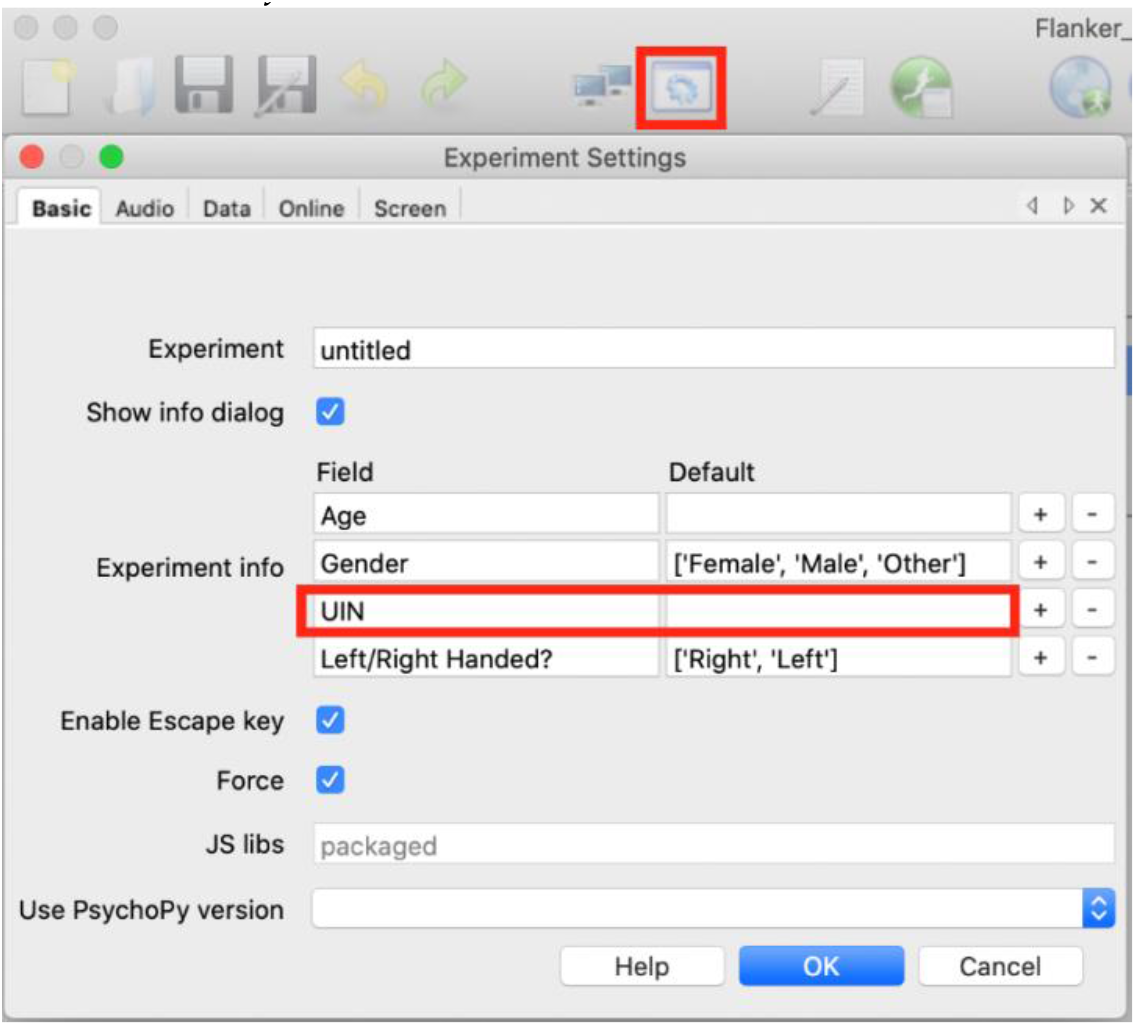
**PsychoPy** Experiment Settings

In the experiment folder, there is a folder, *Stimuli*, and a file, *stimuliFile.xlsx*, inside it, which contains **stimID, stim_type, stim, corr_key** (shows the correct response) and **cong** (indicates whether the stimulus is congruent or incongruent) (Figure 10). This .*xlsx* file is the input for the Conditions of the loop component **Trials_Loop** (Figure 11).

**Figure 10.**
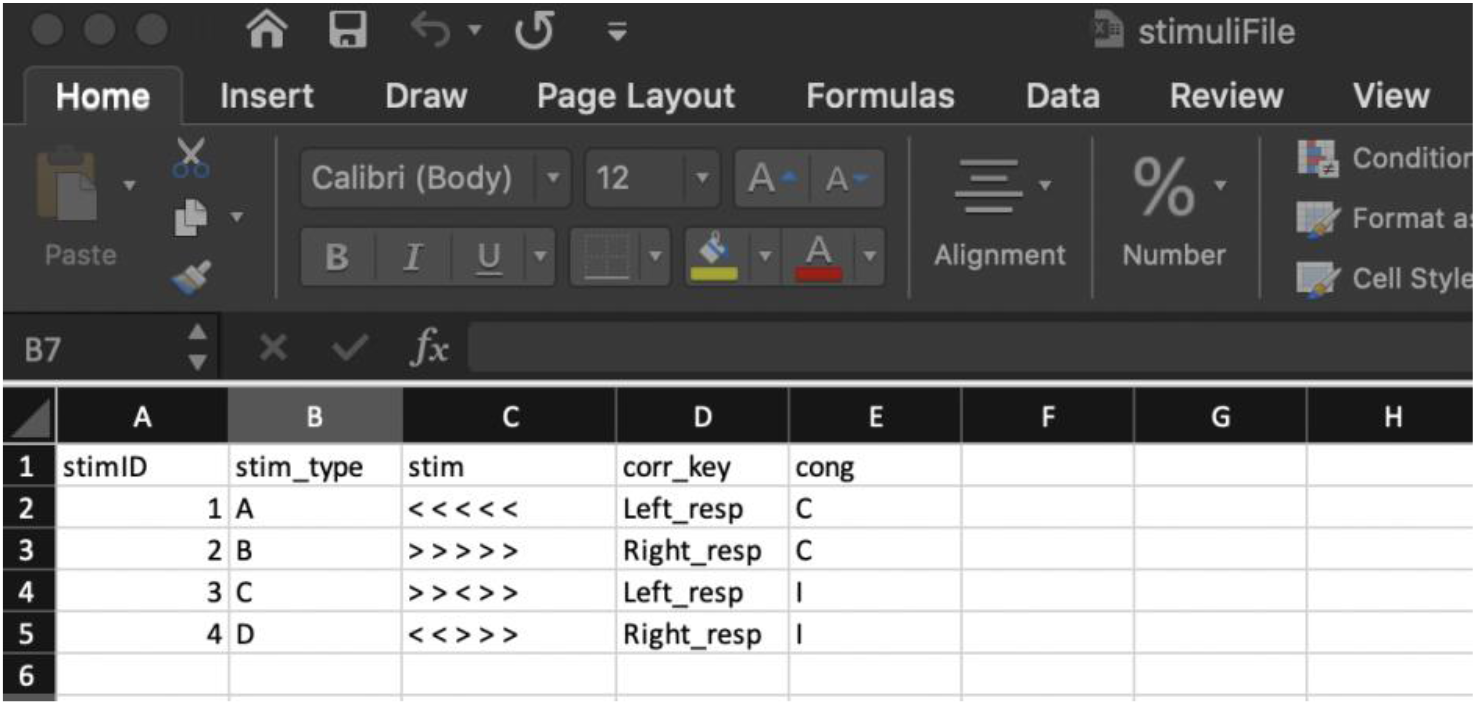
the Excel file used for stimulus presentation in our **PsychoPy** example experiment

**Figure 11.**
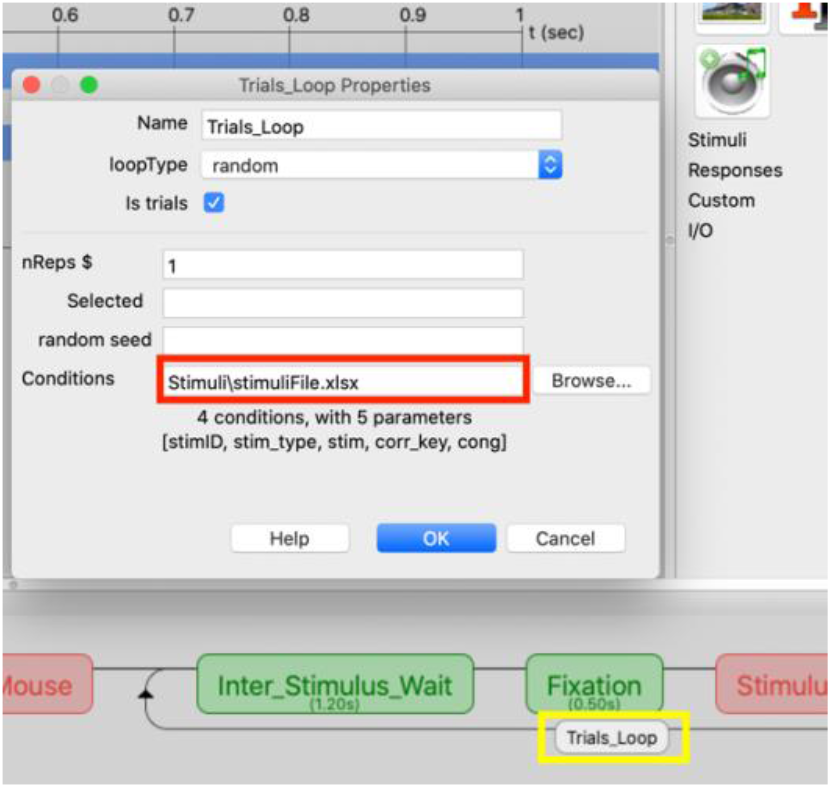
Choosing *stimuliFile.xlsx* as the input for the Conditions of the loop

### 4. 3. Adding Automatic Data Acquisition to the Experiment

To start recording the data streams available on **LSL** into a .*xdf* file, the code script shown in Figure 12 is added in the custom code named **xdf_record,** which is inside the **End Routine** tab of the **Recording_Start** routine. This script uses Windows **command line** to open the **LabRecorder** in the background of the experiment and starts recording the data. It is important to note that all the streams are created and initialized before the script that calls **LabRecorder** in the background.

**Figure 12.**
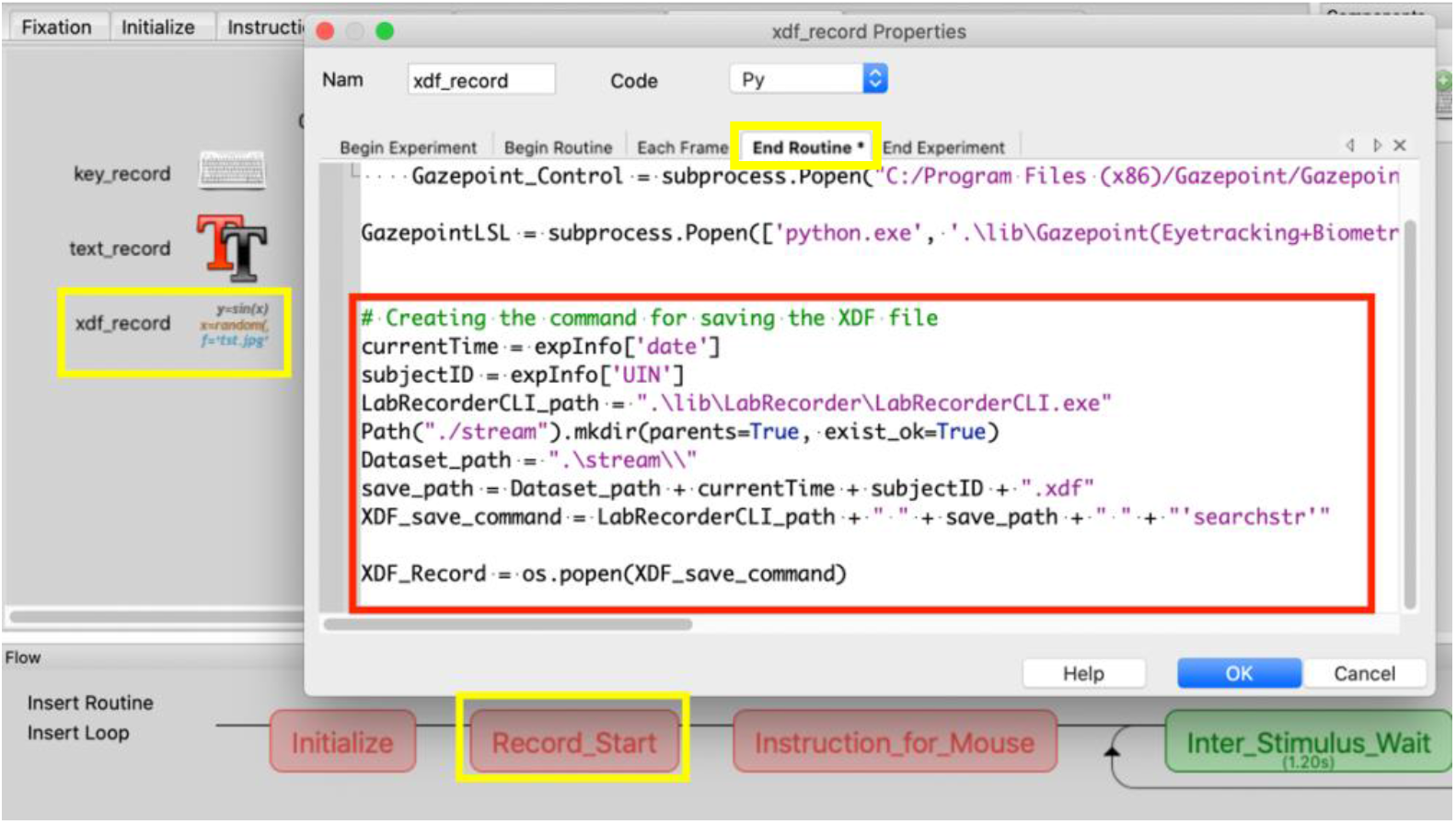
**PsychoPy** custom code from starting **LabRecorder** automatically from **command line**

In the following, a step by step process on how to add different streams to the experiment is shown.

#### 4.3.1 Mouse data + Flanker task markers

After opening the experiment *Flanker_Mouse.psyexp*, inside the **Initialize** routine, there is a custom code named **initialize_code.** After opening **initialize_code**, in the **Begin Experiment** tab, first, the required modules are imported (Figure 13), then 3 **LSL** streams are created. The first stream is for sending stimulus markers (Figure 14), the second stream is for sending user response markers (Figure 15), and the third stream is for sending mouse coordinates to **LSL** (Figure 16).

**Figure 13.**
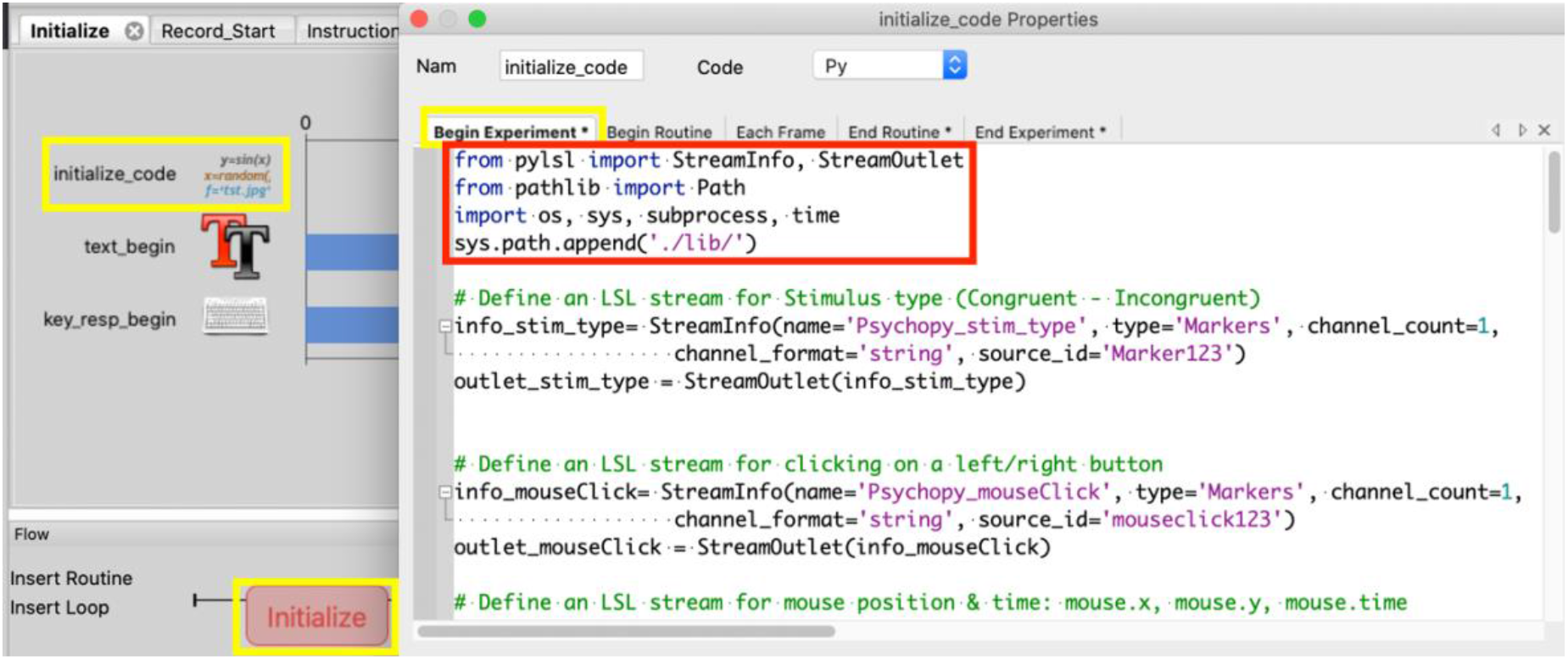
Importing the required modules

**Figure 14.**
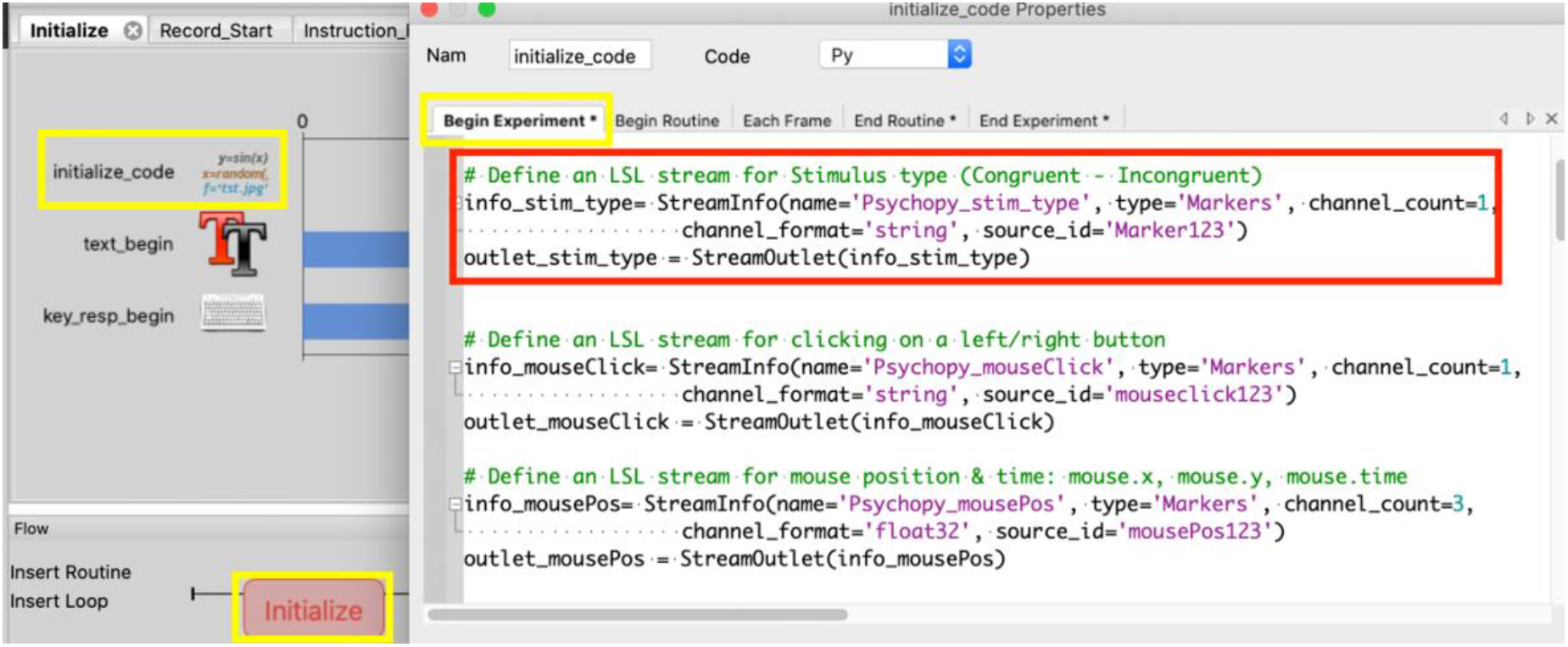
Defining a stream for sending stimulus type (congruent/incongruent) to **LSL** in each stimulus presentation

**Figure 15.**
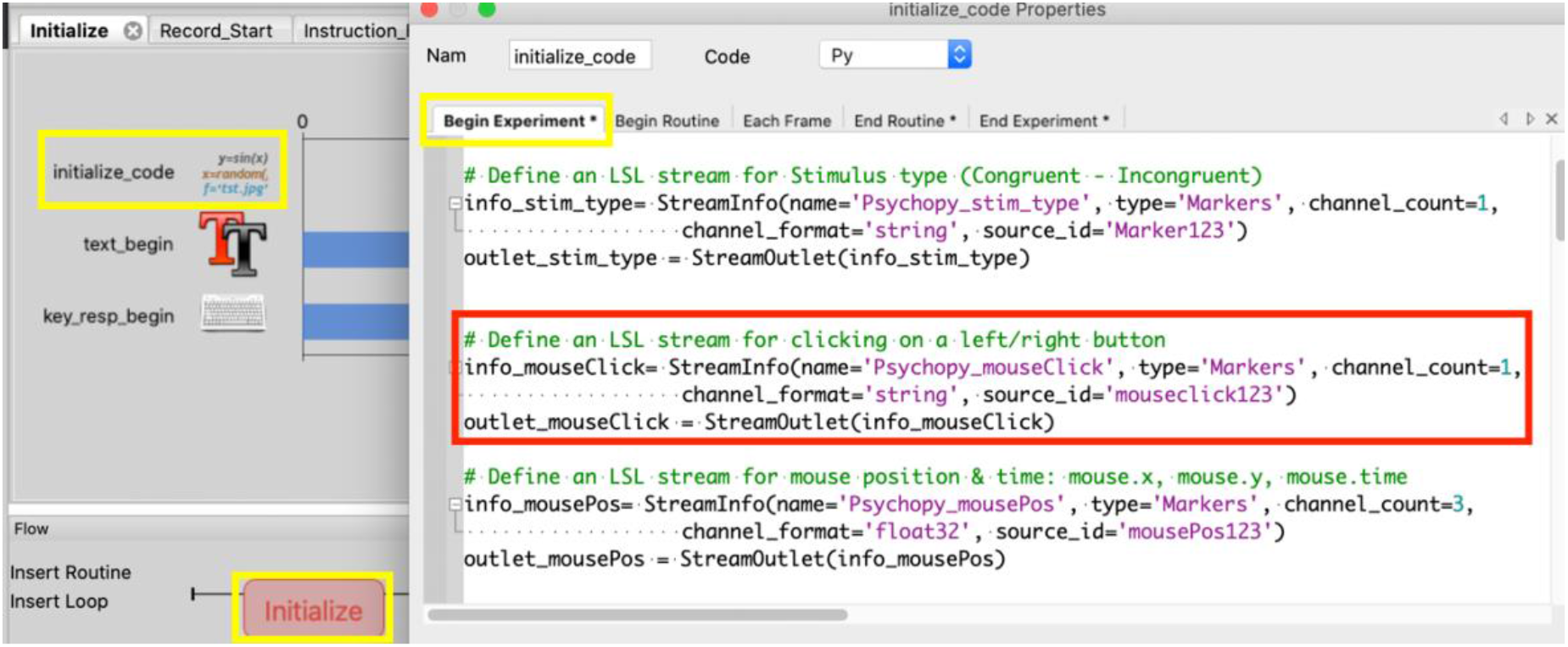
Defining a stream for sending user response (clicking on the left/right box) to **LSL**

**Figure 16.**
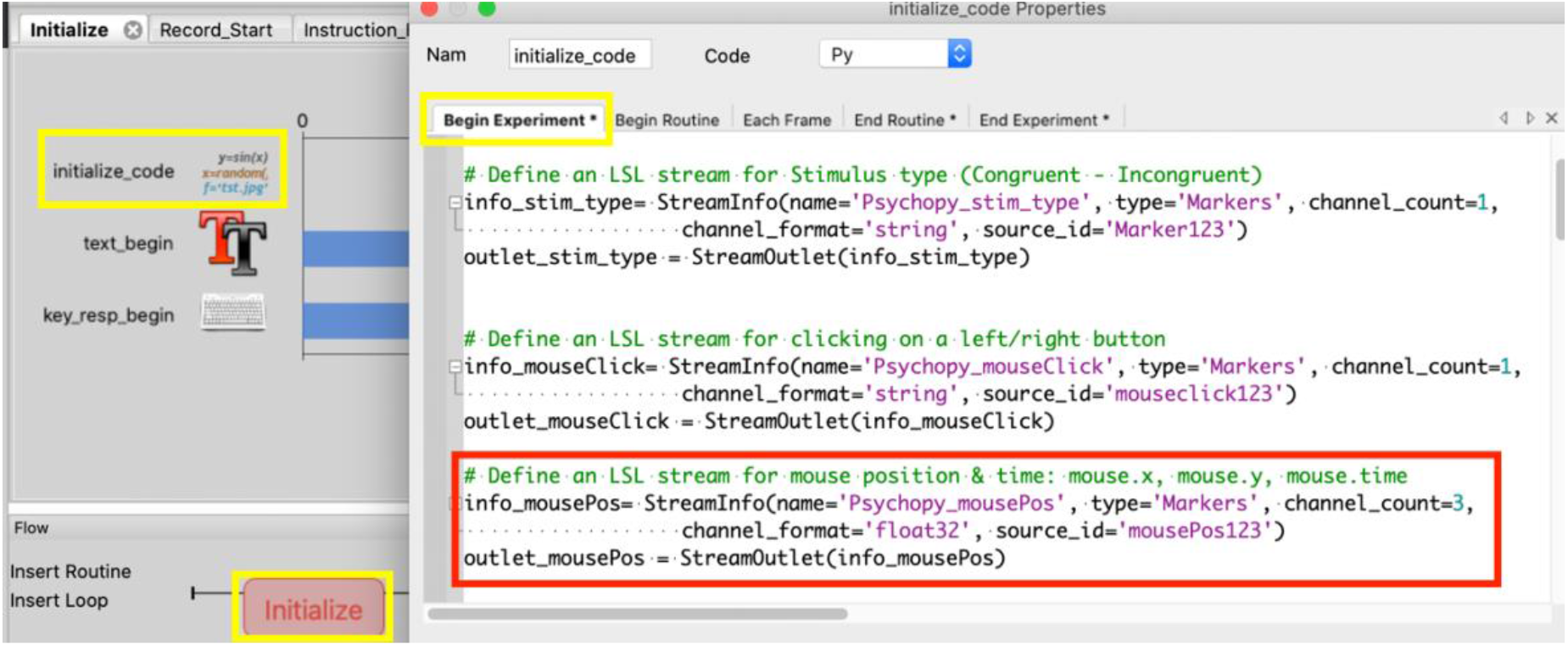
Defining a stream for sending mouse coordinates to **LSL** in each trial

In order to send a stimulus marker to **LSL**, in the **Begin Routine** tab of the custom code **stimulus_code**, which is inside the **Stimulus_Presentation** routine, we added the code shown in Figure 17. In order to send the user response data to **LSL**, in the same custom code, we added the code shown in Figure 18 in the **End Routine** tab (since the option “End Routine on valid click” is selected for the **mouse** component in the **Stimulus_Presentation** routine). Finally, to send mouse coordinates data to **LSL**, in the **Each Frame** tab (since the mouse coordinates should be sent in each refresh of the monitor screen), we added the code shown in Figure 19. Similarly, if keyboard is used instead of mouse for user response, to send keyboard data to **LSL, key_resp.keys** should be used as the argument for the outlet.push_chunk/outlet.push_sample (This is not shown here for brevity).

**Figure 17.**
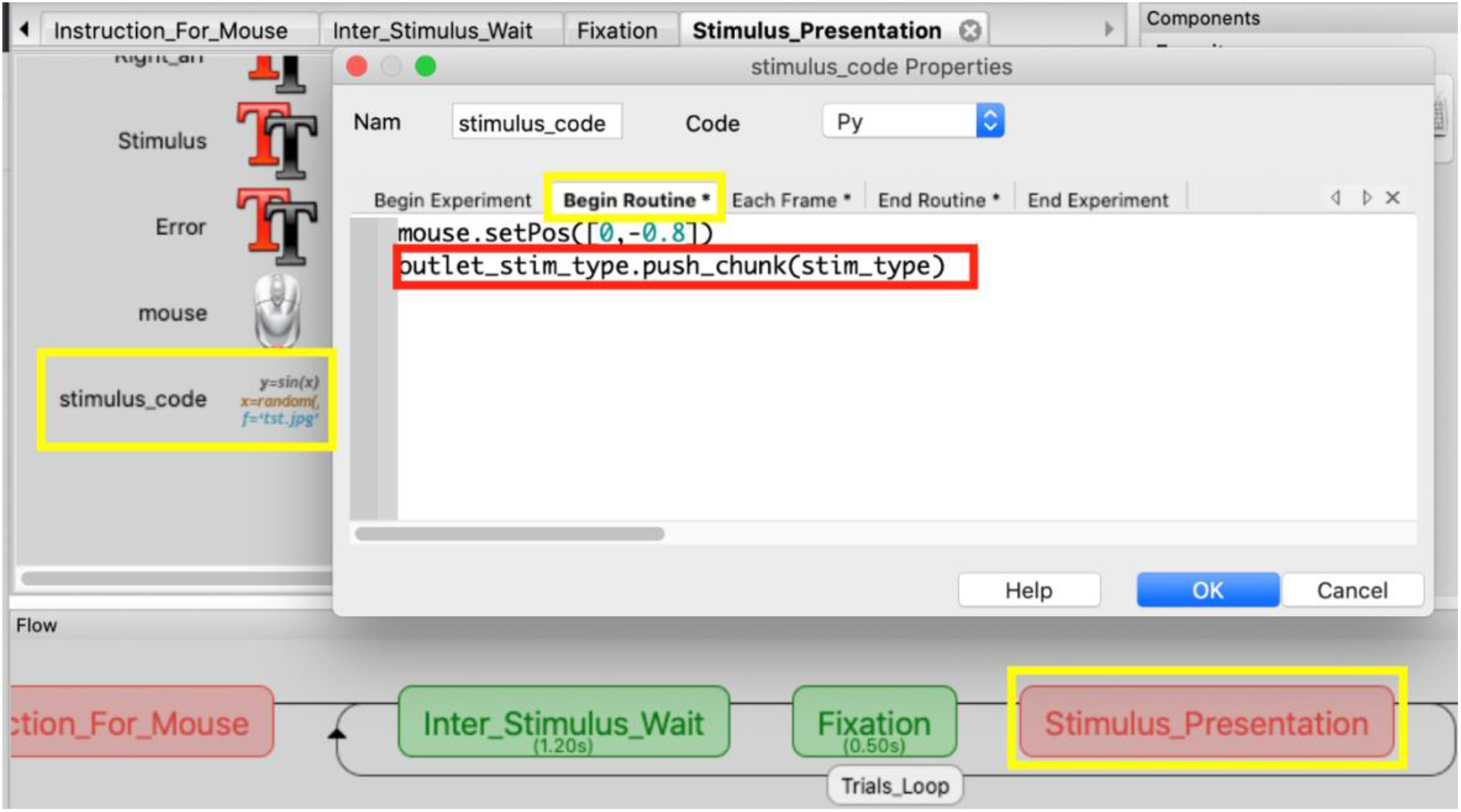
Sending the type of stimulus to **LSL** at the onset of stimulus presentation on the screen

**Figure 18.**
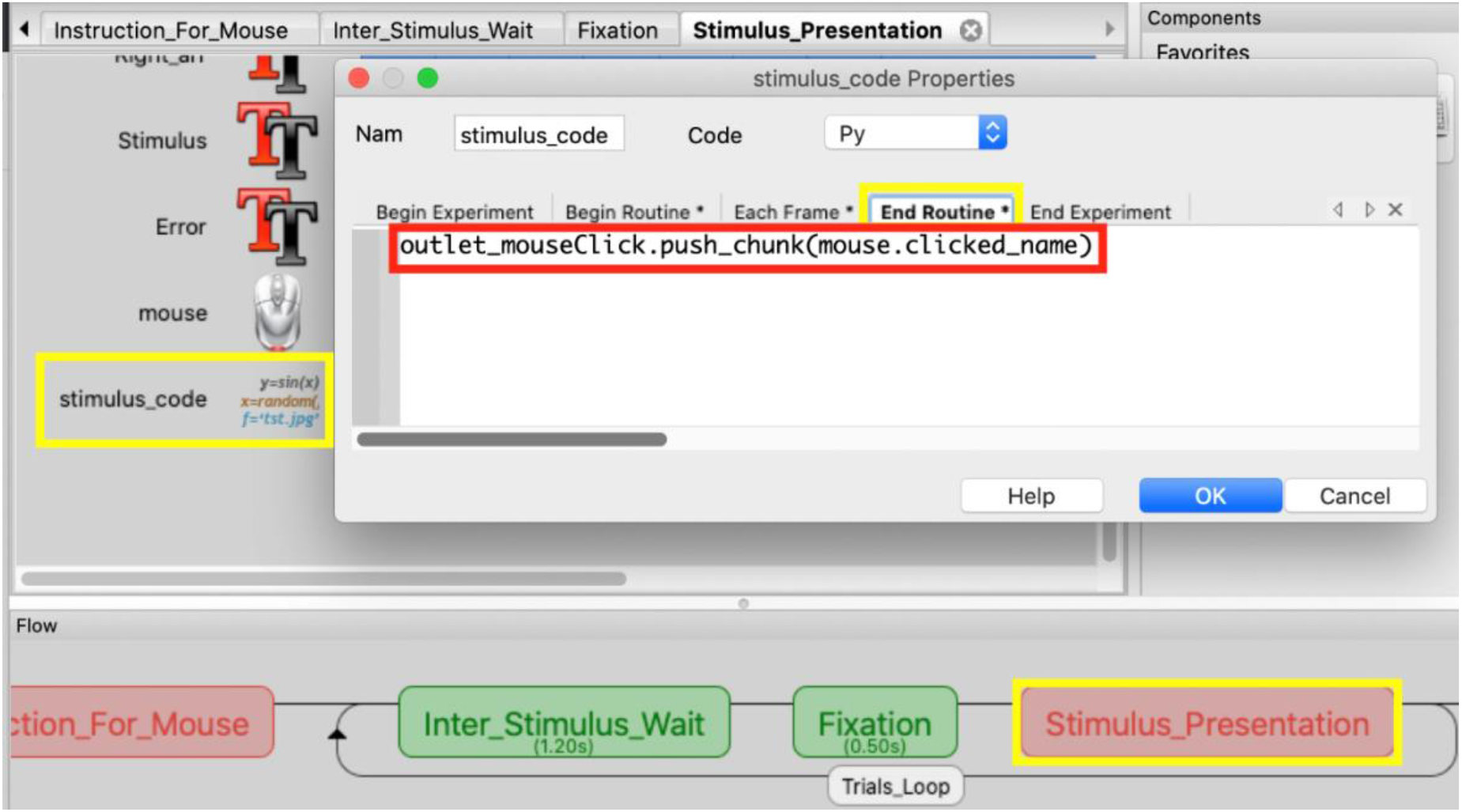
Sending the user response data (whether the user clicked on the left/right box) to **LSL** at the moment he clicks on the left/right box.

**Figure 19.**
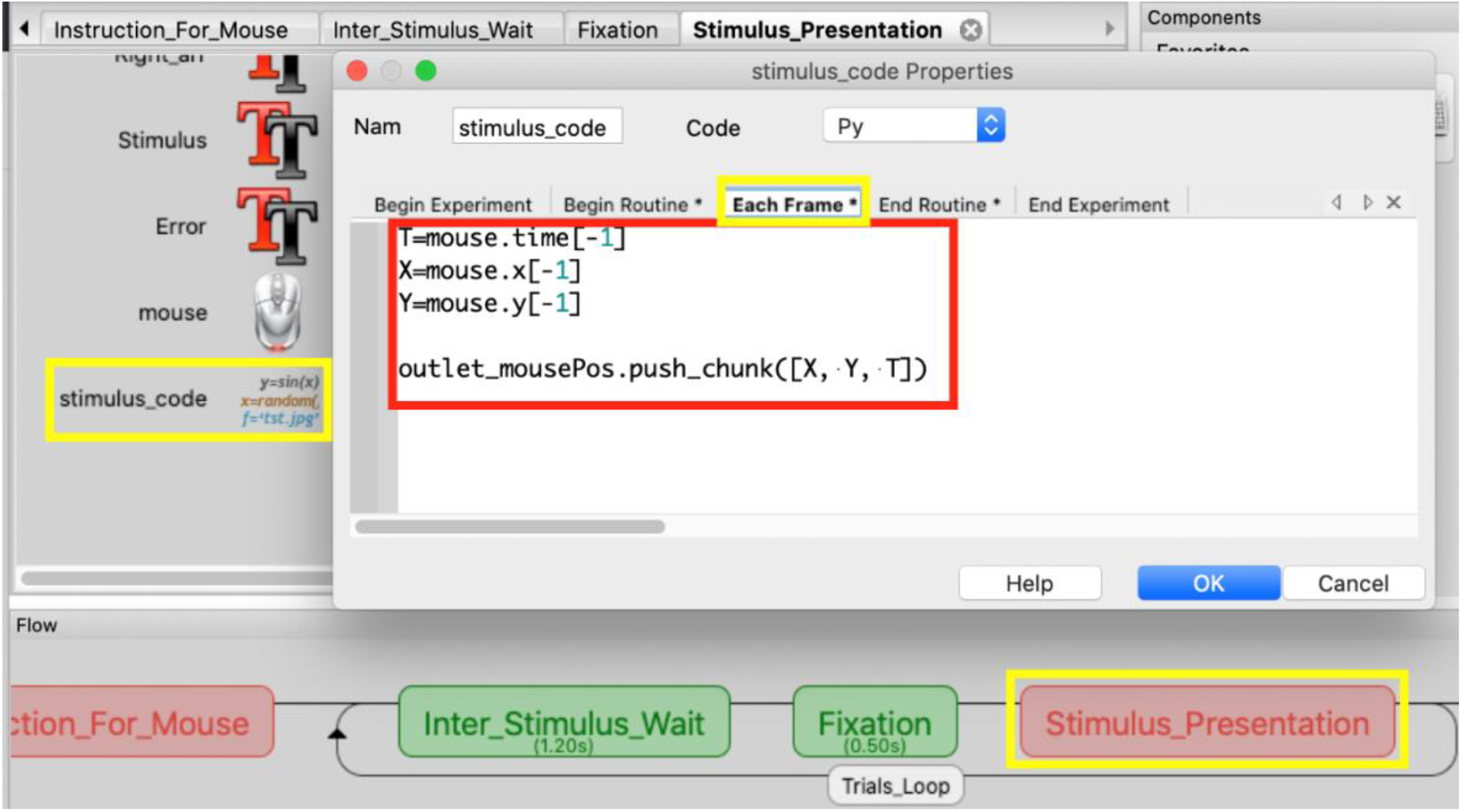
Sending mouse coordinates in each refresh of the monitor screen to **LSL**

Features such as maximum velocity, maximum acceleration, total distance of the mouse movement, area under the curve, maximum absolute deviation from a straight line, etc. can be extracted easily from Mouse data by a package for R and Python called *mousetrap* (Kieslich 2017).

#### 4.3.2 Mouse Data + Flanker task markers + EEG

The following shows how to add different EEG devices to the experiment.

- **g.tec Unicorn (8-channel EEG device)**

We have developed a C++ program (using Unicorn SDK) to send the data from Unicorn device to **LSL**. The application can be downloaded from our GitHub and copied inside the experiment *lib* folder. Then the application can be simply called from **PsychoPy** by adding the script shown in Figure 20 in the **Begin Experiment** tab of the **initialize_code** inside the **Initialize** routine. This will automatically send data from a paired Unicorn device to **LSL**.
- **BrainProducts LiveAmp (16, 32 and 64-channel EEG device)**

We have developed three C++ applications (using LiveAmp SDK) to send data from 16, 32 and 64-channel LiveAmp devices to **LSL**. The application associated with the desired device can be downloaded from our GitHub and copied inside the experiment *lib* folder. Then, the application can be called from **PsychoPy** by adding the script shown in Figure 21 in the **Begin Experiment** tab of the **initialize_code** which is inside the **Initialize** routine. This will automatically send data from a LiveAmp device to **LSL**.
- **OpenBCI Cyton (8-channel) and OpenBCI Cyton + Daisy (16-channel EEG Device)**

First pyOpenBCI needs to be installed using: **pip install pyOpenBCI**. Then *OpenBCILSL* folder should be download from our GitHub and copied in the experiment *lib* folder. Then, to send data from OpenBCI automatically to **LSL,** the code shown in Figure 22 should be added in **Begin Experiment** tab of the **initialize_code** inside the **Initialize** routine. In the *OpenBCILSL.py* file, in case Daisy module (16-channel) is used, **daisy = True** and the number of channels in the **LSL** stream info = 16; otherwise, **daisy = False** and the number of channels in the **LSL** stream info = 8 (Figure 23).
- **NeuroSky Mindwave (1-channel EEG device)**

First, **mindwavelsl** needs to be installed using the command **pip install mindwavelsl**. Then, the lines of code shown in Figure 24 should be added in the **Begin Experiment** tab of the **initialize_code**, which is inside the **Initialize** routine.
- **Muse (4-channel EEG device)**

First, **BlueMuse** needs to be downloaded from: https://github.com/kowalej/BlueMuse/releases/download/v2.1/BlueMuse_2.1.0.0.zip and installed as instructed in https://github.com/kowalej/BlueMuse. Then **muselsl** module should be installed using: **pip install muselsl.** To run **BlueMuse** automatically in the background when the experiment is started, the lines shown in Figure 25 should be added in the **Begin Experiment** tab of the **initialize_code** inside the **Initialize** routine.

**Figure 20.**
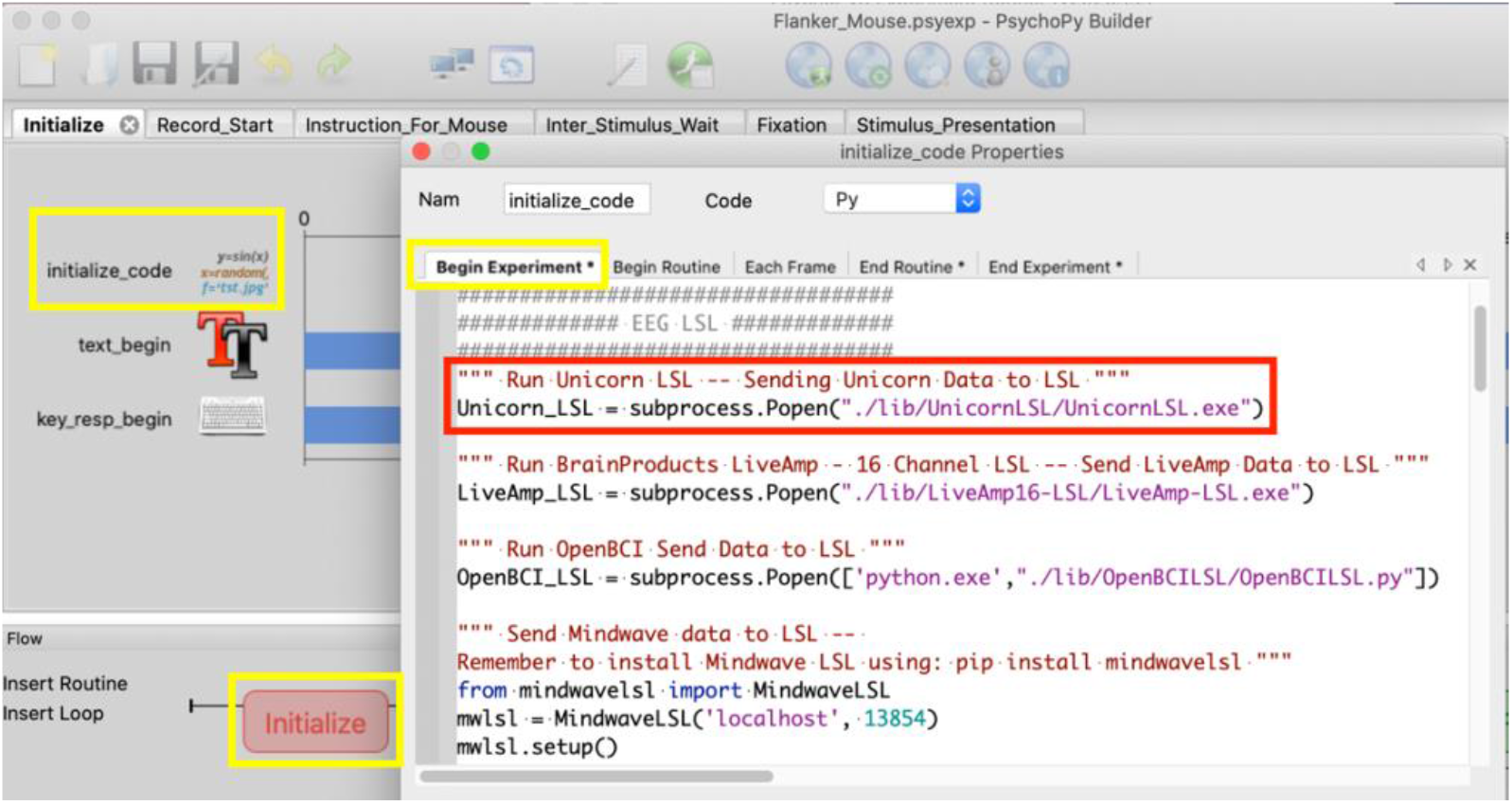
Running the application for sending g.tec Unicorn data to **LSL**

**Figure 21.**
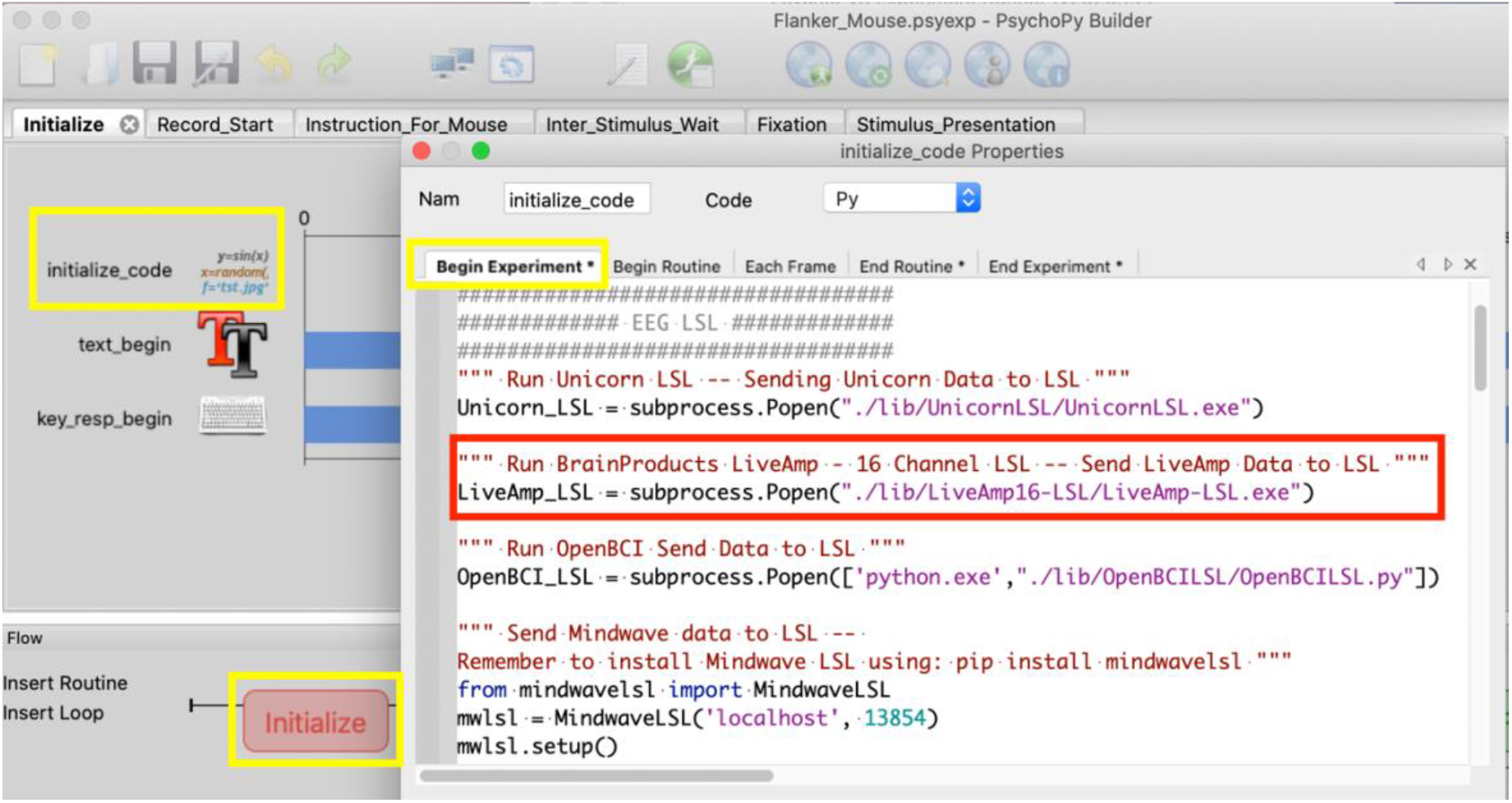
Running the application for sending BrainProducts LiveAmp data to **LSL**

**Figure 22.**
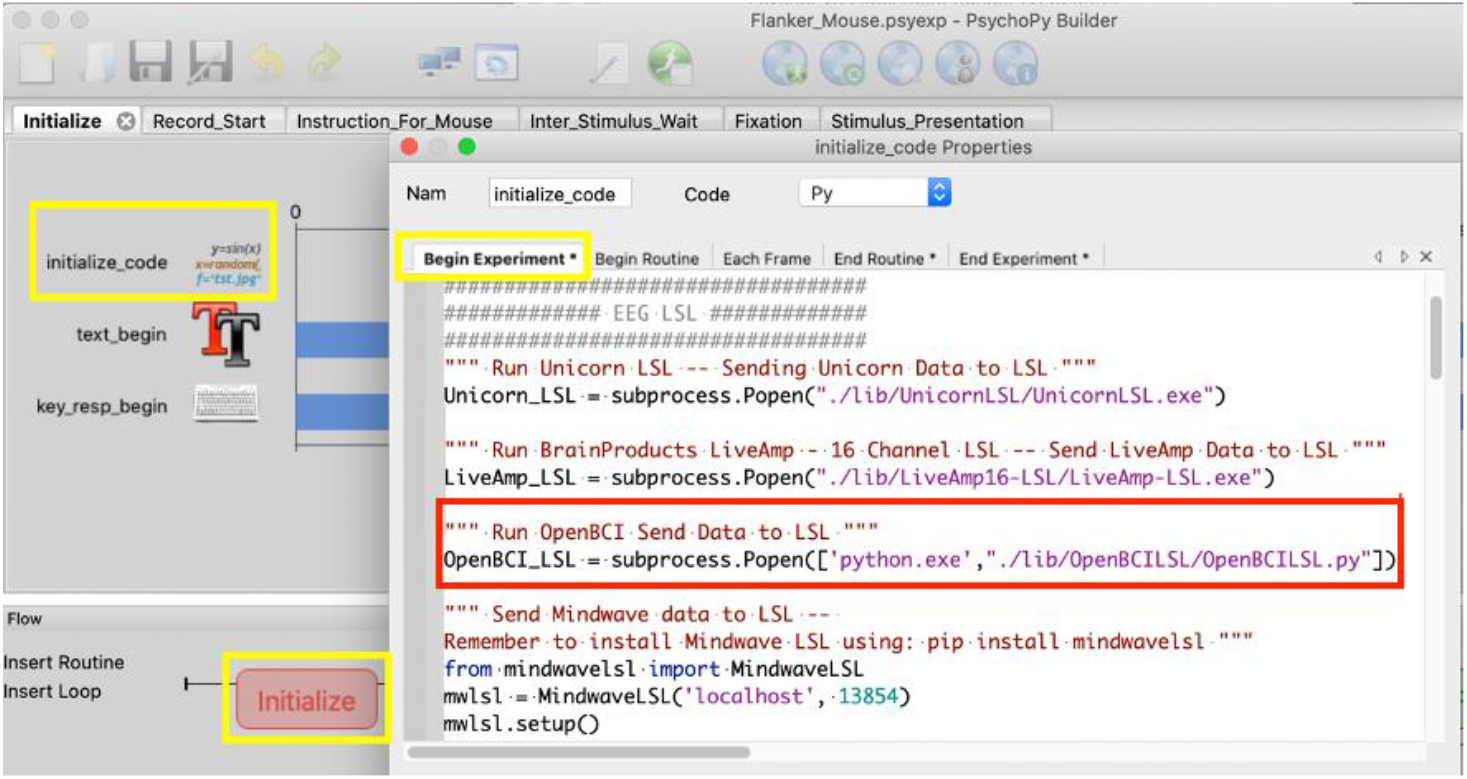
Calling the Python script for sending OpenBCI Cyton/OpenBCI Cyton + Daisy data to **LSL**

**Figure 23.**
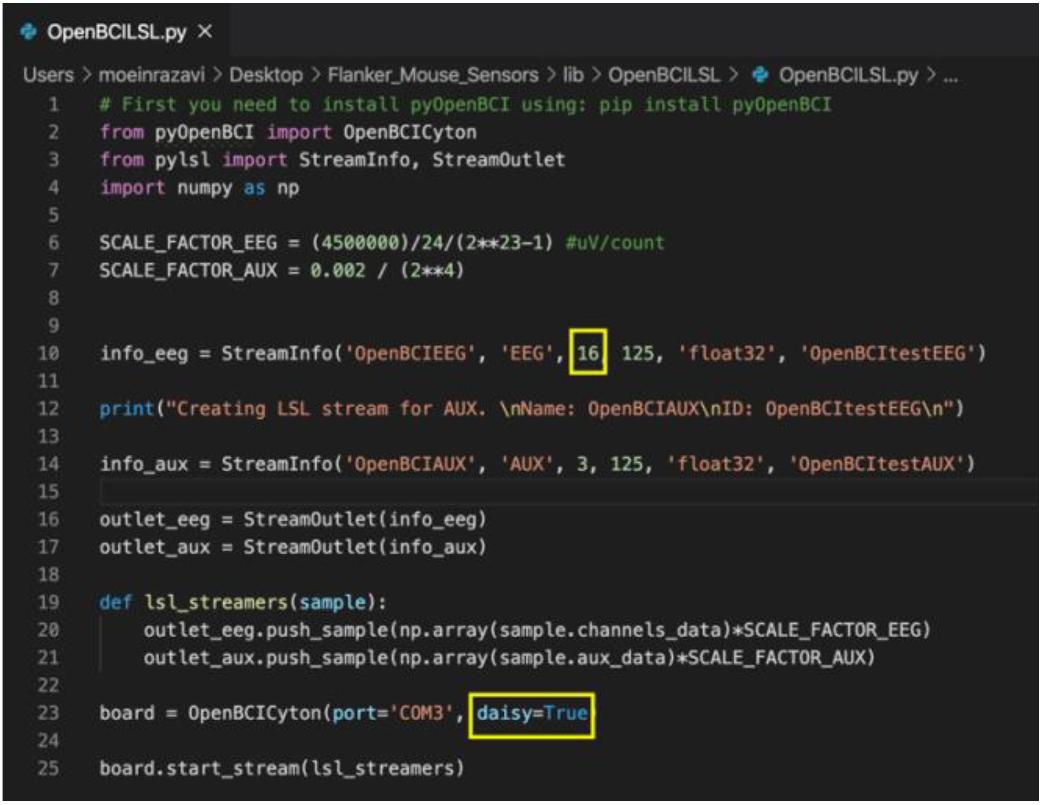
Python script for sending OpenBCI Cyton/OpenBCI Cyton + Daisy data to **LSL**

**Figure 24.**
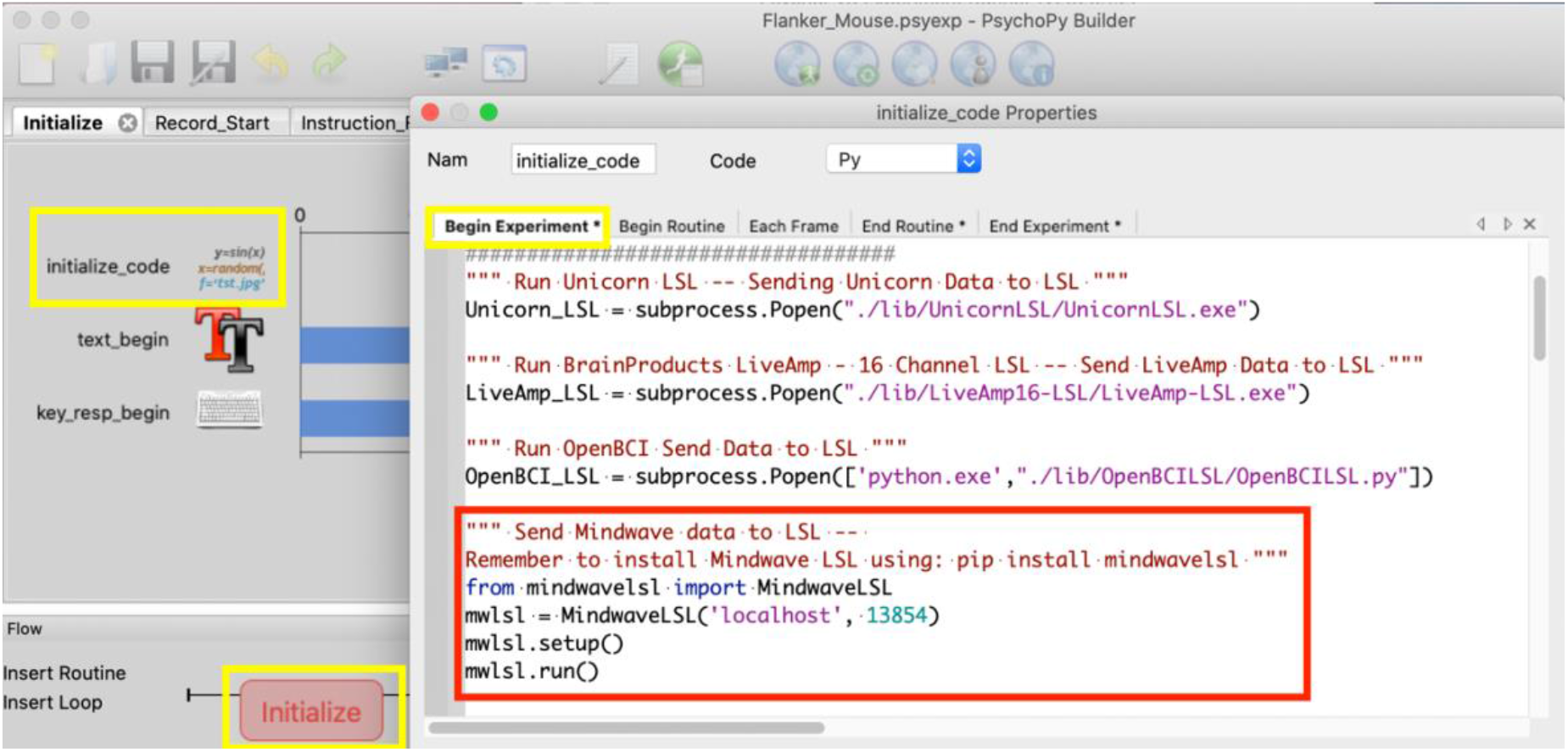
Sending NeuroSky Mindwave data to **LSL**

**Figure 25.**
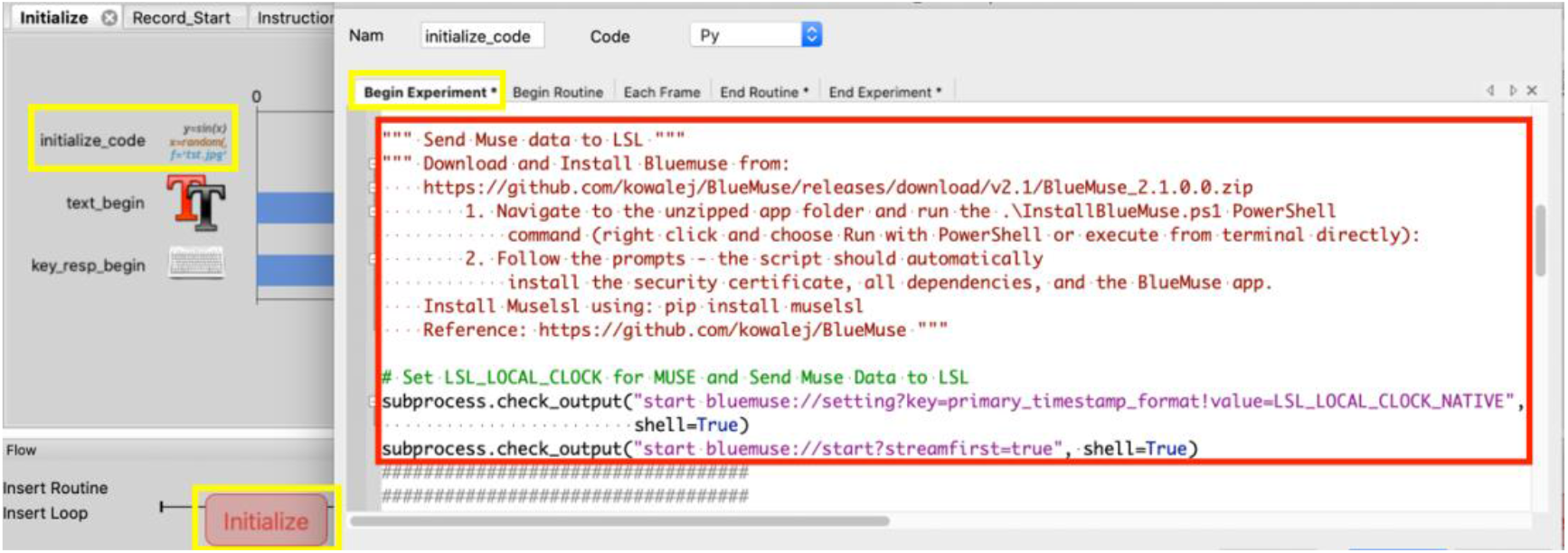
Sending Muse data to **LSL**

EEG data can be analyzed in **EEGLAB** (MATLAB Toolbox; https://sccn.ucsd.edu/eeglab/index.php) and Python **MNE** module. In order to read the .*xdf* files in **EEGLAB, xdfimporter** extension needs to be added to **EEGLAB**. To analyze data using Python MNE, **pyxdf** module should be installed for Python.

#### 4.3.3 Mouse Data + Flanker task markers + EEG + GSR (Arduino)

For GSR, 2 different devices are used, 1) eHealth Sensor v2.0 Arduino shield, and 2) Gazepoint Biometrics kit. For eHealth Arduino shield, first, add <eHealthDisplay.h> and <eHealth.h> libraries need to be added to Arduino IDE. To send data from Arduino to **LSL**, first its data should be sent to the Serial port with a script in Arduino IDE (https://www.arduino.cc/en/main/software) (Figure 26) and deploying this script on the Arduino device. Then, a separate Python/C/C++/etc. script can receive the data from the Serial port and send it to **LSL**. We have developed a Python script named *Serial2LSL.py*, which can be downloaded from our GitHub: https://github.com/moeinrazavi/eHealth-GSR-LSL. In this script the Serial port name needs to be changed based on the name of the port that the Arduino is connected to (Figure 27). Then, the Python script *Serial2LSL.py* should be called from **PsychoPy** as a subprocess (Figure 28). This will send the data from Arduino automatically to **LSL** when it is connected to the Serial port. The so-called Arduino IDE and Python scripts can be found in the *lib* folder.

**Figure 26.**
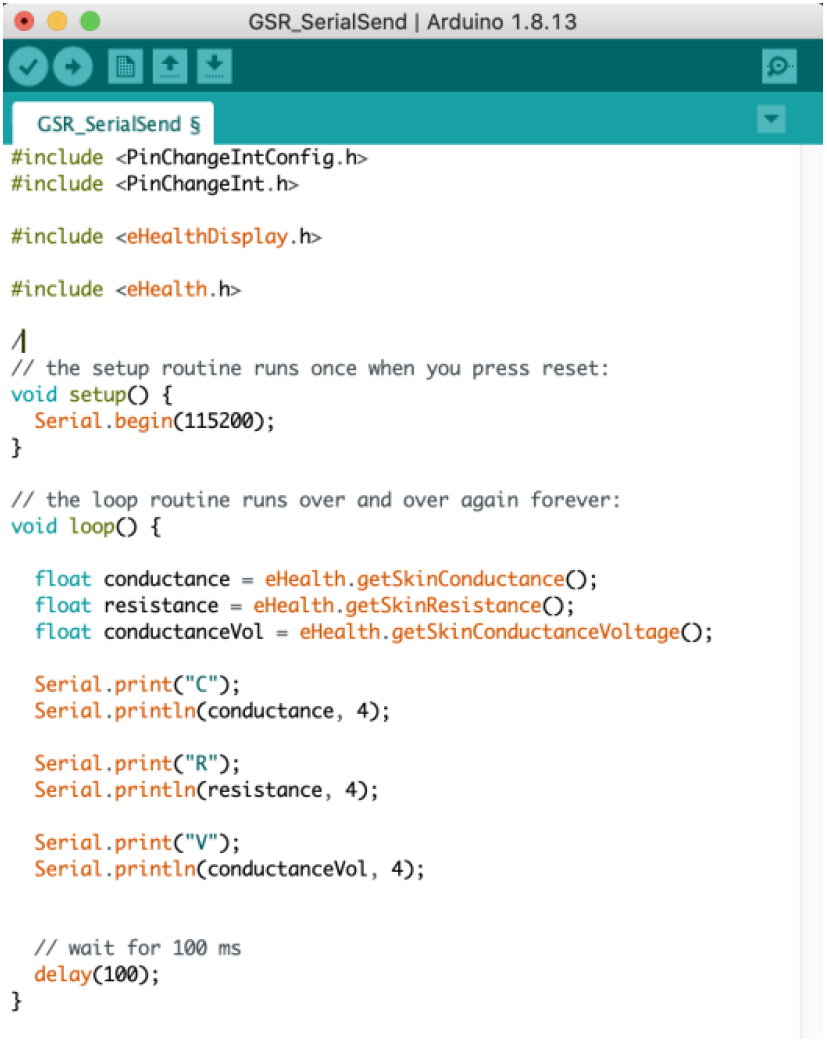
Arduino IDE script to send Arduino data to Serial port

**Figure 27.**
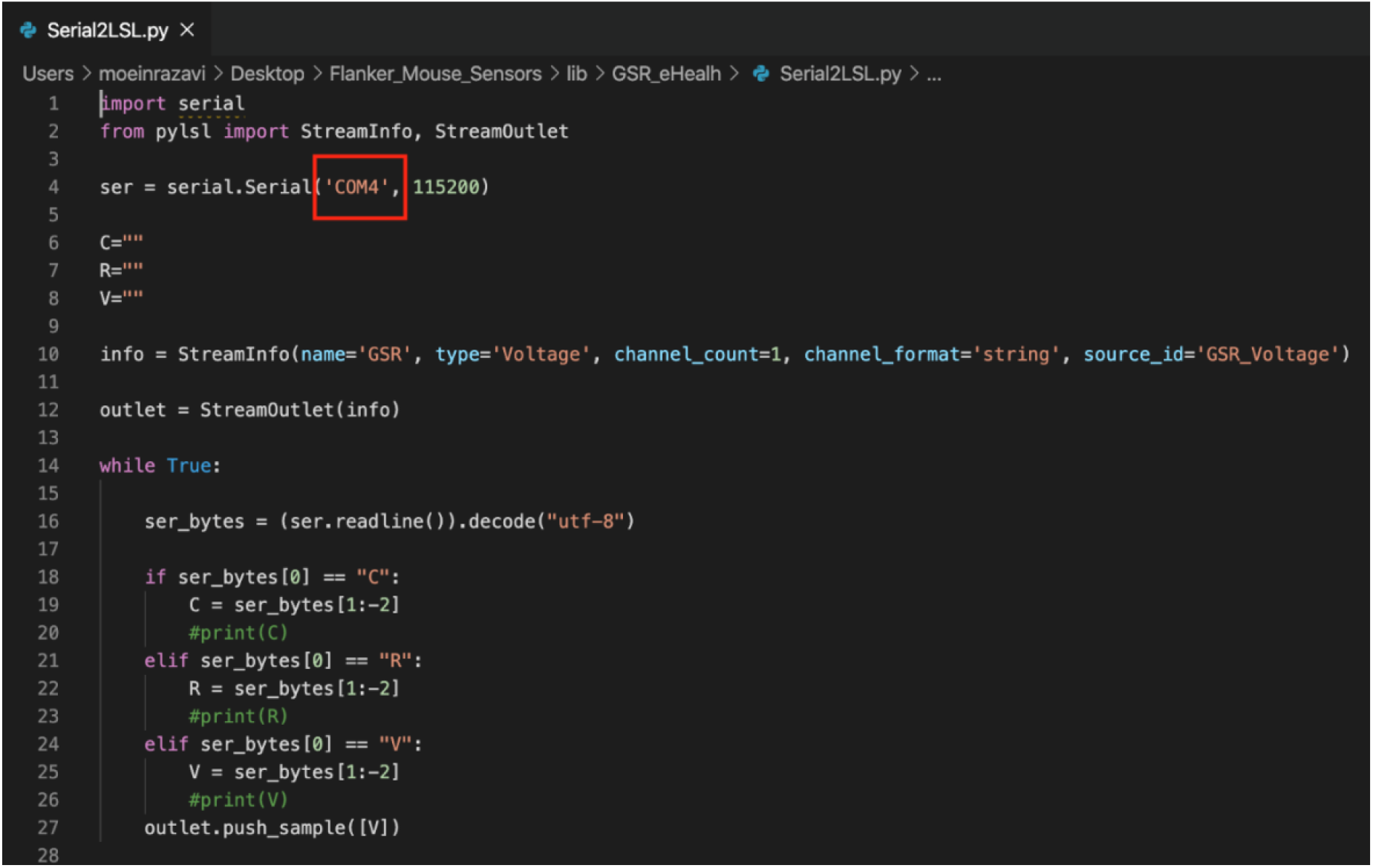
Python script for receiving data from Serial port and sending it to **LSL**

**Figure 28.**
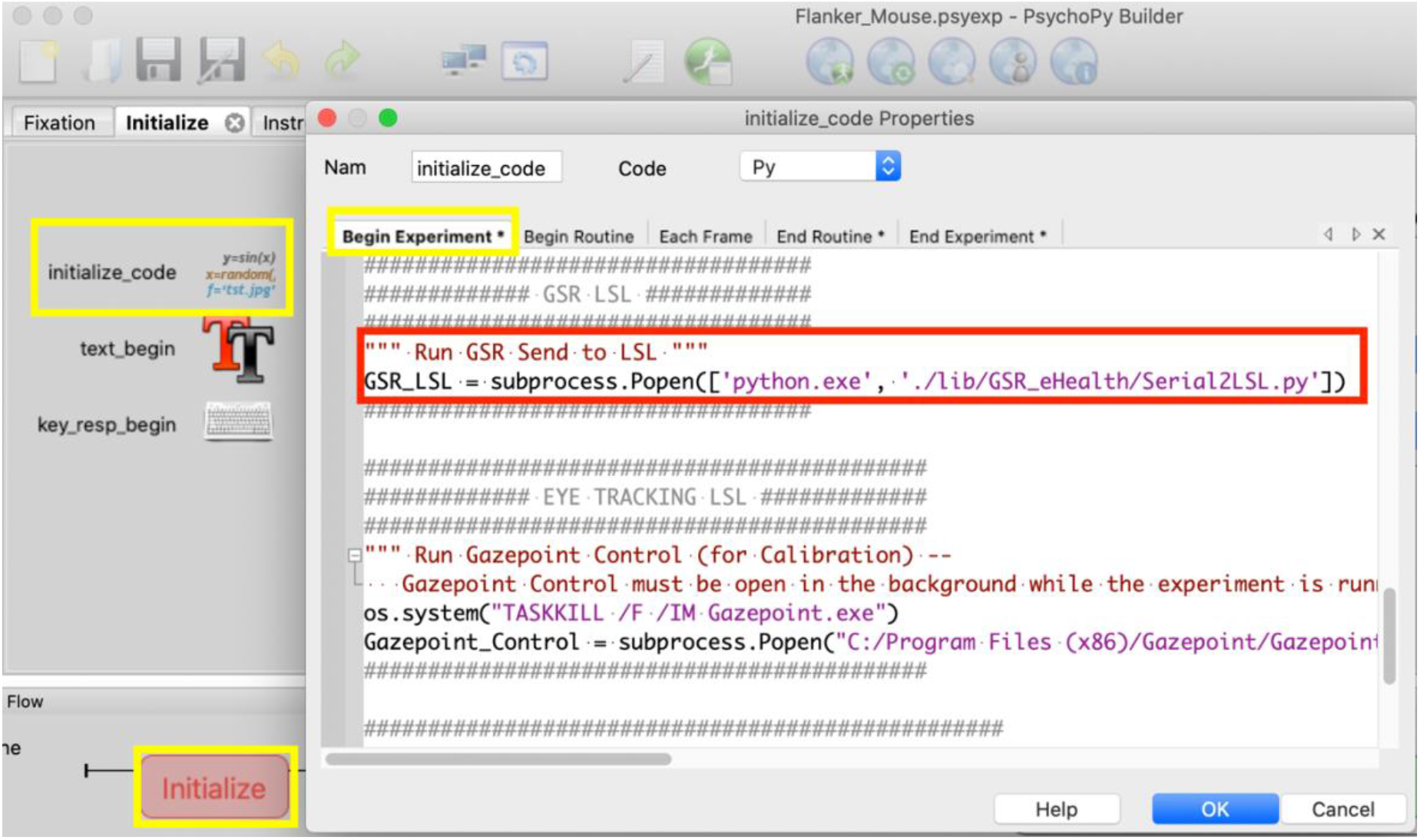
Calling the Python script for sending Arduino data to **LSL**

GSR data can be analyzed using **ledalab** (a MATLAB Toolbox) which can be downloaded from http://www.ledalab.de/. To analyze the data in **ledalab**, the .*xdf* file needs to be opened in MATLAB workspace (See Appendix). In the next subsection, it is shown how to send **GSR** data from the **Gazepoint Biometrics** kit (along with Gazepoint eyetracking data) to **LSL**.

#### 4.3.4 Mouse Data + Flanker task markers + EEG + GSR + Eyetracking

First, download the Gazepoint installer from https://www.gazept.com/downloads/ to install the Gazepoint Control application. This application is used for eyetracking calibration and needs to be run in the background of the experiment while collecting the data from Gazepoint. We have developed a Python script using Gazepoint SDK to send eyetracking and biometrics data (GSR, heart-rate, etc.) to **LSL**. This Python script can be accessed by downloading the folder *Gazepoint*(*Eyetracking+Biometrics*)-*LSL* from our GitHub: https://github.com/moeinrazavi/MultiModal_MultiSensory_Flanker_Task/tree/master/lib. The folder should be added to the experiment **lib** folder. The lines of code in Figure 29 are used to open Gazepoint Control. Then the code in Figure 30 is used to minimize the experiment screen to have access to Gazepoint Control for calibration. And finally, Figure 31 shows the code to check whether the Gazepoint Control is not closed by the user, and by running the *LSLGazepoint.py* as a subprocess, it starts sending data to **LSL**.

**Figure 29.**
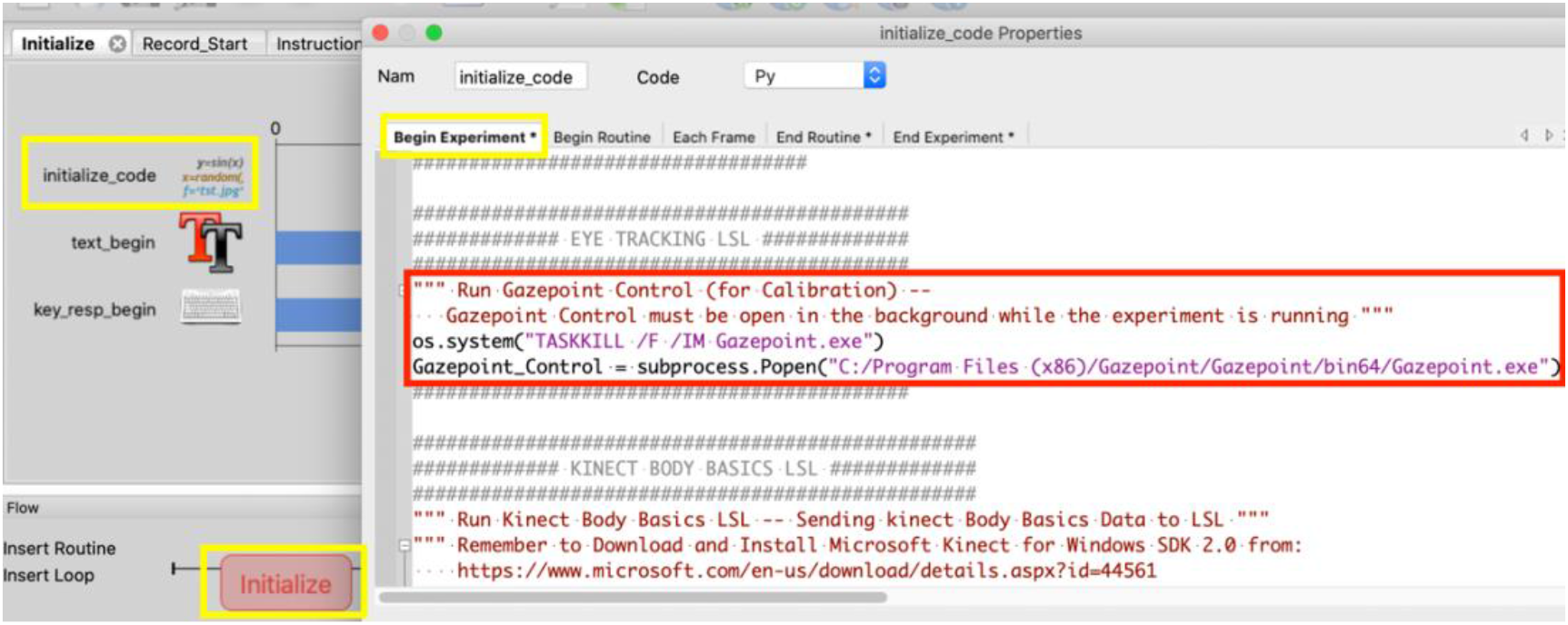
Script for running Gazepoint Control application.

**Figure 30.**
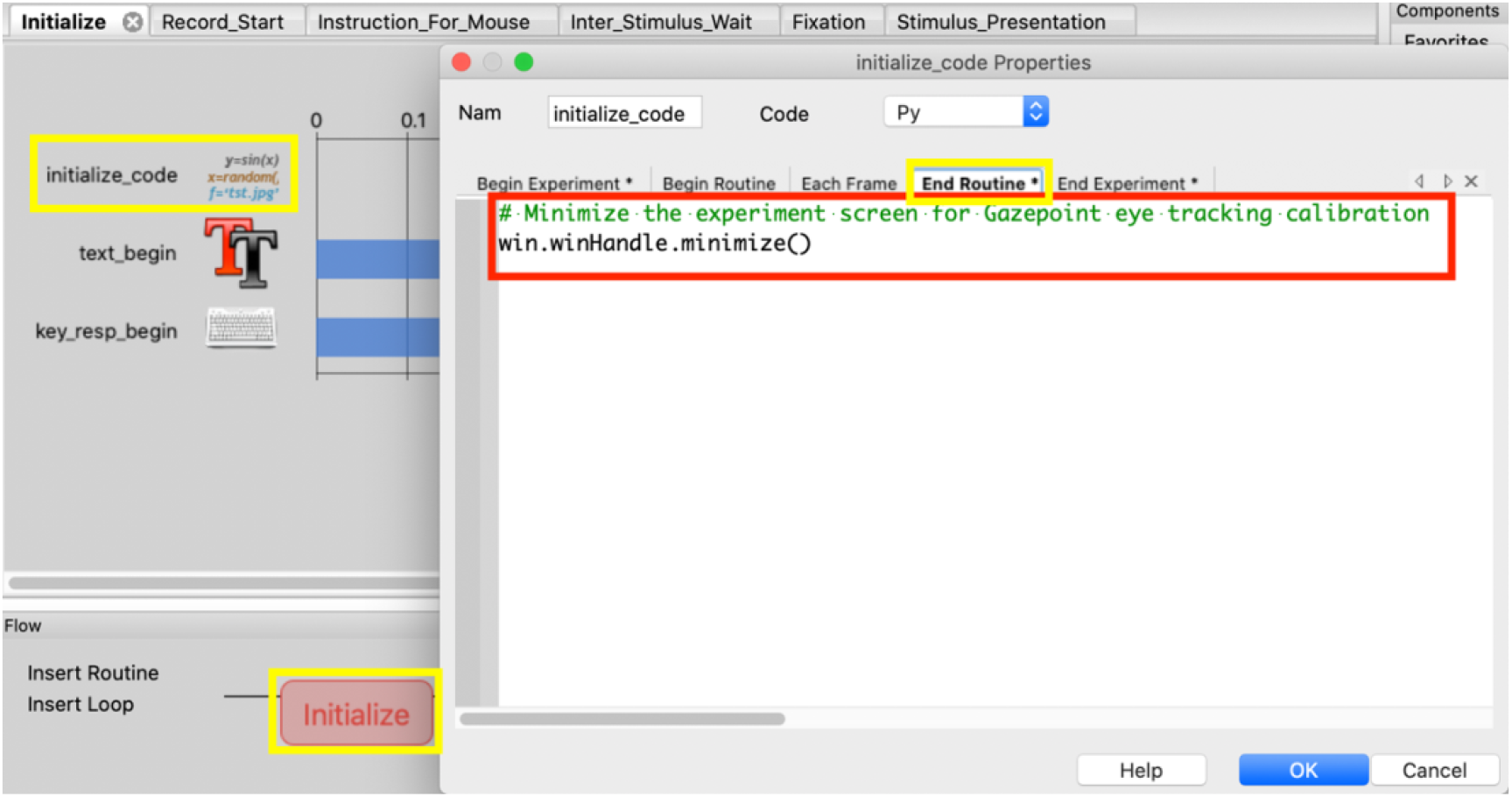
Script for minimizing the experiment screen to access the Gazepoint Control calibration

**Figure 31.**
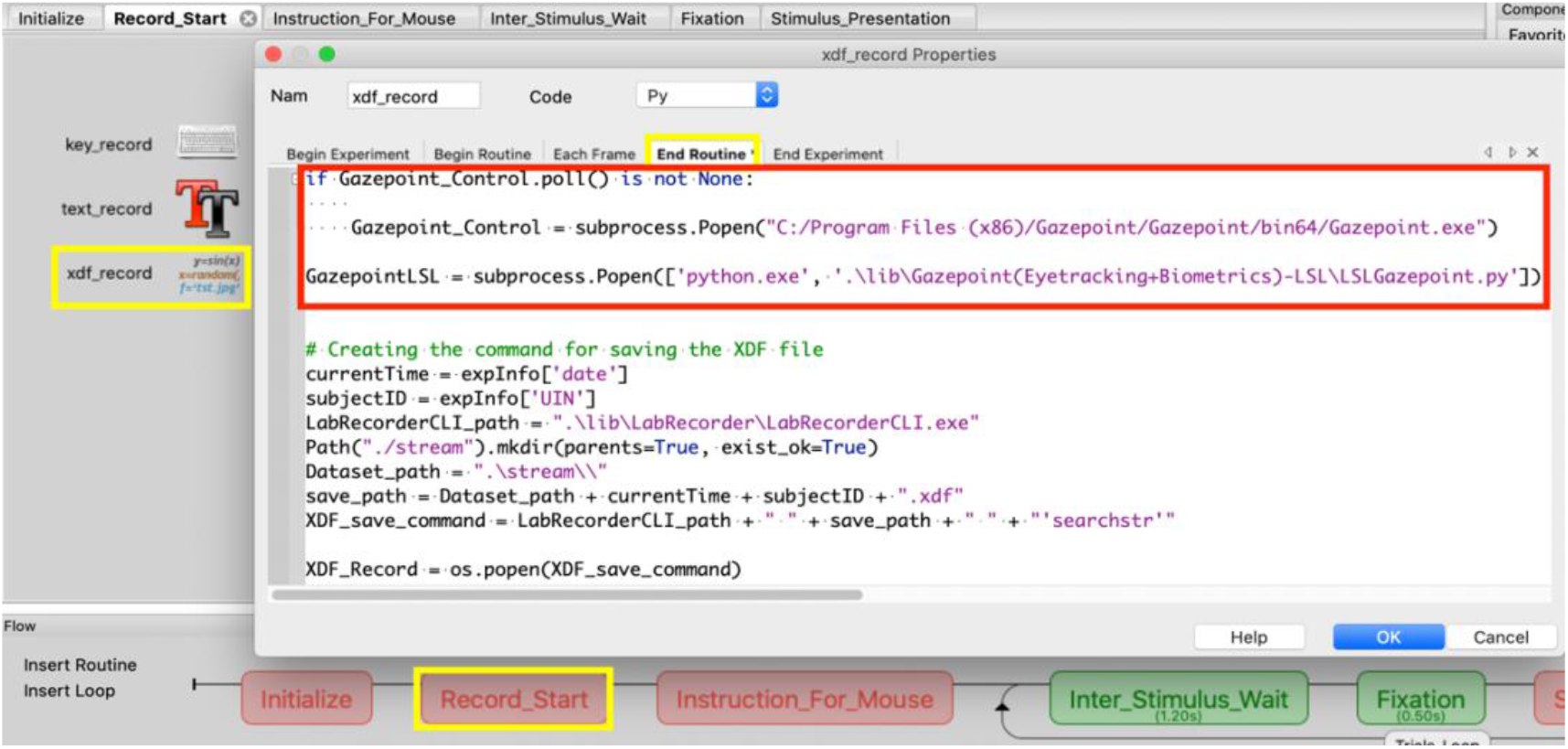
If Gazepoint Control closed by the user, run it again and send Gazepoint eyetracking (+ GSR and heartrate) to **LSL**

#### 4.3.5 Mouse Data + Flanker task markers + EEG + GSR + Eyetracking + Body Motion

Kinect for Windows v2 is used to record body motion. To do so, the Microsoft Kinect for Windows SDK 2.0 should be downloaded from https://www.microsoft.com/en-us/download/details.aspx?id=44561. We have developed a C++ application based on Kinect Body Basics SDK to send data from a connected Kinect device to **LSL**. This application can be accessed by downloading the folder *Kinect-BodyBasics-LSL* from our GitHub: https://github.com/moeinrazavi/Kinect-BodyBasics-LSL and copying this folder in the experiment *lib* folder. Then, the code shown in Figure 32 should be added to the **Begin Experiment** tab of **initialize_code** in the **Initialize** routine.

**Figure 32.**
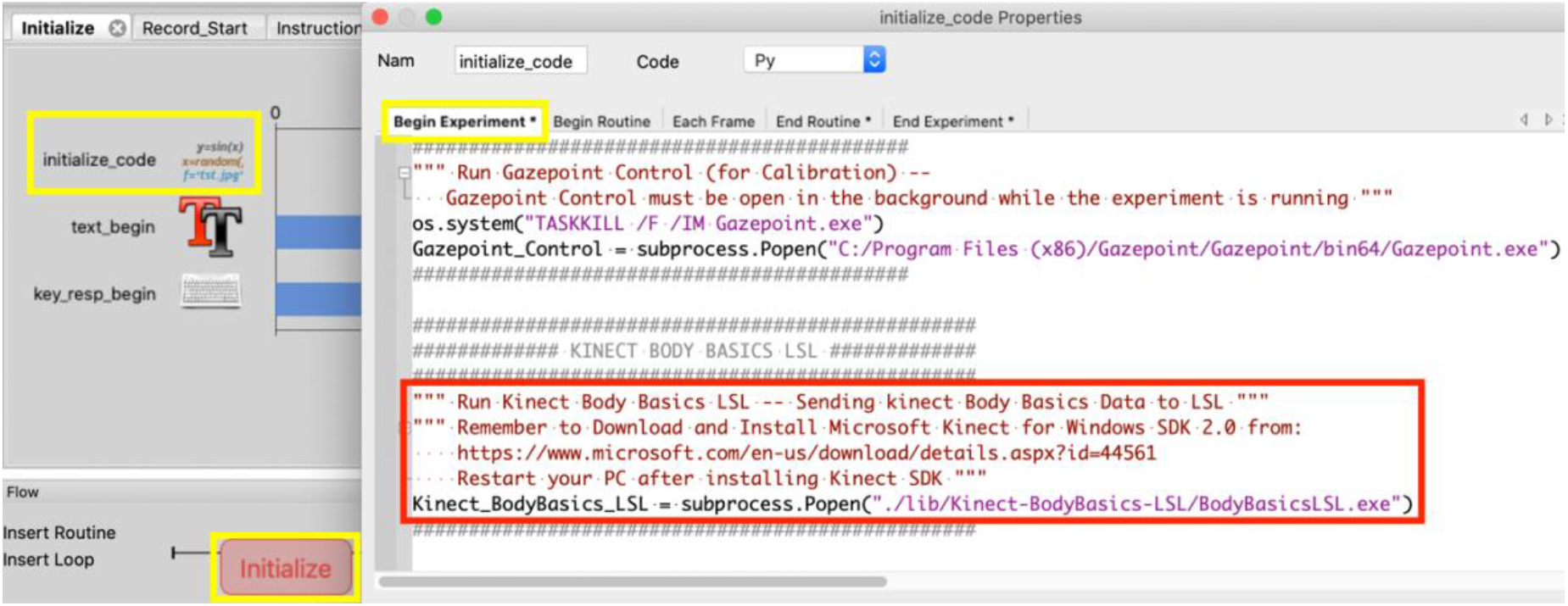
Running the application for sending Kinect body motion data to **LSL**

At the end of experiment, in order to stop recording, the lines of code shown in Figure 33 can be added to the **End Experiment** tab.

**Figure 33.**
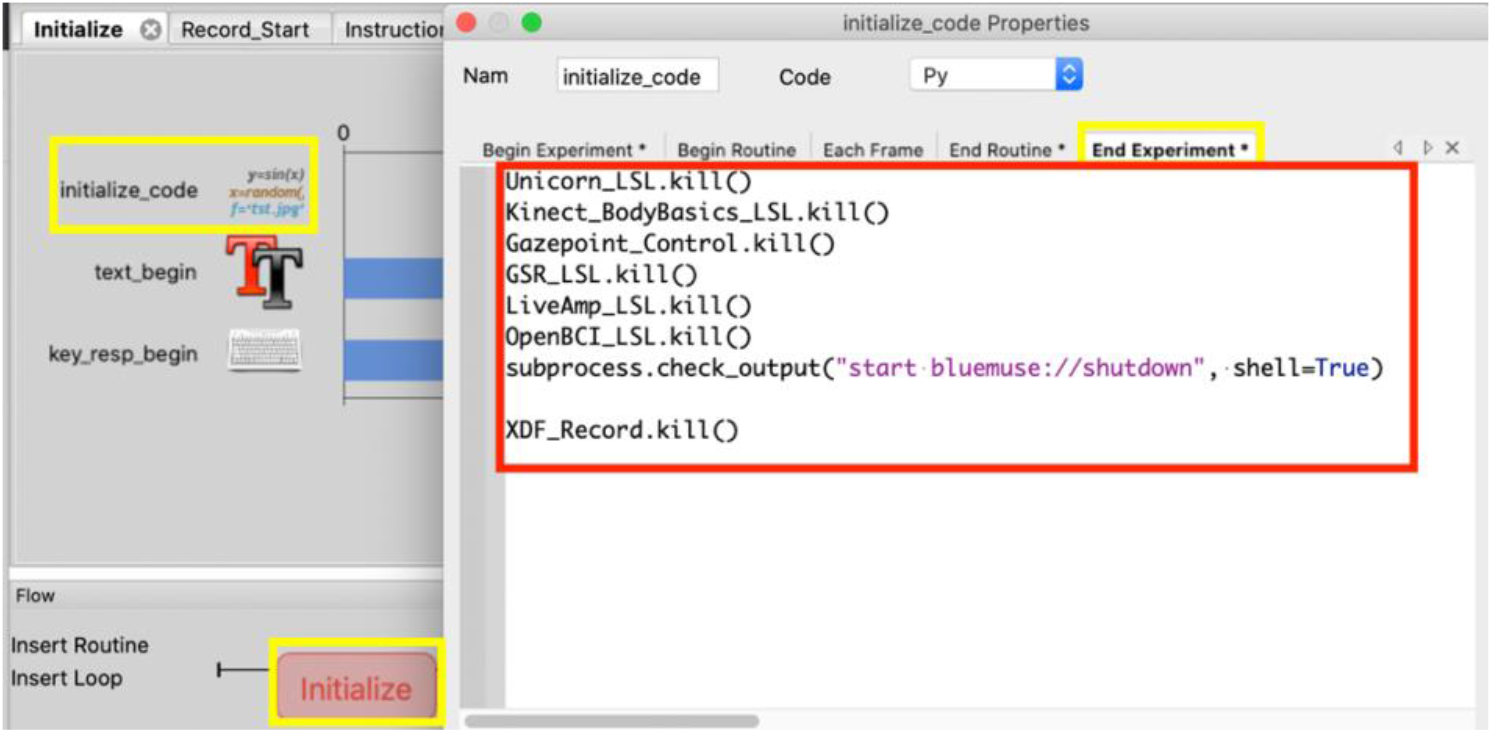
Script for stop recording the data from all the devices

## 5. Building a Multisensory Virtual Reality (VR) Experiment (Unity)

In this section, first it is shown how to create simple interactable game objects in a **VR** environment using **Unity**. Then, a tutorial is provided on how to synchronize **VR** environment data (e.g., objects positions and orientations), user data (i.e., marker for pressing HTC VIVE controller trigger, positions of player and controllers), and data from multiple devices such as EEG (g.tec Unicorn, Muse, Neurosky Mindwave, BrainProducts LiveAmp, OpenBCI Cyton and OpenBCI Cyton + Daisy), GSR (e-Health Sensor Platform v2.0 for Arduino), Eyetracking (HTC VIVE Pro with Tobii Eyetracking), and Body Motion (Kinect) together by **LSL**. Finally, it is explained how to embed **LabRecorder** in **Unity** to save data automatically on disk.

### 5. 1. Software and Plugin Installation

In order to design a **VR** environment, **Unity** (https://unity3d.com/get-unity/download) and **Steam** (https://store.steampowered.com/about/) need to be installed which are available to download for free.

### 5. 2. Create simple interactable objects in **VR** environment

After installing **Unity** and Steam, in order to create a **VR** environment in Unity, a new 3D project should be created from Projects part in Unity Hub (Figure 34).

**Figure 34.**
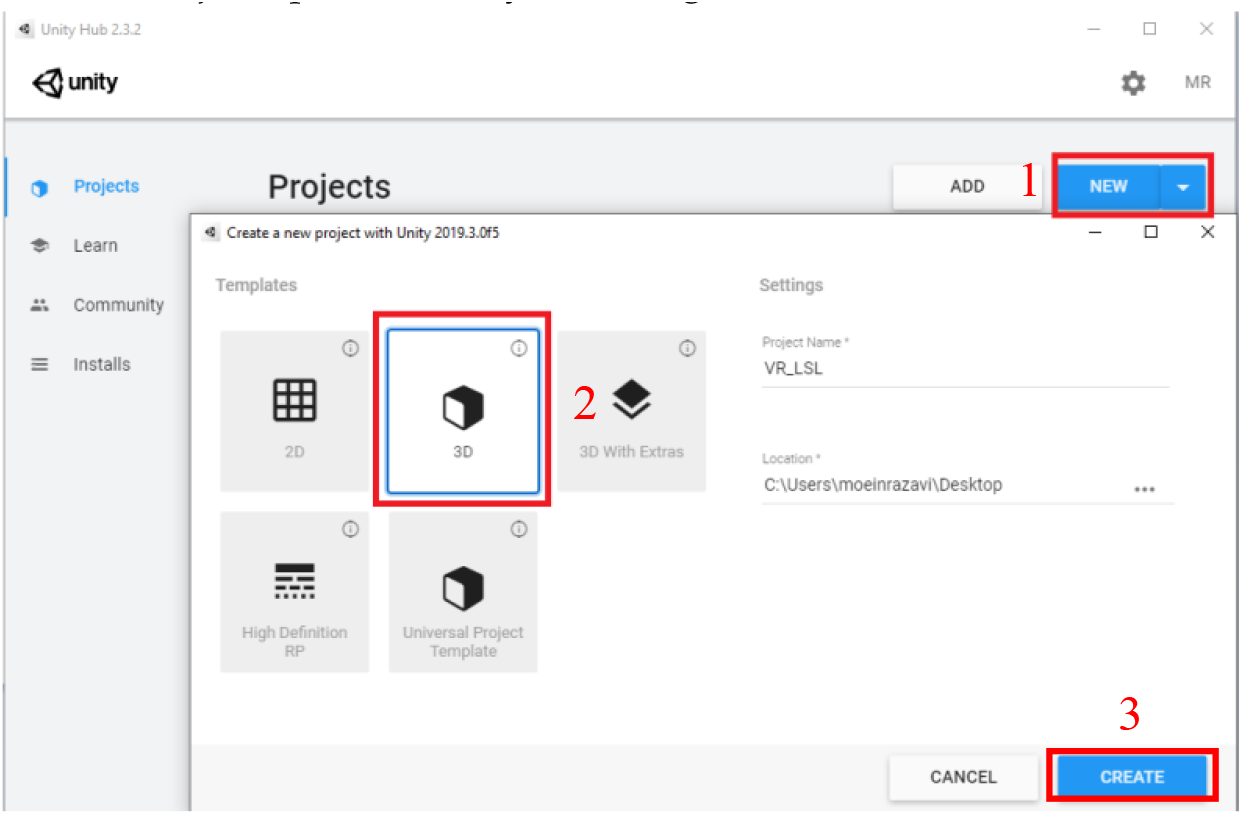
Create a new 3D project in **Unity**

After creating the project, there is a Main Camera object in **Unity** SampleScene which is fixed to the environment and useless for **VR** purposes (Figure 35). It will be shown later how to attach a camera object to the player that can move with it.

**Figure 35.**
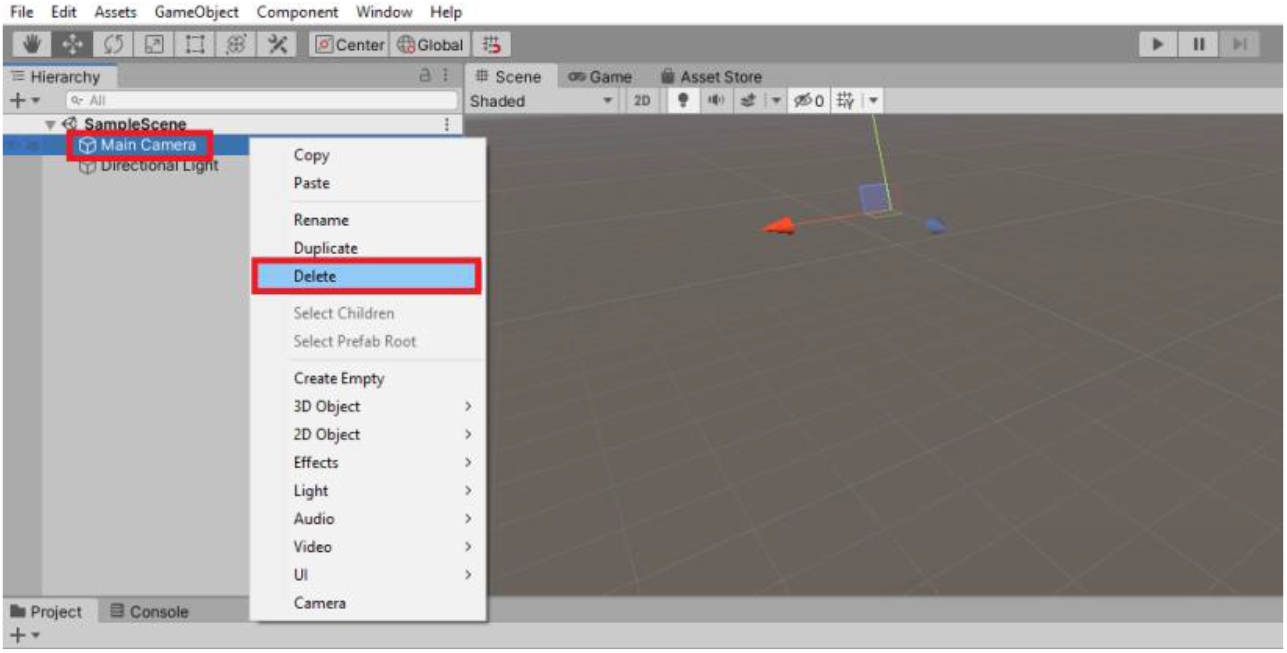
Delete Main Camera object from the Scene

To utilize **VR** properties and functions, SteamVR Plugin should be imported to the project. SteamVR Plugin can be downloaded and imported from Asset Store in **Unity**. In the pop-up windows that appear after importing SteamVR Plugin, “import” (Figure 36.a) and “Accept All” (Figure 36.b) should be selected, respectively.

**Figure 36.**
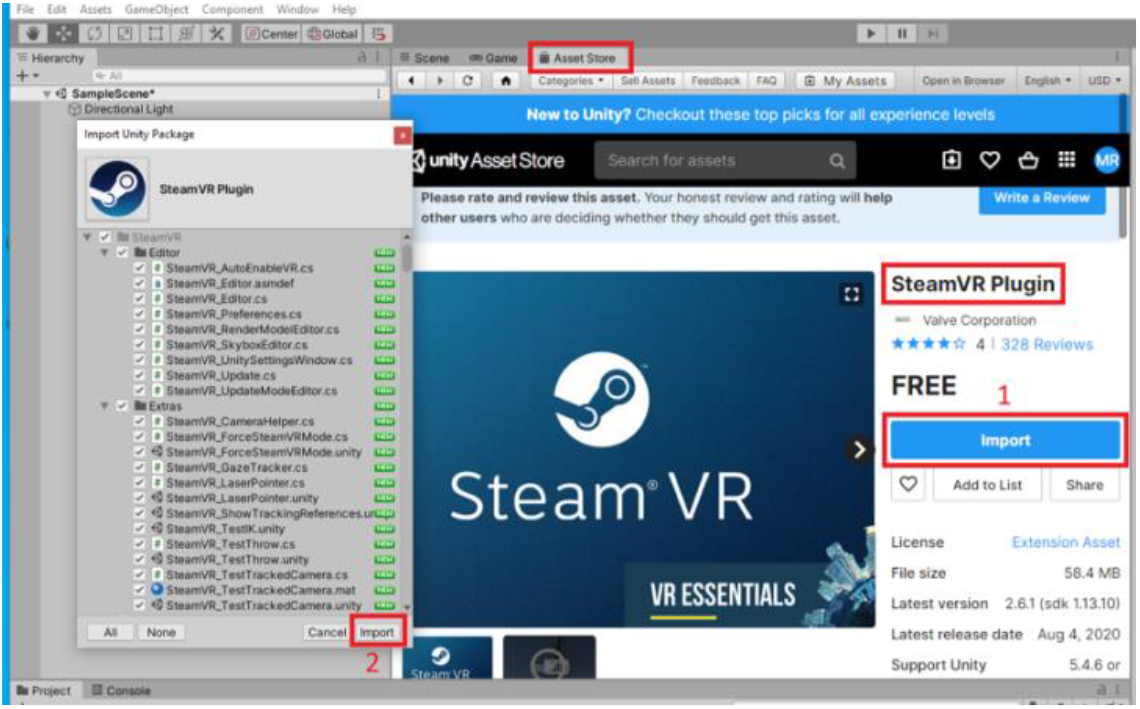
**a.** Import SteamVR Plugin

**Figure 36.**
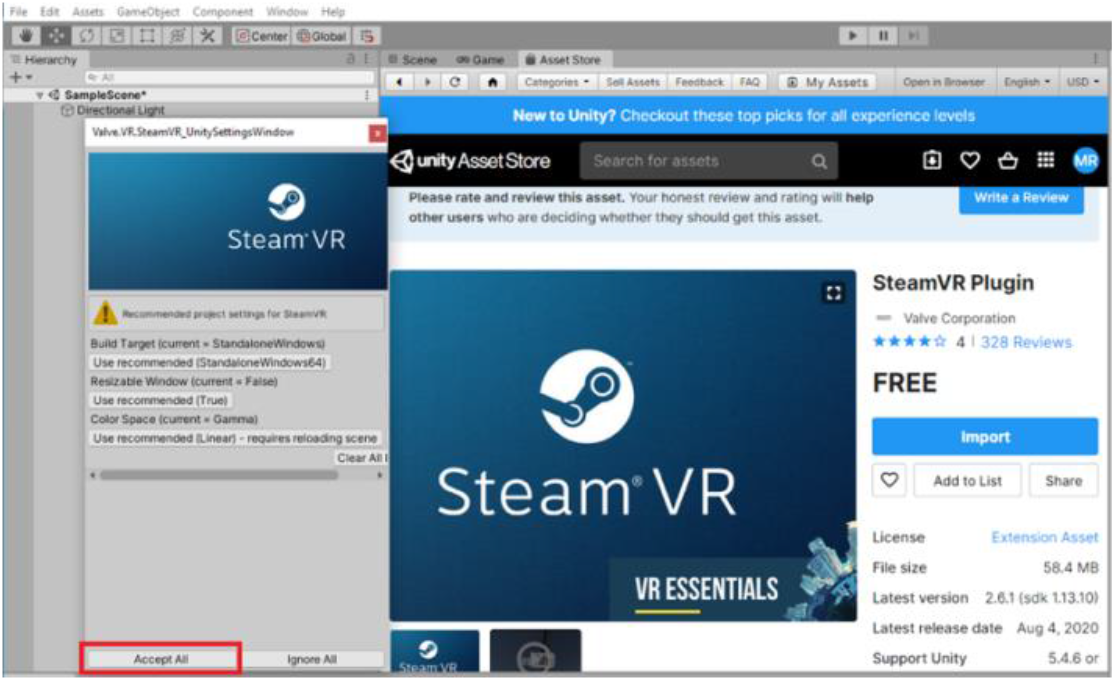
**b.** Import SteamVR Plugin

Next, simple objects (e.g., Plane and Cube) are added to the Scene (Figure 37). To make the objects interactable with the user, first the **Interactable** and then **Throwable** features should be added to the **Cube** object (Figure 38). By doing this, the user can move and throw the Cube object.

**Figure 37.**
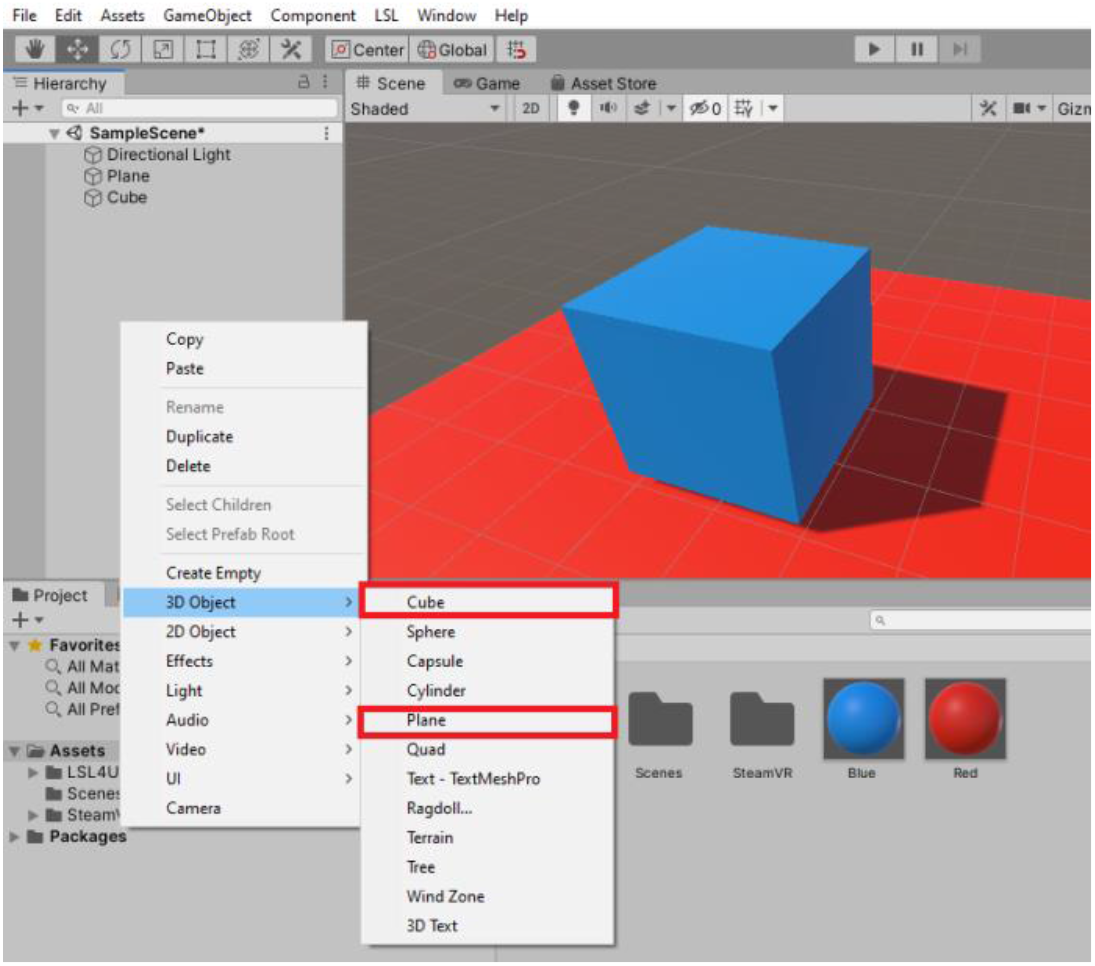
Add Plane and Cube objects to the Scene

**Figure 38.**
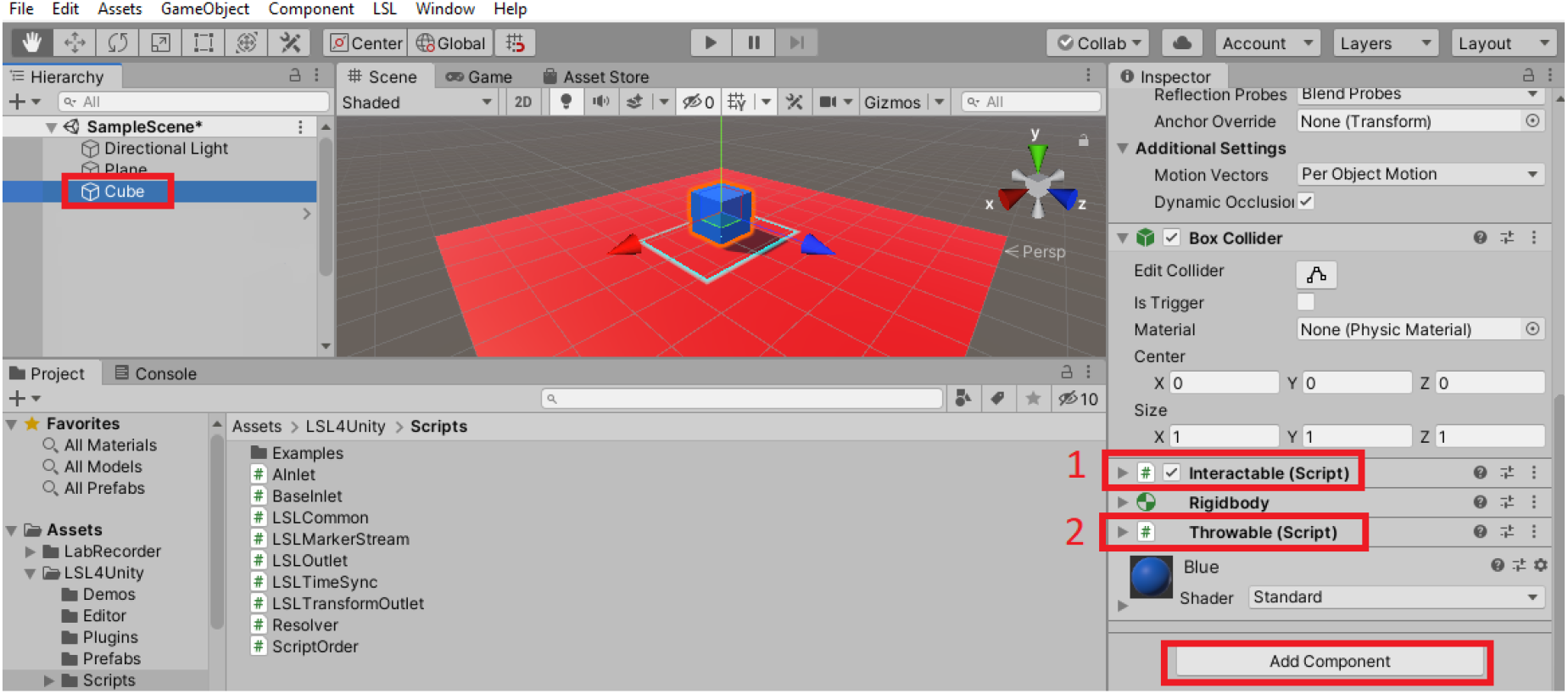
Add **Interactable** and then **Throwable** components to the **Cube**

Then a Player object is added to the Scene by dragging it to the Scene (Figure 39). The Player object will define the position of the user in the **VR** environment. After that, [CameraRig] object is added to the Player by dragging it to the Player (Figure 40). This will attach the **VR** camera and controllers (here HTC VIVE Pro camera and controllers) to the Player object.

**Figure 39.**
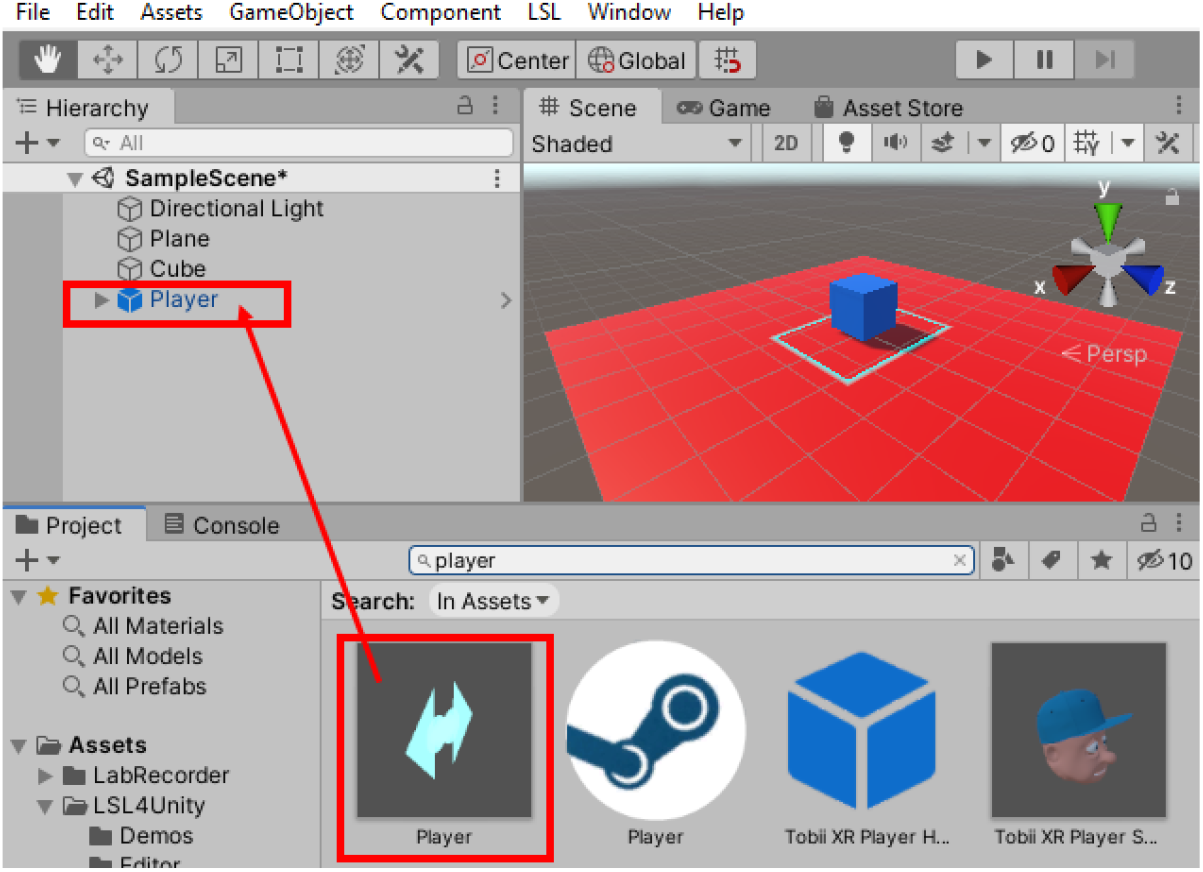
Add Player object to the Scene

**Figure 40.**
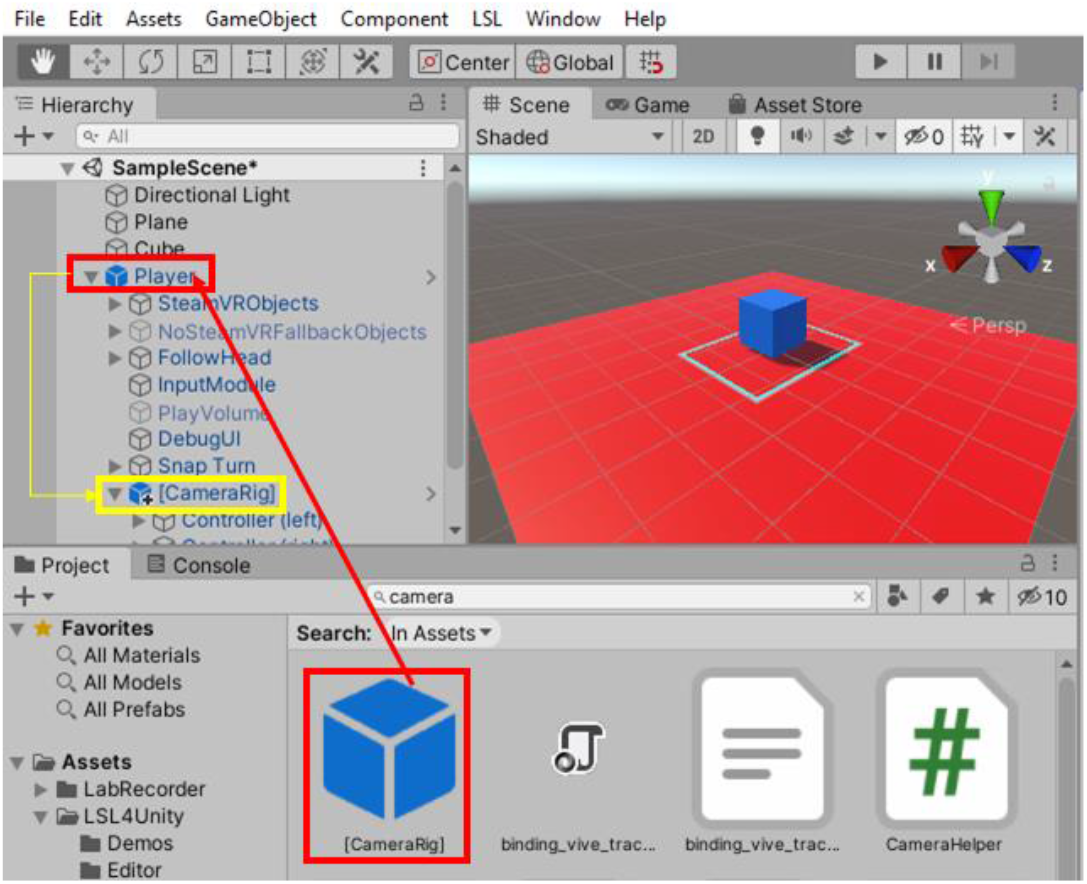
Add [CameraRig] object to the Player

In the following, it is shown how to synchronize the data from the **VR** environment, user and different devices using **LSL**.

### 5. 3. Synchronize data from **VR**, user and different devices by **LSL**

In this step, to use **LSL** features, **LSL4Unity** plugin should be added to the *Assets* folder in the main project folder (here **VR_LSL**). In addition, to send HTC VIVE eyetracking and controller trigger press data to **LSL**, we developed two C# scripts which should also be added to the *Assets* folder (Figure 41). The **LSL4Unity** plugin and these scripts can be accessed from our GitHub: https://github.com/moeinrazavi/VR_LSL.

**Figure 41.**
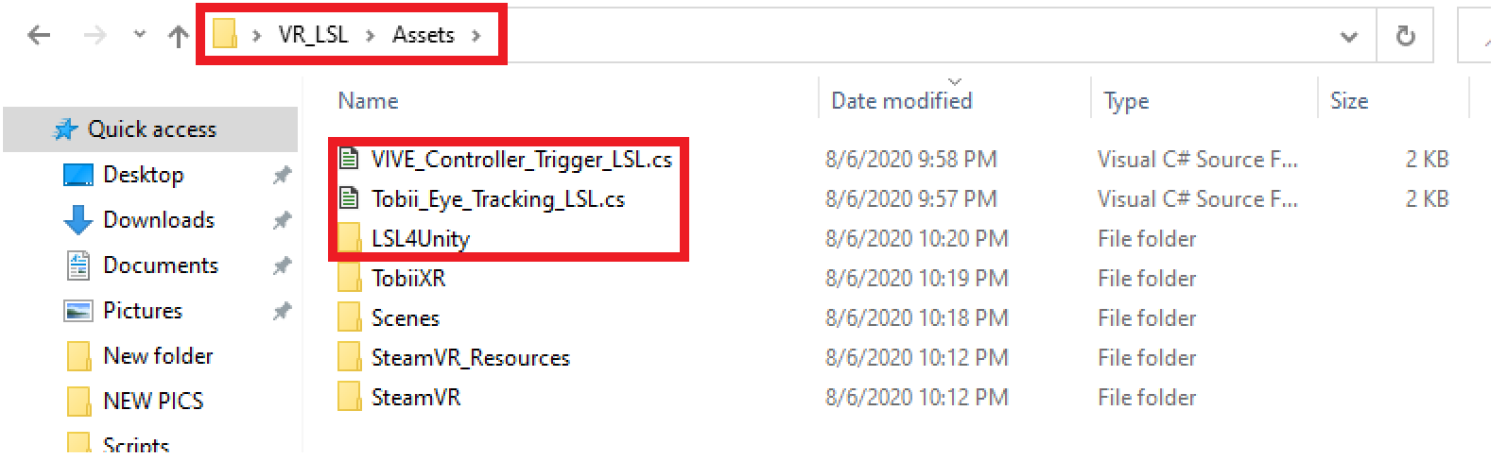
Add **LSL4Unity** plugin and developed scripts for sending HTC VIVE eyetracking and controller trigger press data to **LSL** in the project’s *Assets* folder.

#### 5.3.1 Send **VR** objects data to **LSL**

Each object in **Unity** has a component named Transform which defines the Position and Rotation of that object in the environment (Figure 42). The LSLTransformOutlet function in **LSL4Unity** plugin can be attached to each object and send data from its Transform component to **LSL**. Figure 43 shows how to add this function to the Cube object by dragging the function to the object. The Sample Source field of this function should be set to the object that its Transform data needs to be sent to **LSL** (Figure 43). The LSLTransformOutlet function provides up to 3 types of data to be sent to **LSL**. The first type contains 4 channels that send the rotation data in the Quaternion system (angles with x,y,z and w axes); the next one has 3 channels that are associated with the rotation data in the Euler system (angles with x,y and z axes). The last one includes 3 channels that send the position data (x, y and z coordinates) to **LSL** (Figure 44).

**Figure 42.**
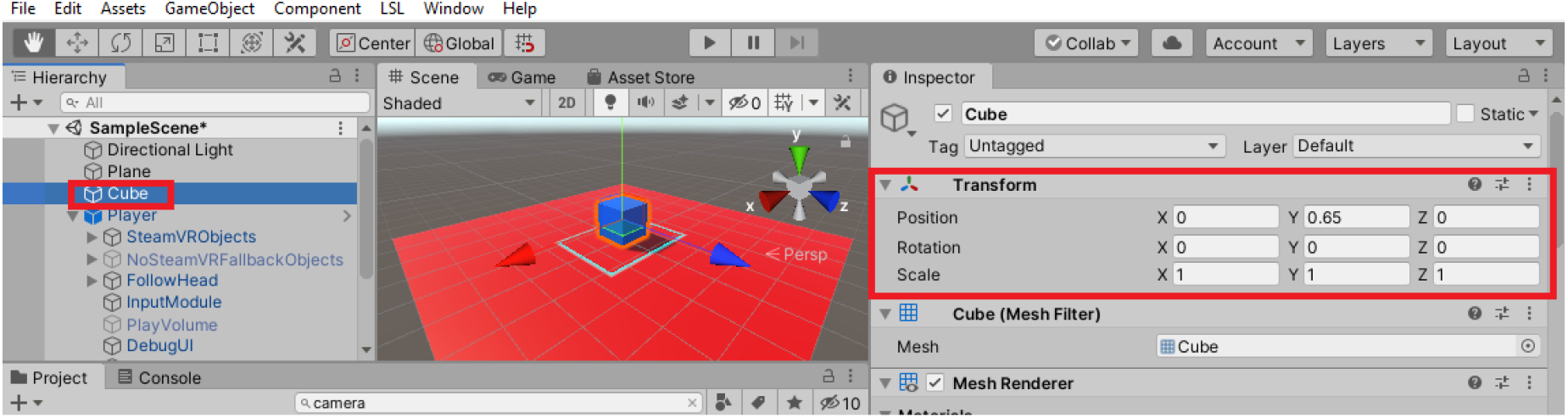
Transform component of an object

**Figure 43.**
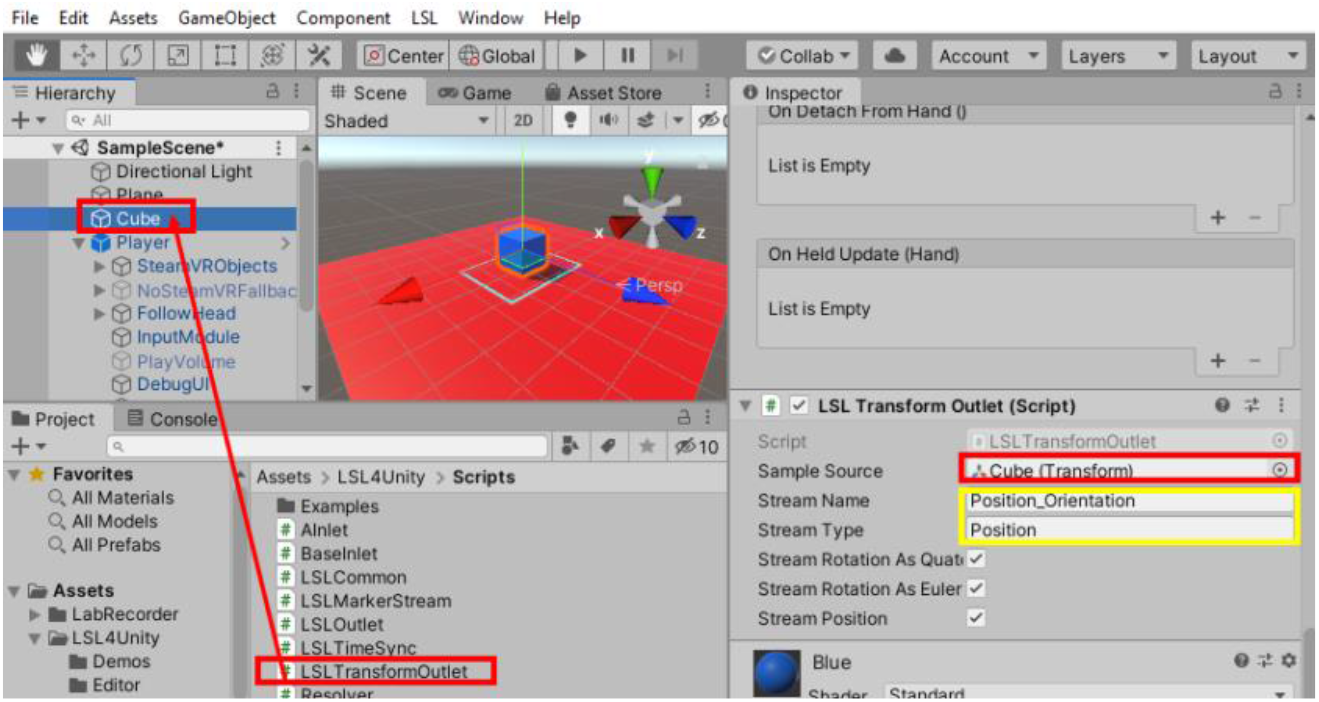
Add LSLTransformOutlet function to the Cube object

**Figure 44.**
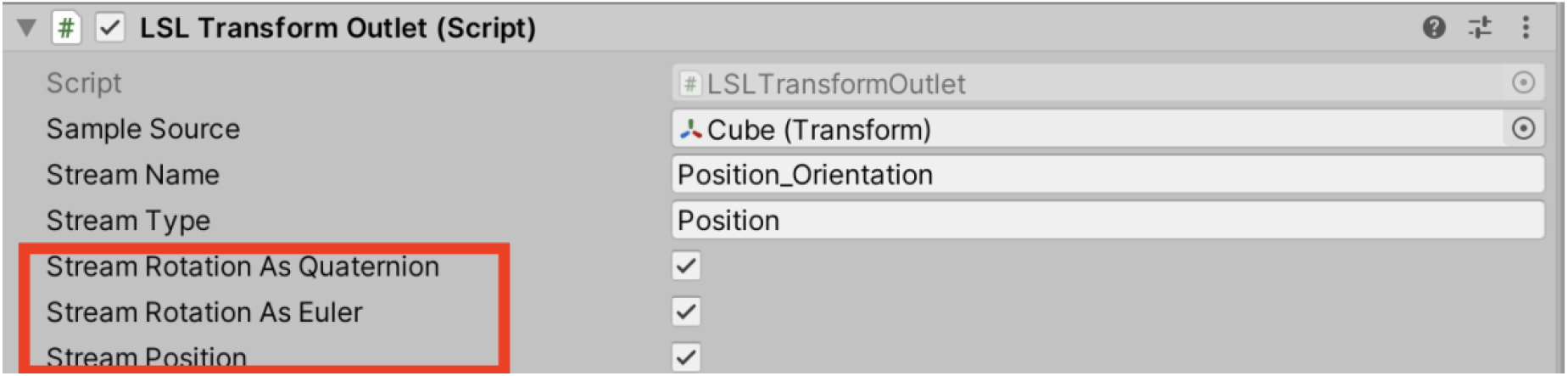
LSLTransformOutlet data streaming choices

#### 5.3.2 Send **VR** objects data + user data to **LSL**

Here it is shown how to send the camera rotation, camera position coordinates and controller trigger press markers to **LSL**. The camera is attached to the Player object.

First, to send the camera rotation and position data to **LSL**, similar to the previous subsection, the LSLTransformOutlet function is attached to Camera object under the Player object and Camera is chosen as the Sample Source of this function (Figure 45). Similar to camera, the position and rotation of the controllers can be sent to LSL by attaching LSLTransformOutlet function to the Transform component of each controller (not shown here for brevity).

**Figure 45.**
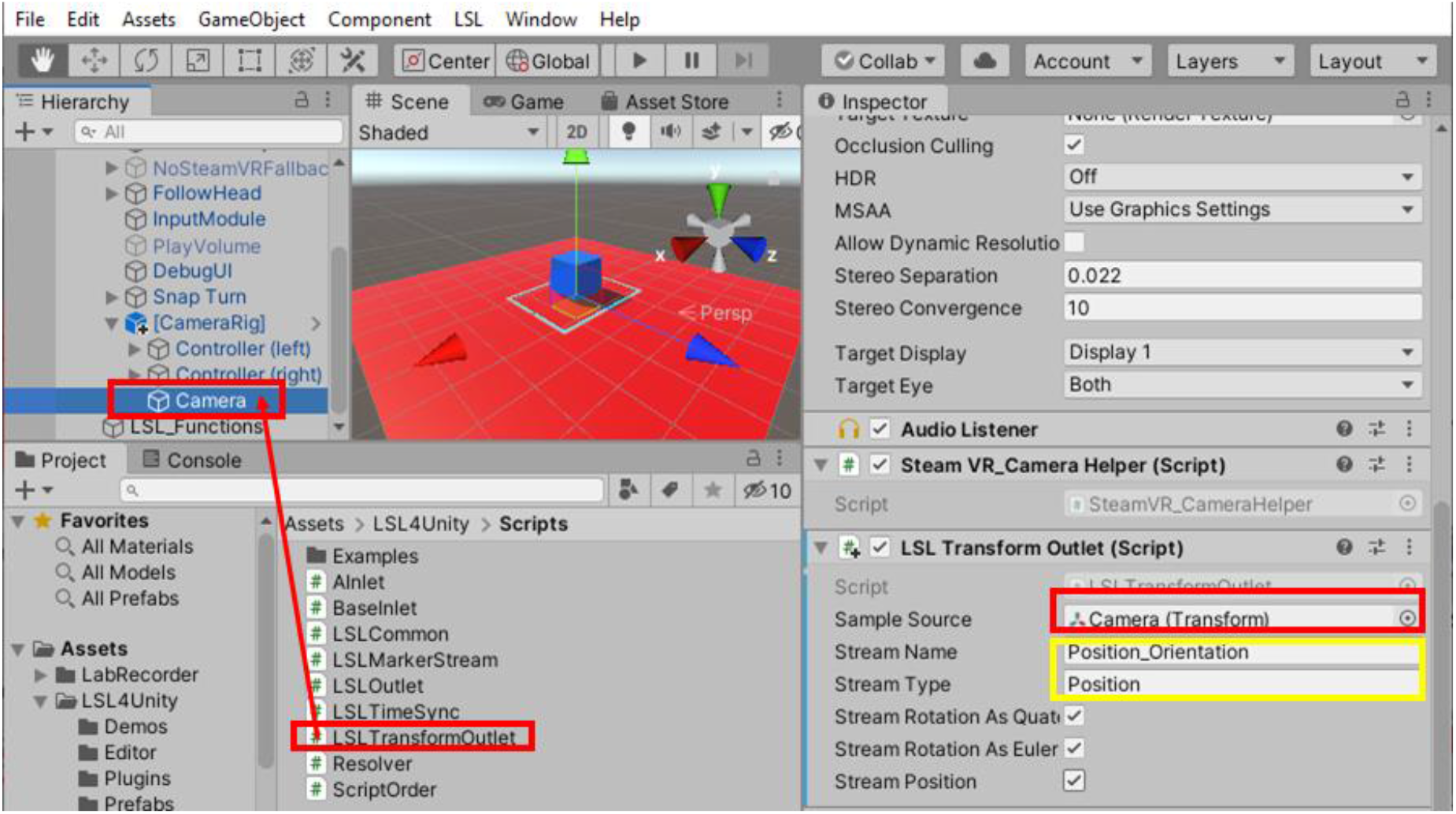
Add LSLTransformOutlet function to the Camera object

Then, to have access to controllers, SteamVR Input need to be generated by clicking on SteamVR Input in Window tab. On every pop-up window that appears, **Yes** and finally **Save and generate** buttons should be selected (Figure 46). The SteamVR Input makes controller button actions accessible.

**Figure 46.**
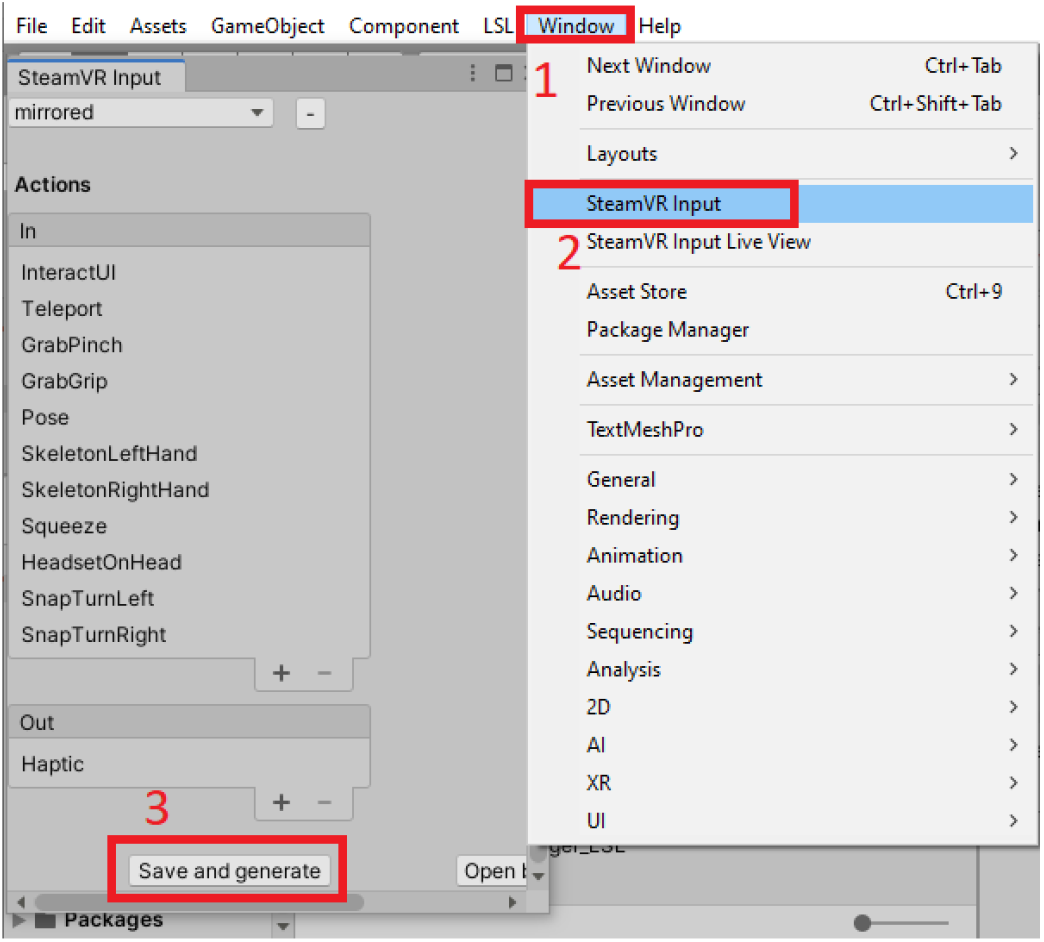
Generate SteamVR Input

#### 5.3.3 Send **VR** objects data + user data + data from different devices to **LSL**

In order to send data from controllers, eyetracking and other devices, an empty GameObject is created and added to LSL_Functions (Figure 47).

**Figure 47.**
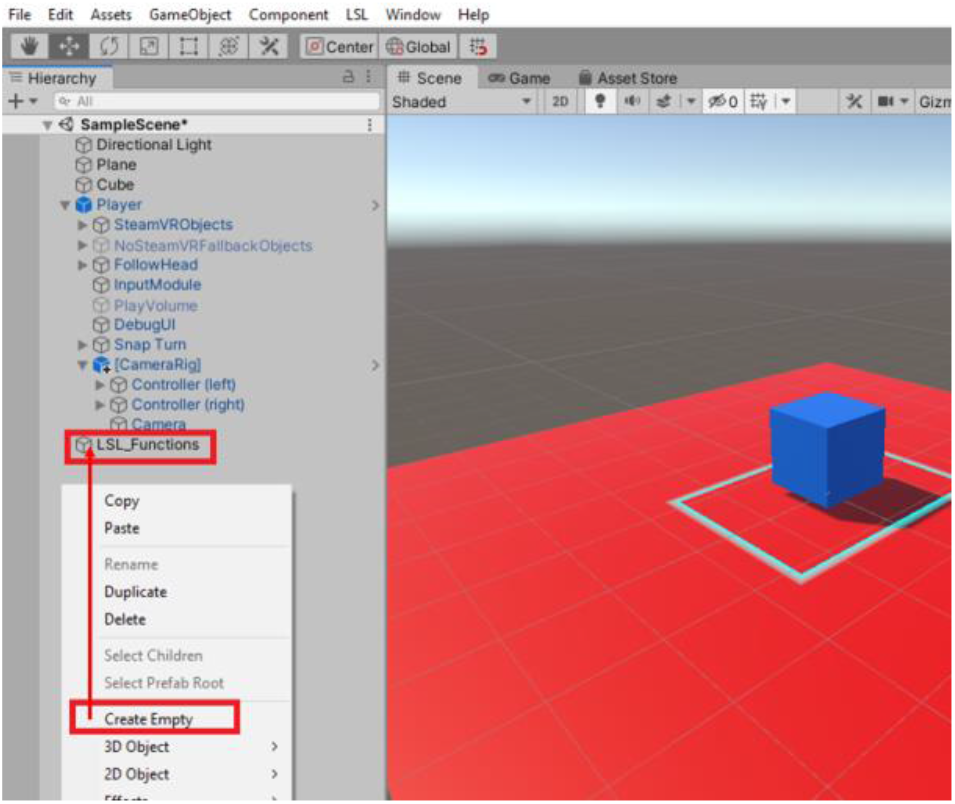
Create an empty GameObject (here renamed to LSL_Functions)

We have developed C# scripts to send data from different devices to **LSL**. In the following, it is shown how to add these scripts to LSL_Functions object, step by step. A brief explanation for some of the scripts is also provided.

##### • Sending controller trigger press marker to LSL

To send the controller trigger press markers to **LSL**, the **Steam VR_Activate Action Set On Load** function needs to be added to LSL_Functions object. This function allows access to controller button actions. The Action Set in this function must be set to \actions\default. Then the **VIVE_Controller_Trigger_LSL** script that we have developed in C# and the **LSL Marker Stream** function need to be added to LSL_Functions. The **VIVE_Controller_Trigger_LSL** function uses **LSL Marker Stream** to send controller trigger press data to **LSL** (Figure 48). In **VIVE_Controller_Trigger_LSL** panel, the Trigger_Data should be set on \actions\default\in\GrabPinch; this will assign the action of sending data to **LSL** to the controller trigger. Choosing Right Hand or Left Hand for the **Hand Type,** will determine whether to send data from the right-hand or the left-hand controller to **LSL**.

**Figure 48.**
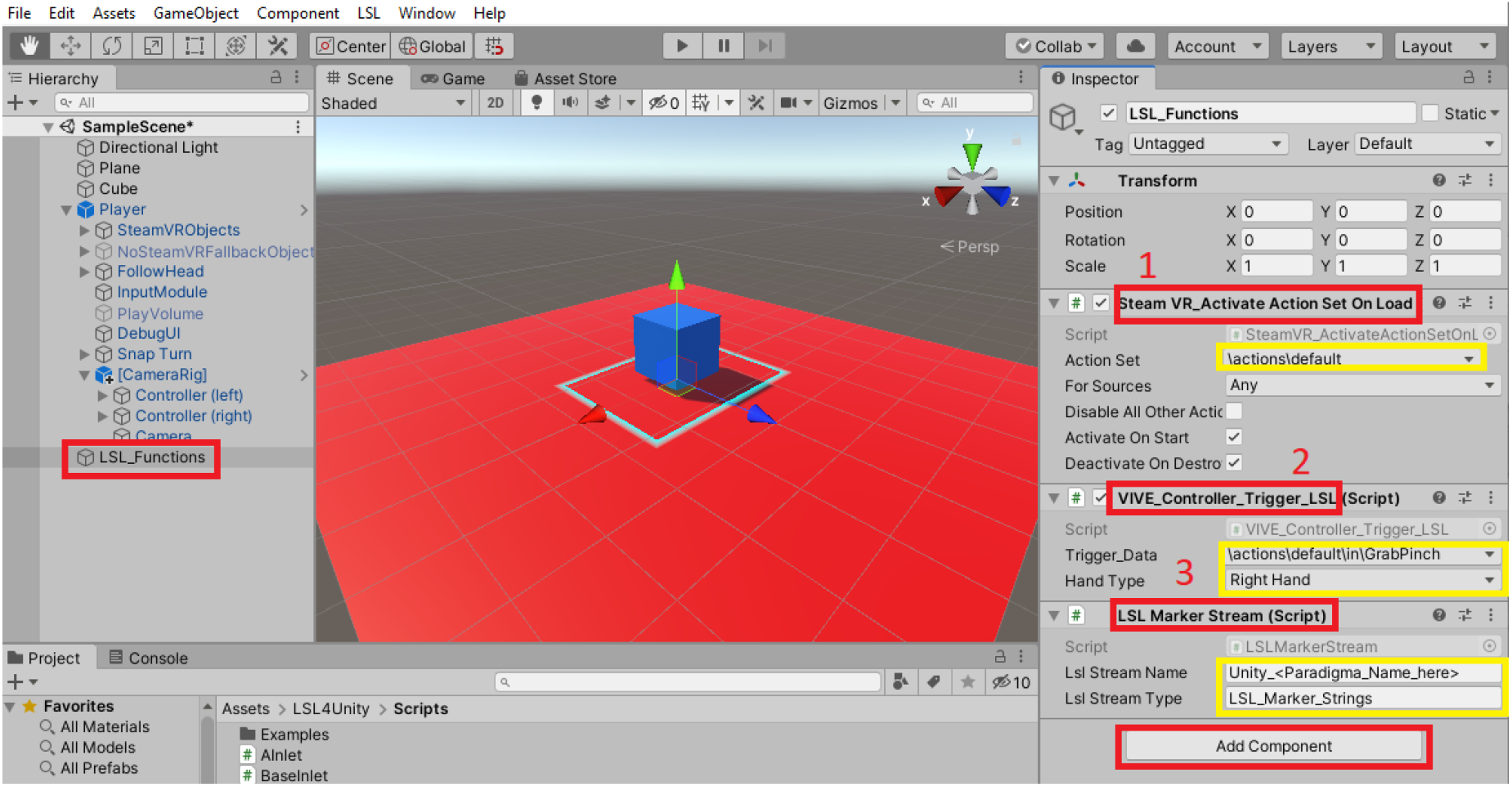
Add the scripts to send controller trigger press data to **LSL**

##### • Sending Eyetracking data to LSL

In order to send the Eyetracking (HTC VIVE Pro with Tobii Eyetracking) data to **LSL**, first, Tobii XR SDK for **Unity** needs to be downloaded from https://vr.tobii.com/sdk/downloads/. Then, while the project is open, opening the downloaded file will start importing the required libraries for Tobii eyetracking in **Unity**.

Then the **Tobii_Eye_Tracking_LSL** function that we developed with C# should be added to the **LSL Functions** object (Figure 49).

**Figure 49.**
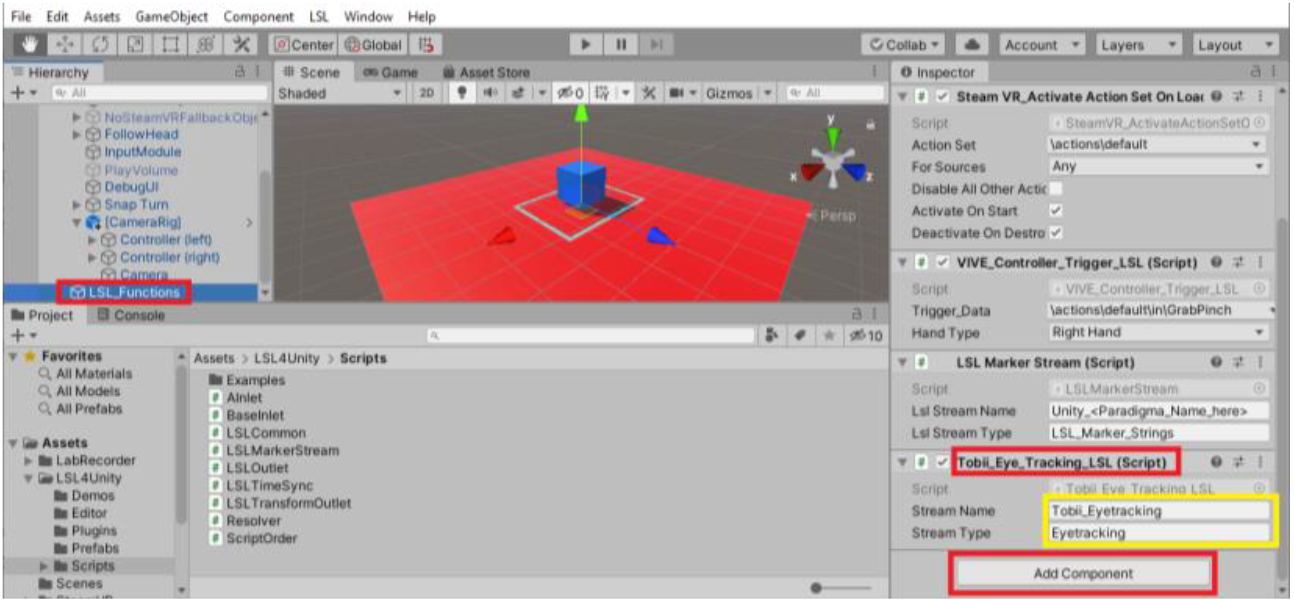
Add the script to send eyetracking data to **LSL**

##### • Send data from other devices to LSL

Here it is shown how to send data from other devices (EEG (g.tec Unicorn, Muse, Neurosky Mindwave, BrainProducts LiveAmp, OpenBCI Cyton and OpenBCI Cyton + Daisy), GSR (e-Health Sensor Platform v2.0 for Arduino), and Body Motion (Kinect)) to **LSL**. To do that, first, the required folders should be downloaded from our GitHub and copied in the *Assets* folder (Figure 50). Also, the **Multiple_Devices_LSL** script that we developed in C# needs to be copied it in the *Assets* folder (Figure 51). This script uses Windows Command Prompt to automatically send data from the devices mentioned above to **LSL** (Figure 52).

**Figure 50.**
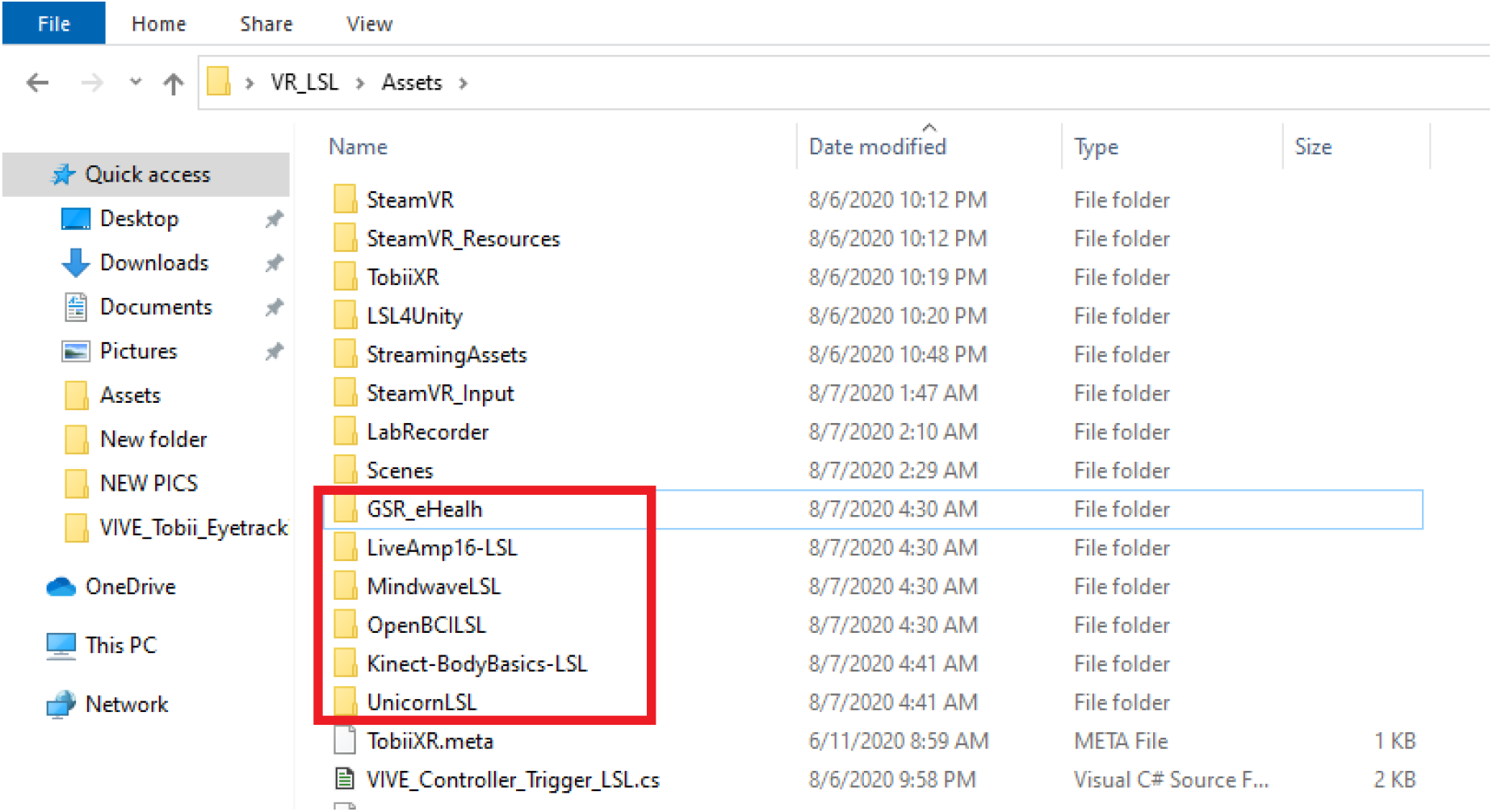
Add files associated with different devices to the *Assets* folder

**Figure 51.**
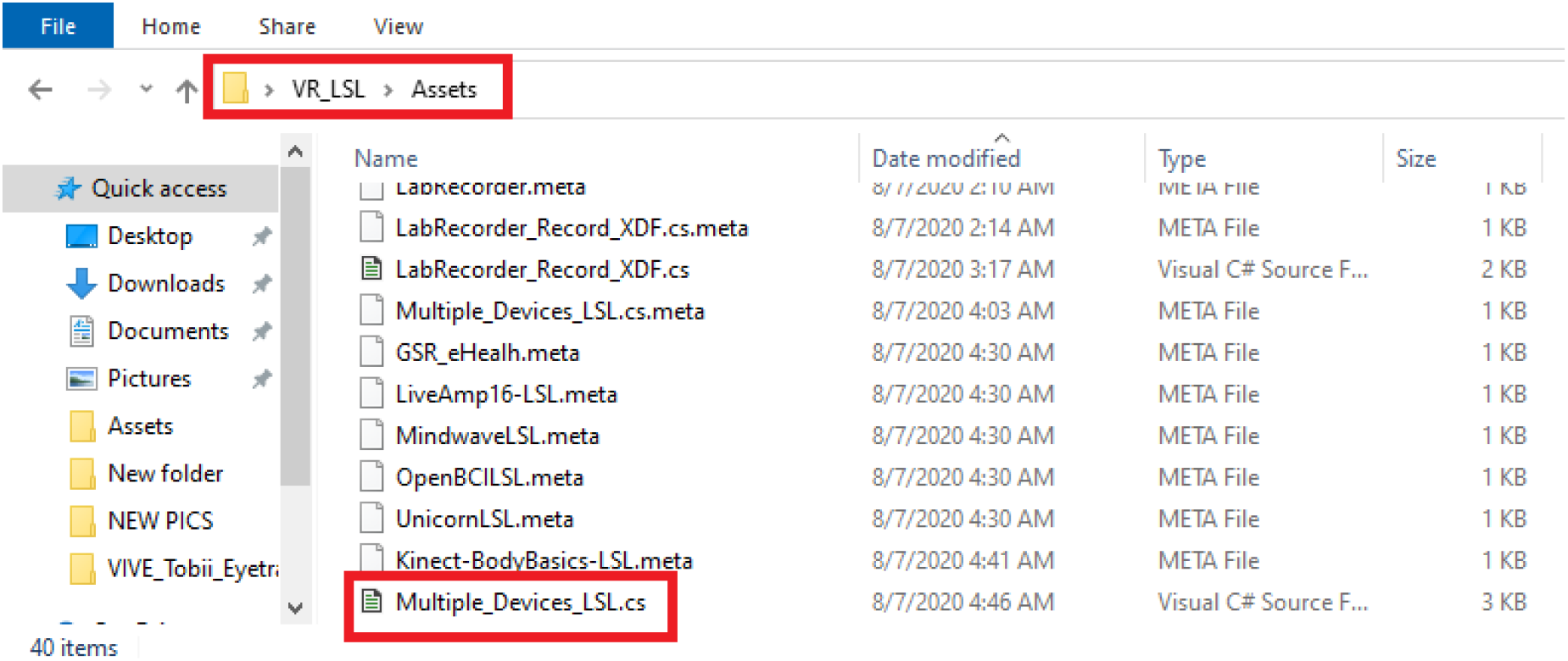
Add the **Multiple_Devices_LSL** script to *Assets* folder

**Figure 52.**
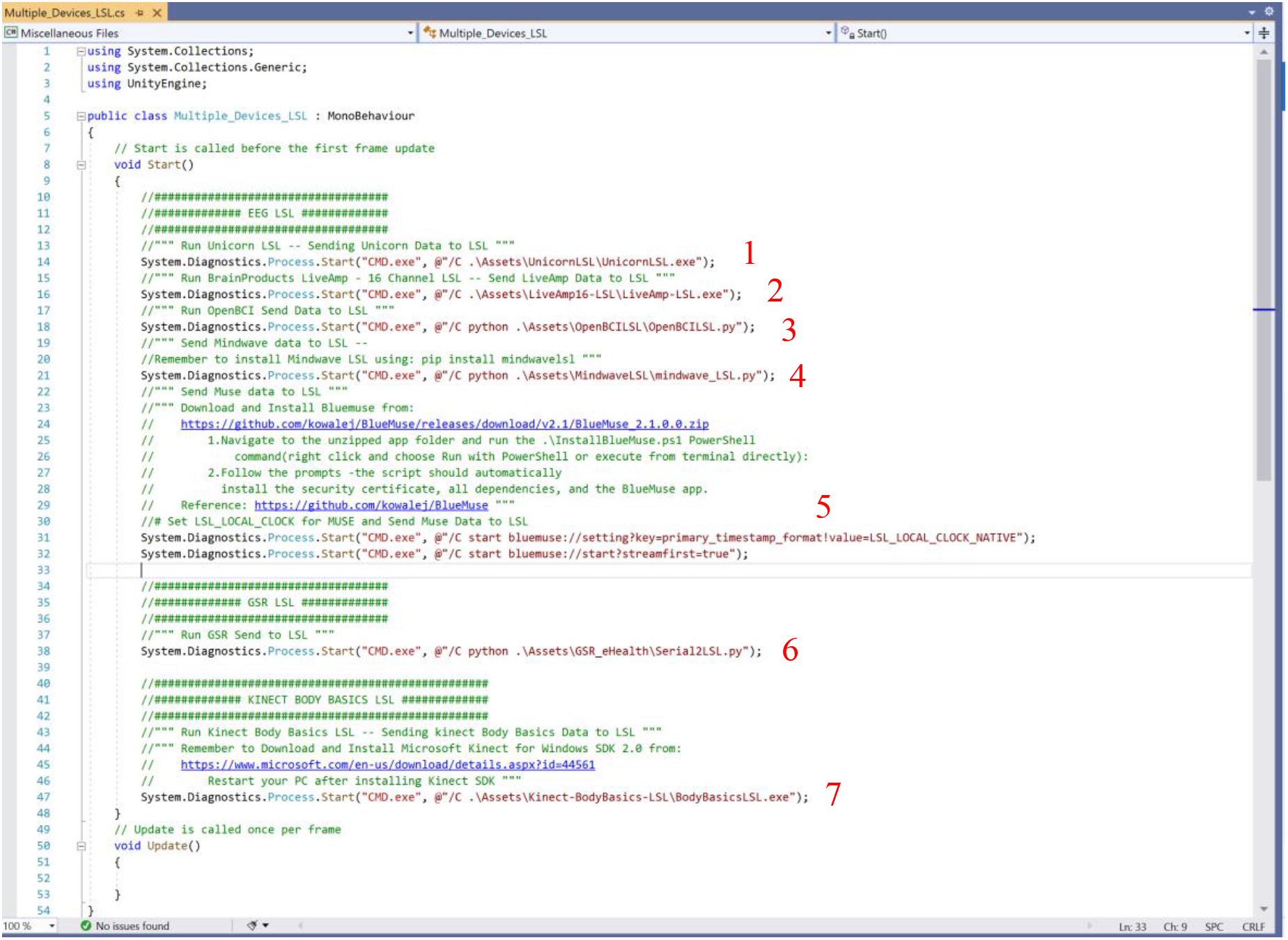
**Multiple_Devices_LSL** script: Send data from 1) g.tec Unicorn, 2) BrainProducts LiveAmp, 3) OpenBCI, 4) NeuroSky Mindwave, 5) Muse, 6) eHealth v2 GSR sensor and 7) Kinect to **LSL**

In Figure 53, it is shown how to add **Multiple_Devices_LSL** script to **LSL_Functions** object in **Unity**.

**Figure 53.**
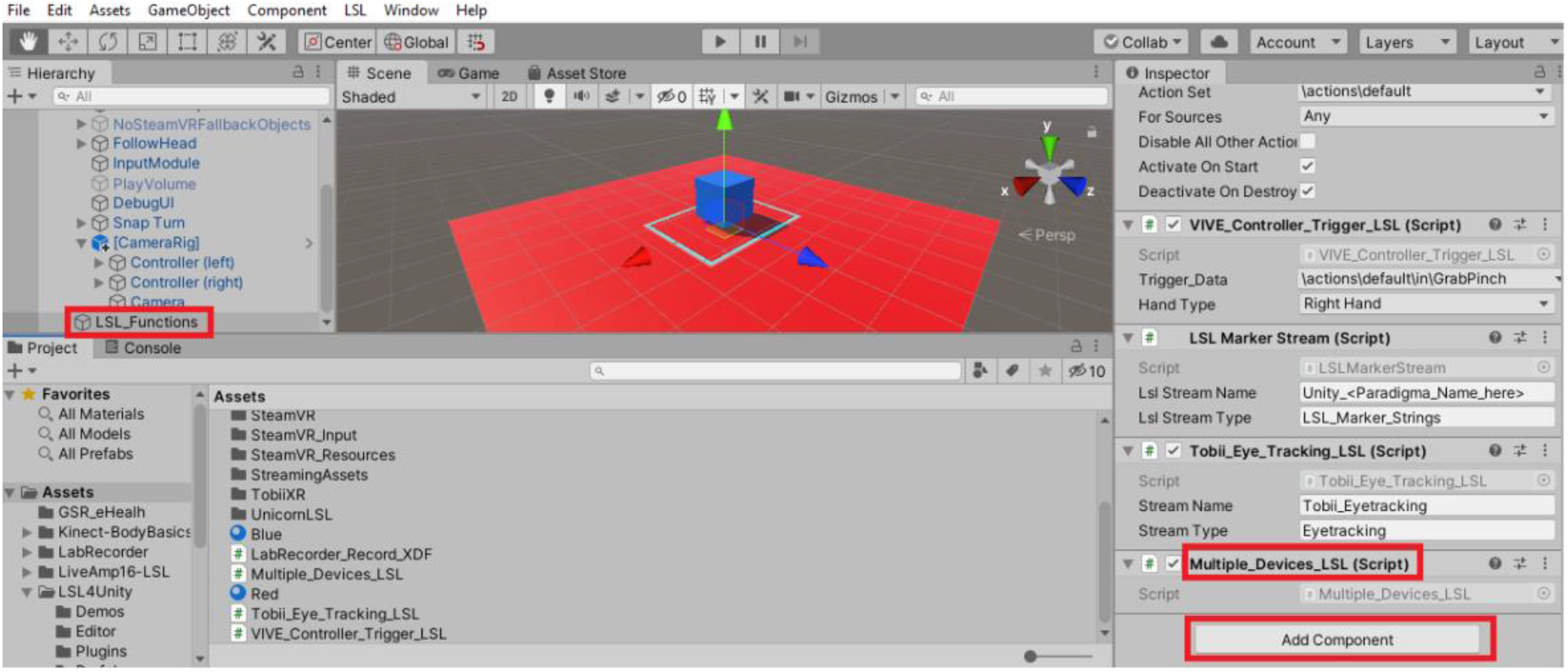
Add the **Multiple_Devices_LSL** script to **LSL_Functions** object

In the following, it is shown how to call **LabRecorder** to start recording the data available on **LSL** automatically by embedding **LabRecorder** in **Unity**.

First, the *LabRecorder* folder available on our GitHub (https://github.com/moeinrazavi/VR_LSL/tree/master/Assets/LabRecorder) is added to the *Assets* folder (Figure 54). Also, the script **LabRecorder_Record_XDF** that we developed in C# should be added to the *Assets* folder (Figure 55). Then the **LabRecorder_Record_XDF** script must be added to the **LSL_Functions** object (Figure 56). By doing this, when the Play button is pressed in **Unity**, all the data sent to **LSL** will start being recorded automatically.

**Figure 54.**
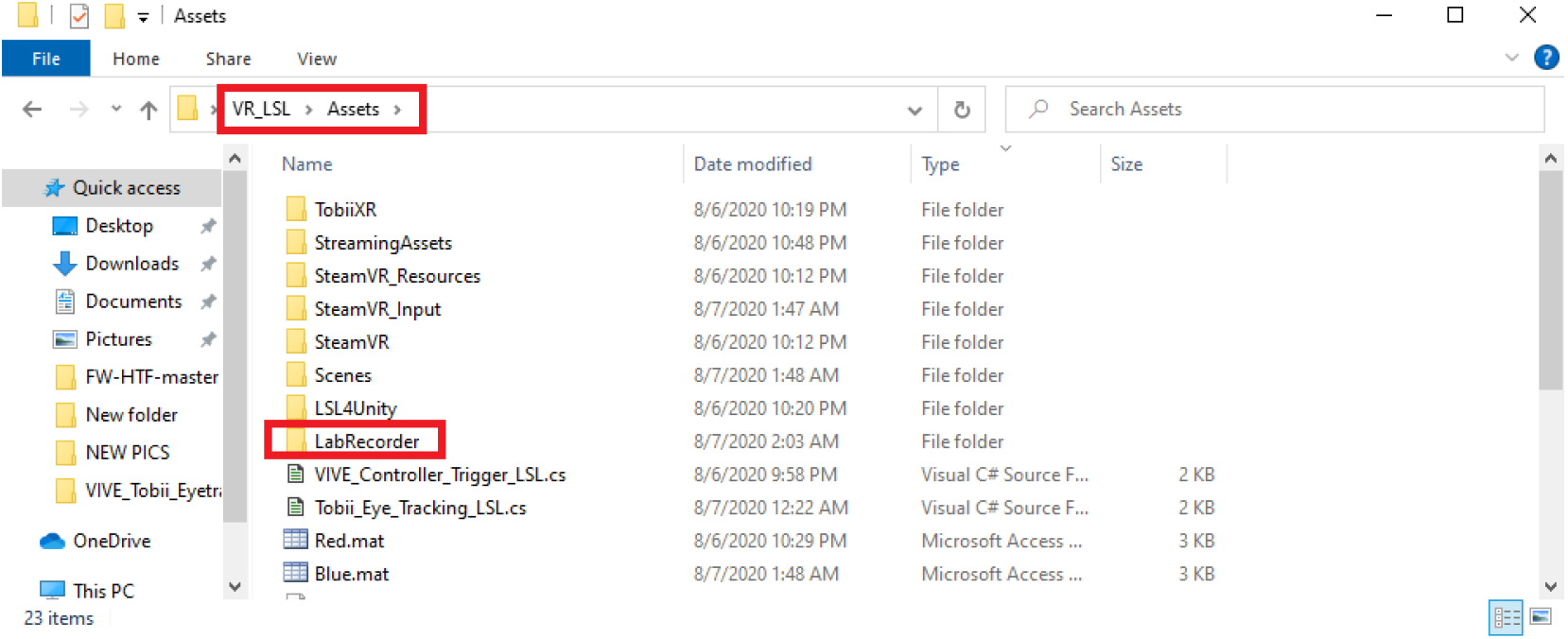
Add the **LabRecorder** file to *Assets* folder

**Figure 55.**
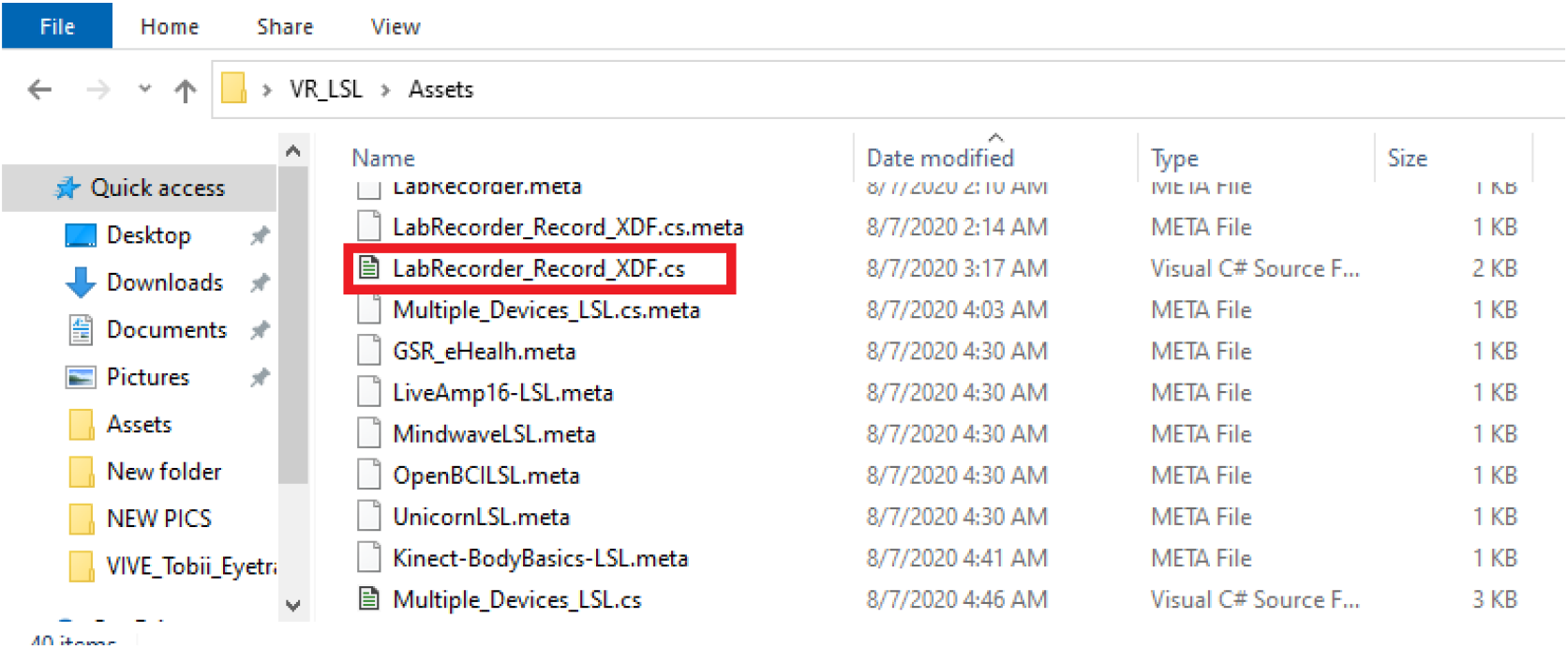
Add the LabRecorder_Record_XDF script to *Assets* folder

**Figure 56.**
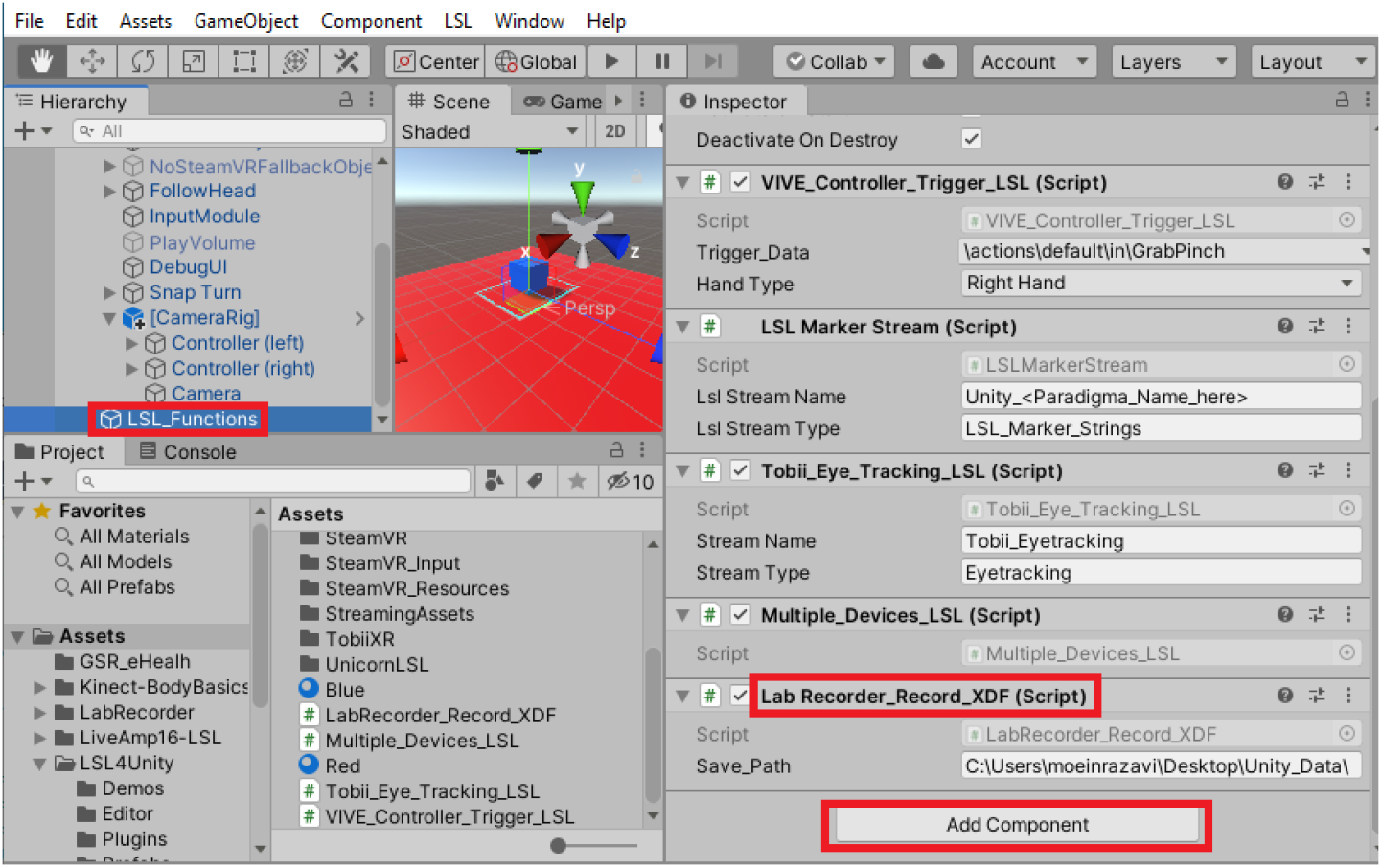
Add the LabRecorder_Record_XDF script to LSL_Functions object

## 6. Results and Discussion

### 6.1. Results

After the experiment is finished, for both **PsychoPy** and **Unity** projects, all the streams will be saved with their associated timestamps in a single .*xdf* file. Each stream can be easily accessed with the assigned **name, type** and **source_id** inside the Python script, MATLAB script or **EEGLAB** toolbox. Ojeda et al. (2014) created an open-source **EEGLAB** toolbox, **MoBILab**, for analyzing data from multiple sensors at the same time (EEG, body motion, eye movement, etc.) [25].

In order to open .*xdf* file using Python, see Appendix.

### 6.2. Discussion and Future Work

The ability to use multiple measures that are synchronized together to distinguish factors affecting behavior and brain functionality is getting more attention. Previous works have mostly used one or a couple of measures to study the human mind and behavior. Some studies showed that using various measures (multimodal experiments) can improve the accuracy and the confidence for interpretation of the results. That said, an accurate and easy to use system to integrate and synchronize multiple measures with different sampling rates is often lacking. In this paper, a practical and comprehensive method on integrating and synchronizing multiple different measures together is provided. We developed some applications that make the integration process easier and accessible for different devices. Once the experiment is created and all the streams are defined, it is straightforward for the experimenter to run the experiment for each subject as everything starts being recorded and saved on the disk automatically. It is also very time-saving in preparing the system for the multisensory experiments. An important customizable feature of the proposed system, which is useful for Brain-Computer Interface and Neurofeedback purposes, is that the user can easily define markers for special behaviors of the signals (e.g. abrupt changes in the signals) [26]. It is expected that for future works, adding stimuli from multiple sources that involve different human senses (e.g., tactile, hear, smell, taste, etc.) can result in higher accuracy and new findings. For instance, Marsja et al. (2019) found that changes in bimodal stimuli (both visual and auditory) conveyed a shift in the performance of spatial and verbal short-term memory tasks, while changes in visual or auditory stimuli individually did not lead to a significant shift in the performance of the mentioned tasks [27]. This can be easily achieved by the aid of our proposed system, by sending the markers indicating the onset, offset, and other information related to multiple stimuli in different streams simultaneously. As multimodal behavioral data are interwoven, using methods that enable the fusion of multimodal data would obtain a wide range of new findings in the human brain and behavior research that have never been found before. For this purpose, the state-of-the-art deep learning models are powerful tools that have recently been used for combining and analyzing data from multiple sources together. Gao et al. 2020 conducted a survey study on using deep learning techniques for multimodal data fusion and how they can help find new interpretations of the data [28]. Thus, deep learning models can be beneficial for multimodal data obtained from human studies as well.

## Appendix

### Install PsychoPy and pylsl

On windows, install the standalone version of **PsychoPy** from https://www.psychopy.org/download.html.

### Install pylsl module on PsychoPy

In order to install **pylsl** (version≥1.13) on PsychoPy using pip. To install **pylsl** on **PsychoPy** use the command: “C:\Program Files\PsychoPy3\python.exe” -m pip install pylsl --user in Windows command line.

### Opening .xdf file in Python

In order to open .*xdf* file in Python, first it is required to install **pyxdf** in python using pip in command line: **pip install pyxdf**. The .*py* file in the link: [pyxdf_example](https://github.com/xdf-modules/xdf-python/blob/d642dbf86f17b8dd94cce56ff339dd57e6d3774a/example/example.py), is an example of opening .*xdf* files in Python. It is recommended to use **Spyder** (https://docs.spyder-ide.org/installation.html) as the Python platform to open the .*xdf* files, since the Variable Explorer panel in **Spyder** allows to track the variables. The fields of a .*xdf* file in Python are shown in shown Figure A1.

**Figure A1.**
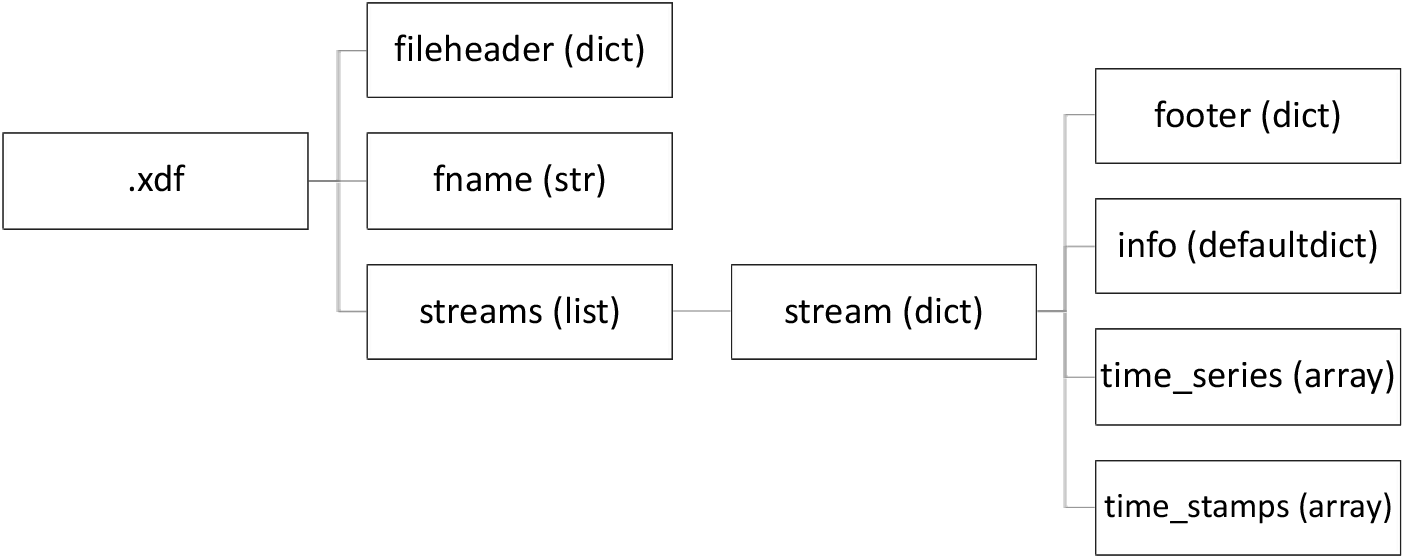
Fields of a .*xdf* file in Python

### Opening .xdf file in MATLAB

In order to open .*xdf* file in MATLAB, first the folder including the *load_xdf.m* function (download from [xdf_importer_GitHub](https://github.com/xdf-modules/xdf-Matlab/blob/87bf5117edfed420ec728c76f50bb71f3722b41c/load_xdf.m)) should be added to MATLAB path (using Set Path in MATLAB Home tab). Then, the .*xdf* file can be loaded in MATLAB workspace by running **load_xdf(“ADDRESS_TO_XDF_FILE.xdf”)** in MATLAB command window. The fields of a .*xdf* file in MATLAB are shown in shown Figure A2.

**Figure A2.**
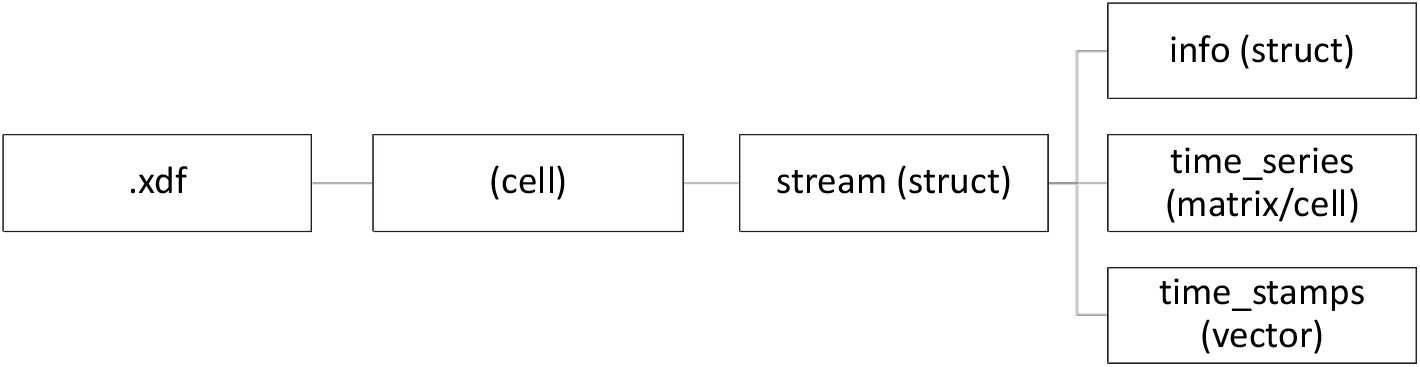
Fields of a .*xdf* file in MATLAB

